# Genomic insights into present local adaptation and future climate change vulnerability of a keystone forest tree species in East Asian

**DOI:** 10.1101/2022.04.04.486908

**Authors:** Yupeng Sang, Zhiqin Long, Xuming Dan, Jiajun Feng, Tingting Shi, Changfu Jia, Xinxin Zhang, Qiang Lai, Guanglei Yang, Hongying Zhang, Xiaoting Xu, Huanhuan Liu, Yuanzhong Jiang, Pär K. Ingvarsson, Jianquan Liu, Kangshan Mao, Jing Wang

## Abstract

Rapid global climate change is posing a huge threat to biodiversity. Assessments of the adaptive capacity for most taxa is usually performed on the species as a whole, but fails to incorporate intraspecific adaptive variation that may play a fundamental role in buffering future shifting climates. Here we generate a chromosome-scale genome assembly for *Populus koreana*, a pioneer and keystone tree species in East Asia temperate forests. We also obtain whole-genome sequences of 230 individuals collected from 24 natural populations. An integration of population genomics and environmental variables was performed to reveal the genomic basis of local adaptation to diverse climate variable. We identify a set of climate-associated single nucleotide polymorphisms (SNPs), insertions-deletions (Indels) and structural variations (SVs), in particular numerous adaptive non-coding variants distributed across the genome of *P. koreana*. We incorporate these variants into an environmental modelling scheme to predict spatiotemporal responses of *P. koreana* to future climate change. Our results highlight the insights that the integration of genomic and climate data can shed on the future evolutionary adaptive capacities of a species to changing environmental conditions.

## Introduction

Climate change is predicted to become a major threat to biodiversity and there is ample evidence of climate-induced local extinctions among plant and animal species ^1^. To escape demographic collapses and extinction, species have to shift their range and migrate to suitable locations, or persist in the same location by genetically adapting to changing environmental conditions from standing genetic variation and *de novo* mutations ^2^. However, migrating in order to keep pace with rapid climate change may be difficult for many organisms, like plants ^3^. Therefore, understanding and predicting the evolutionary potential of a species for future adaptations is not only relevant for understanding whether and how natural species can persist in the context of climate change, but can also benefit conservation and management strategies to cope with global biodiversity loss ^4,5^. The traditional way to assess the capacity for future evolutionary adaptation is via reciprocal transplant experiments or other approaches that involve tracking genetic lineages for many generations ^6^. Doing so is challenging or often unfeasible for many wild non-model organisms due to experimental intractability, long generation times or other challenges to obtain fitness-related phenotypic traits ^7-9^.

Using genomic data to predict the evolutionary potential of populations under climate change provides a different perspective for understanding adaptive evolutionary processes and for assessing the future vulnerability of different populations ^10,11,12^. The first step to evaluate the evolutionary adaptation under changing environmental conditions is to investigate the current spatial patterns of genomic variation, followed by the identification of the genetic basis of local adaptation ^13^. Although there may be millions of variants across the genome within any specific species, relatively few are expected to be related to climate adaptation and hence are relevant for accurate estimates of adaptive capacity. The process of discovering the genomic variants associated with climate adaptation lies at the core of genomic prediction for future climate vulnerability ^14^. Genotype-environmental association approaches are increasingly used to identify loci involved in climate adaptation ^15^. Once candidates for locally adaptive allelic variation have been identified, it is possible to measure genomic vulnerability, which assesses the amount of change in the genetic composition of a population that is required to track future environmental conditions ^10,16^. As such, it goes beyond species-level distribution modelling and provides key insights into assessing the possible maladaptation of populations under future climate change ^4,14^. Therefore, genomic predictions of climate adaptation and maladaptation have immense potential to inform conservation management, especially for threatened species most at risk of local extinction, and/or non-model long-lived species where other experiments are impractical ^17,18^.

Forest trees play a leading role in the global carbon cycle and, along with the characteristics of being the most efficient carbon sink, they will play an increasingly important role in combating climate change and global warming ^9,19^. However, trees are characterized by long lifespans, large body sizes and often have long generation times and large distribution ranges which make them particularly vulnerable to maladaptation under altered climatic scenarios ^20^. With the advance of genomic technologies, it is now possible to characterize genome-wide patterns of genetic diversity even in non-model species ^21,22^. In this context, integrating genomic data into predictive models aimed at quantifying and map spatial patterns of climate maladaptation is especially important for long-lived organisms like trees, for which climate change is likely to happen within the lifetimes of single individuals ^23^.

In the present study, we aim to utilize landscape genomic approaches to investigate the contemporary and future patterns of climate-associated genetic variation for a long-lived poplar species, *Populus koreana*, which is a member of the family *Salicaceae* and is one of the dominant tree species in temperate deciduous forests in East Asia. We present the first *de novo* chromosome-scale reference genome of *P. koreana*, which is then used as a reference for a population genomics study of 230 individuals collected from 24 natural populations across the species’ distribution. We characterize patterns of genome-wide variation, including not only single-nucleotide polymorphisms (SNPs) but also small insertions/deletions (Indels) and larger structural variants (SVs). This variation is further analyzed to decipher genetic diversity, population structure and the demographic history of the species. Finally, we identify candidate loci potentially involved in climate adaptation through genome-wide environmental association studies. By using two different analytical approaches we carry out genomic vulnerability assessment and identify areas where *P. koreana* would be at greater risk due to future climate change.

## Results and Discussion

### Chromosome-scale genome assembly of *P. koreana*

For *de novo* assembly of the *P. koreana* genome, we integrated data from three sequencing and assembly technologies: ∼42.42 Gb of Nanopore long-read sequencing (106×), ∼29.82 Gb of short-read Illumina sequencing (74×), and ∼54. 22Gb of Hi-C paired-end reads (137×) (Supplementary Table 1-4). The final assembly captured 401.4 Mb of genome sequence, with contig N50 of 6.41 Mb and approximately 99.6% (∼399.94 Mb) of the contig sequences anchored to 19 pseudo-chromosomes (Fig. 1a, b; Table 1; Supplementary Table 5), which corresponds to the haploid chromosome number of the species. The high quality of the *P. koreana* assembly was supported by a high mapping rate (99.4%) of Illumina short reads. In addition, we identified 97.8% of the single-copy orthologs from the Benchmarking Universal Single-Copy Orthologs (BUSCO) analysis (Supplementary Table 6), further confirming the continuity and completeness of the assembled *P. koreana* genome.

**Fig. 1.**
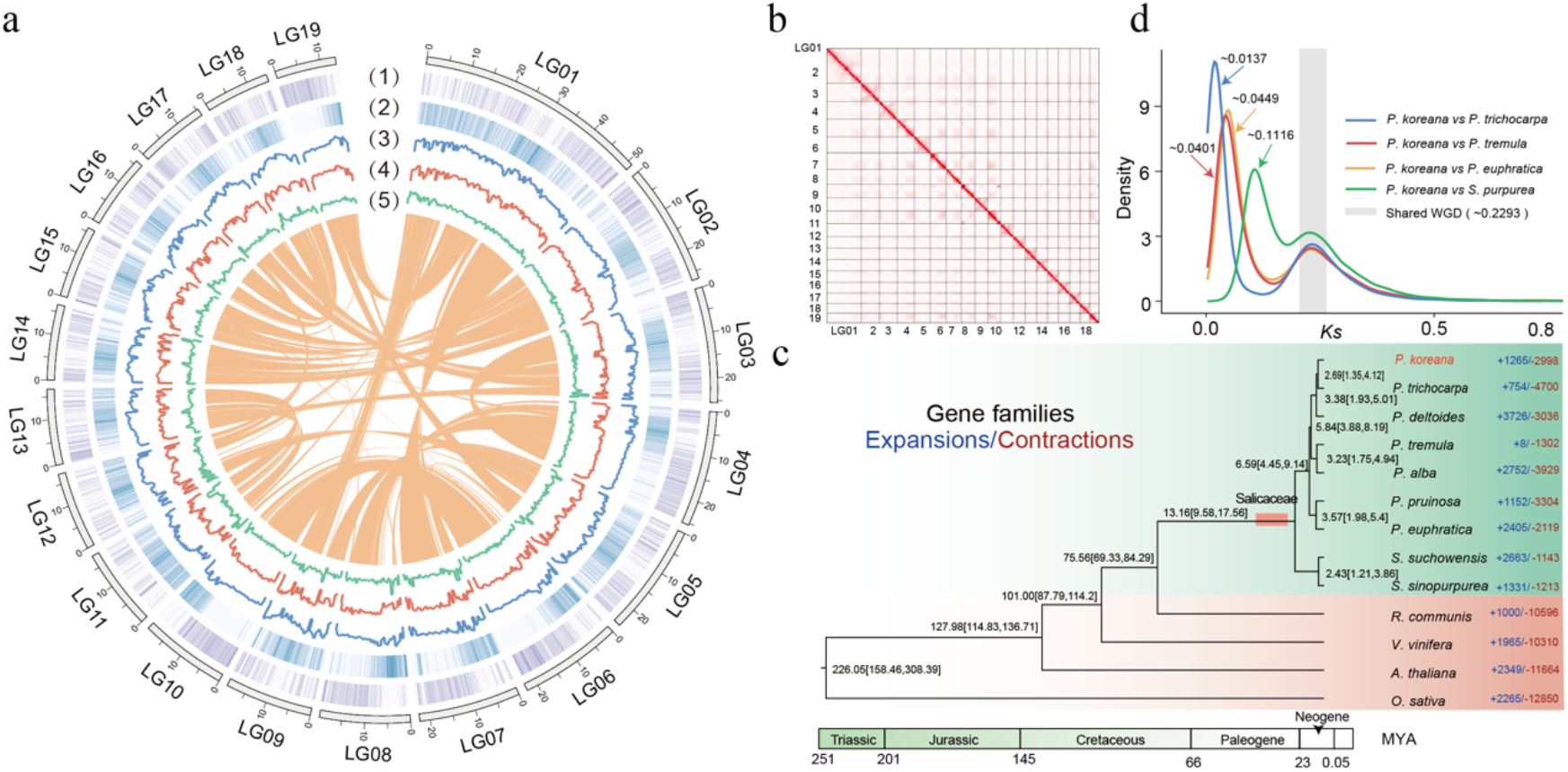
Genome assembly of *Populus koreana* and evolutionary analyses in the *Salicaceae*. **a** Landscape of genomic features and genetic diversity in *P. koreana*. Circles represent, from outermost to innermost, gene density (1), transposable element density (2), the distribution of SNPs (3), Indels (4) and SVs (5) estimated from the population genomic data. Lines in the center represents the intra-genome collinear blocks. **b** Hi-C heatmap showing chromatin interactions at 100 Kb resolution in *P. koreana*. **c** Phylogenetic tree of *P. koreana* and 12 other eudicot species. The number of gene families that expanded (blue) and contracted (red) in each lineage after speciation are indicated beside the tree. The red box indicates the base of the *Salicaceae*. The numbers above nodes in the tree represents divergence times between lineages (million years ago, Mya). **d** Distribution of synonymous substitution rate (*K*s) between syntenic blocks of five species: *P. koreana, P. trichocarpa, P. tremula, P. euphratica* and *S. purpurea*.

**Table 1.**
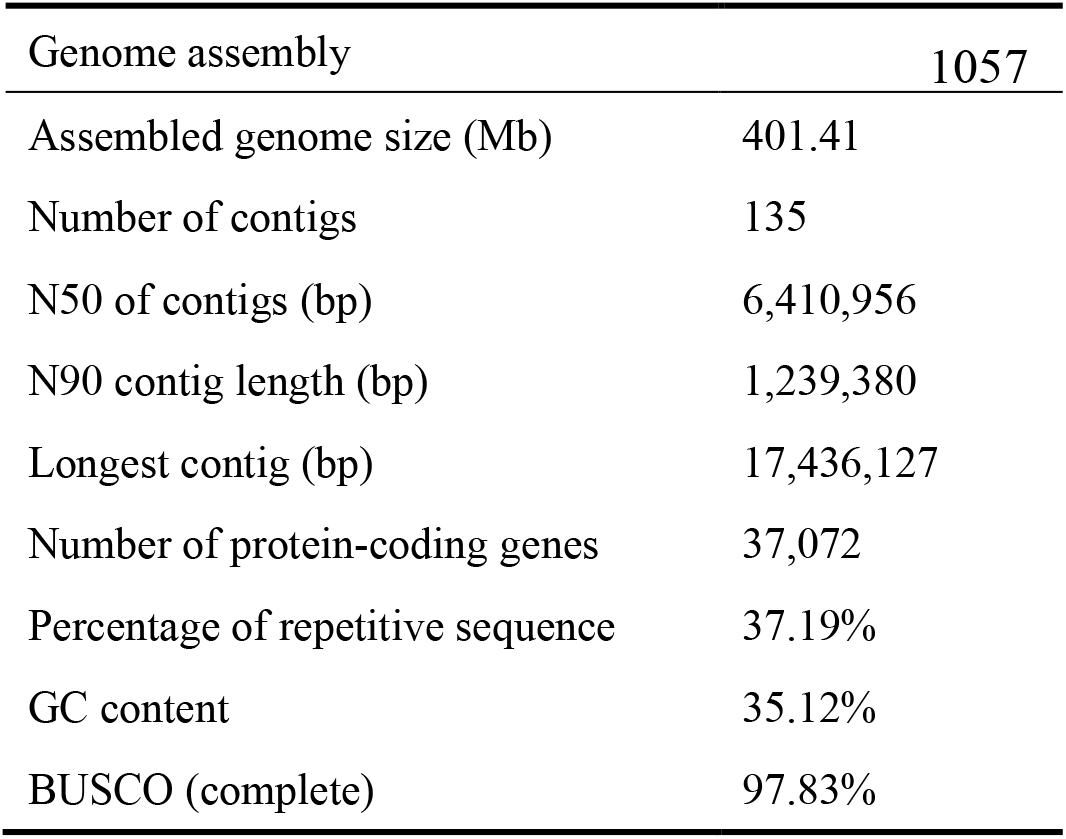
Statistics for the genome assembly and annotation

Repetitive sequences were identified using a combination of homology-based and ab initio approaches. In total, 37.2% of the genome sequences were identified as repetitive elements, including 16.0% of retrotransposons and 17.9% of DNA transposons. Long-terminal repeat (LTR) retrotransposons were found to account for 15.7% of the genome (Supplementary Table 7). After masking the repetitive sequences, we carried out a combination of transcriptome, homology and *ab initio*-based approaches to predict genes. A total of 37,072 protein-coding genes were annotated, with an average coding sequence length of 1,136 bp and an average of five exons per gene (Supplementary Table 8). Of the 37,072 genes, 35,380 (95.4%) could be annotated by at least one public database e. g. Pfam, InterPro, NR, Swiss-Prot, GO and KEGG (Supplementary Table 9). We also identified a set of noncoding RNAs in the *P. koreana* genome (Supplementary Table 10).

To investigate the evolutionary history of *P. koreana*, we performed a gene family clustering using the *P. koreana* genome and 12 other representative angiosperm species, including eight Salicaceae species and four other outgroup species (Fig. 1c). We identified 905 single-copy gene families and used these for phylogenetic tree construction and species divergence time estimation. The phylogenetic analysis showed that *P. koreana* was most closely related to *P. trichocarpa* compared to other selected species in the genus of *Populus*, and the divergence time of the two species was estimated to approximately 2.69 million years ago (Mya). The gene family analysis also revealed 1,265 and 2,998 gene families have undergone significant expansion or contraction in *P. koreana* respectively. Gene Ontology (GO) enrichment analyses showed that the expanded gene families were significantly enriched in stress response, biosynthetic processes, secondary metabolism, and response to external biotic stimulus (adjusted *P* < 0.01) (Supplementary Fig. 1; Supplementary Table 11). Furthermore, investigation of collinear paralogs in the *P. koreana* genome confirmed the occurrence of whole-genome duplication (WGD) (Fig. 1a). By comparing the density distribution of synonymous substitution rates per site (*K*s) of collinear paralogs and orthologs between *P. koreana* and other *Salicaceae* species, the results suggested that all *Salicaceae* species shared the same WGD event before their divergence (Fig. 1d). The shared WGD event was also confirmed by the extensive collinearity between the genomes of *P. koreana* and *P. trichocarpa* (Supplementary Fig. 2) ^24^.

### Population structure, genetic diversity and demographic history

To explore genetic variation in *P. koreana*, we generated whole-genome resequencing data of 230 individuals from 24 populations sampled throughout the natural distribution of the species in Northeast China (Fig. 2a). On average, ∼95% of the clean reads were aligned onto the *P. koreana* genome, with an average depth of 27.4× and coverage of 94.6% (Supplementary Table 12). Using this dataset, we identified a total of 16,619,620 high-quality SNPs and 2,663,202 Indels (shorter than or equal to 50bp). In addition, we also identified a final set of 90,357 large SVs (>50bp).

**Fig. 2.**
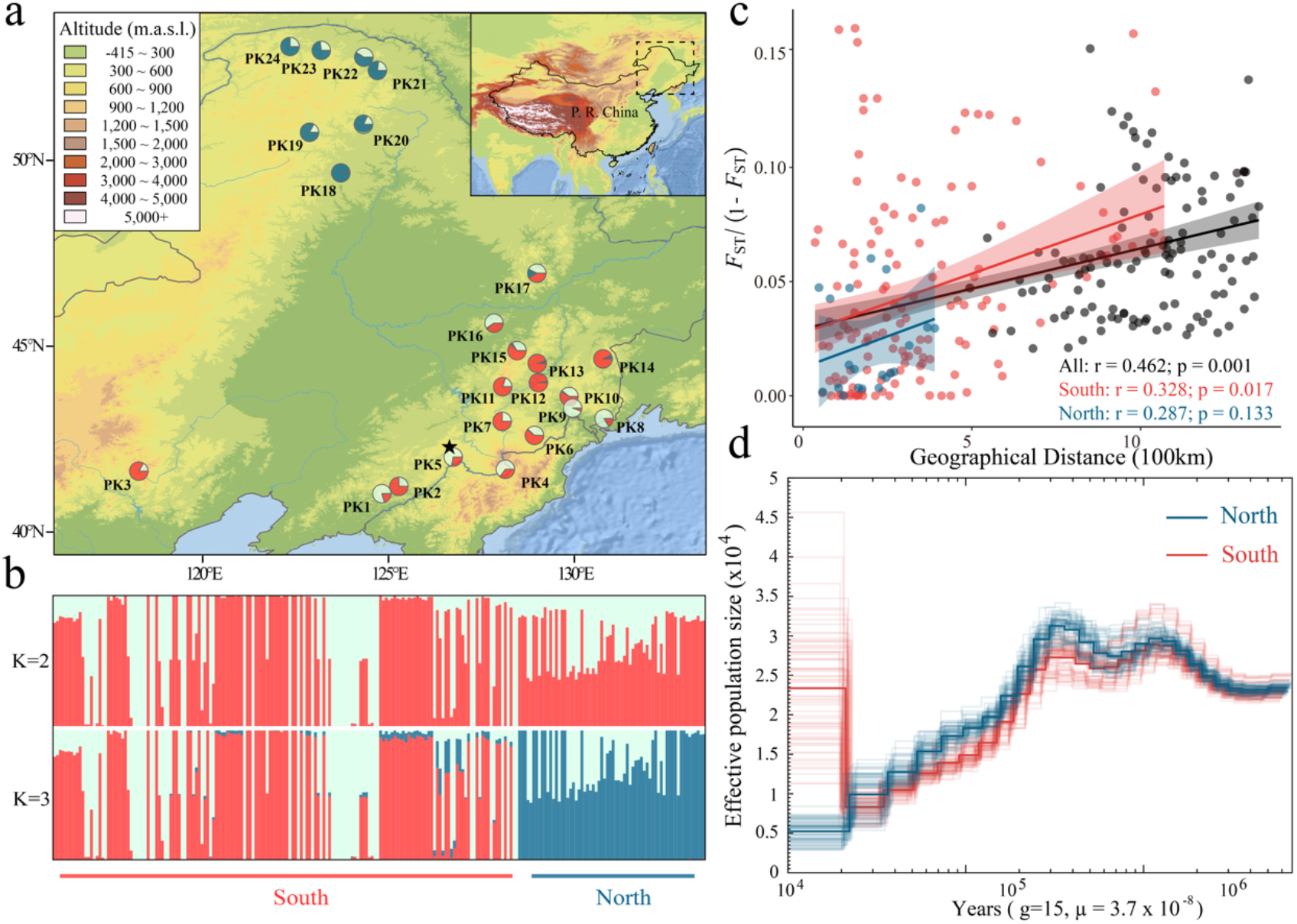
Population genomic analyses of *Populus koreana*. **a** Geographic distribution of 24 natural populations (*circles*) where colors represent ancestral components inferred by ADMIXTURE (according to the substructure at *K* = 3). The location of the individual selected for genome assembly is indicated by a black star. **b** Model-based population assignment using ADMIXTURE for *K* = 2 and 3. The height of each colored segment represents the proportion of the individual’s genome derived from the inferred ancestral lineages. **c** Isolation-by-distance analyses (Mantel’s test) for southern (red dots and line), northern (blue dots and line) and all populations (black dots and line), respectively. **d** Inferred demographic history of southern (blue lines) and northern groups (red lines) of populations from the PSMC model. Bold lines are the median estimates for the seven selected individuals from each of the two groups, whereas faint lines are 140 bootstrap replicates, with 10 replicates being conducted for each of the selected individuals from the two groups.

We first used ADMIXTURE to investigate the genetic structure of the *P. koreana* populations and found that the mode with the number of clusters (*K*) set to 3 exhibited the lowest cross-validation error (Fig. 2a, b; Supplementary Fig. 3), which broadly separated the individuals into two geographical groups (North and South). The North group consists of 66 individuals from seven populations in the Greater Khingan Mountains area while the other 164 individuals from seventeen populations of the Changbai Mountains area, formed the South group. The classification was also supported by a neighbor-joining (NJ) phylogenetic tree which confirmed the two genetic groups (Supplementary Fig. 4). We further examined patterns of genetic differentiation and isolation-by-distance (IBD) between and within each group (Fig. 2c). We detected significant IBD in the southern group and in all populations combined, but not in the northern group, possibly owing to the small number of populations used for the test in the northern group. Moreover, the pattern of IBD was stronger for all populations combined compared to populations in either the southern or northern group alone. It is possible that the allopatric fragmentation into isolated refuges during glacial periods has contributed to the accumulation of genetic differences between the disjunct populations, in particular because no or few distribution records are present in the intermediate areas ^25, 26^. Nevertheless, the genetic differentiation between the two genetic groups was found to be weak (Supplementary Fig. 5, the average *F*_ST_ values: 0.021). The genome-wide screens of genetic variation within and between groups revealed that nucleotide divergence (d_xy_) between the two groups was almost the same as the nucleotide diversity within groups (Supplementary Fig. 6), again suggesting that population structure in *P. koreana* is relatively weak.

To further infer the demographic history of the *P. koreana*, we performed the pairwise sequentially Markovian coalescent (PSMC) to assess change in effective population size (*N*_e_) over the past ∼3-4 million years ago (Mya) (Fig. 2d). We found that different populations of *P. koreana* displayed highly similar demographic trajectories (Supplementary Fig. 7). The inferred *N*_e_ only differed between the southern and northern groups following the last glacial maximum (LGM, 10,000-20,000 years ago), where samples from the northern group showed a steady population decline while a slight population expansion was observed in samples from the southern group. The inferred demographic histories of *P. koreana* populations were also confirmed by the patterns in site frequency spectrum as summarized by Tajima’s D statistics (Supplementary Fig. 8), where Tajima’s D was on average positive in the northern group populations while the average Tajima’s D was slight negative in the southern group.

We estimated nucleotide diversity (π) in 10 Kbp non-overlapping windows across the genome for the 24 populations and found qualitatively similar results, with an average diversity of 1.08% (Supplementary Fig. 9). In addition, the genome-wide decay of linkage disequilibrium (LD) as a function of physical distance showed similar patterns in the southern and northern populations, with *r*^2^ declining below 0.2 after ∼15 Kbp on average (Supplementary Fig. 10). Overall, our results reveal weak population structure in *P. koreana* between southern and northern population groups which might have been geographically isolated following the LGM.

### Identifying genomic variants associated with local climate adaptation

The high-quality reference genome for *P. koreana* coupled with the high-depth resequencing data generated in this study facilitate the precise characterization of genomic information, including not only SNPs, but also Indels and SVs that are usually ignored ^27^. To investigate the extent to which genetic variation is driven by contemporary climate gradients and to detect the environment-associated genetic variants, we used two complementary genotype--environment association (GEA) approaches. First, we tested for GEAs for 19 environmental variables (10 temperature and 9 precipitation-related variables, Supplementary Table 13) using latent fixed mixed modeling (LFMM) ^28^, which tests for associations between genotypes and environment variable while accounting for background population structure. With a q-value cut-off of 0.05, we identified a total of 3,013 SNPs, 378 Indels, and 44 SVs (Supplementary Fig. 11), involving 514 genes that were significantly associated with one or more environmental variables (Fig. 3; Supplementary Fig. 12; Supplementary Table 14). In general, we found that these environment-associated variants were widely distributed across the genome of P. *koreana* and did not cluster in specific regions.

**Fig. 3.**
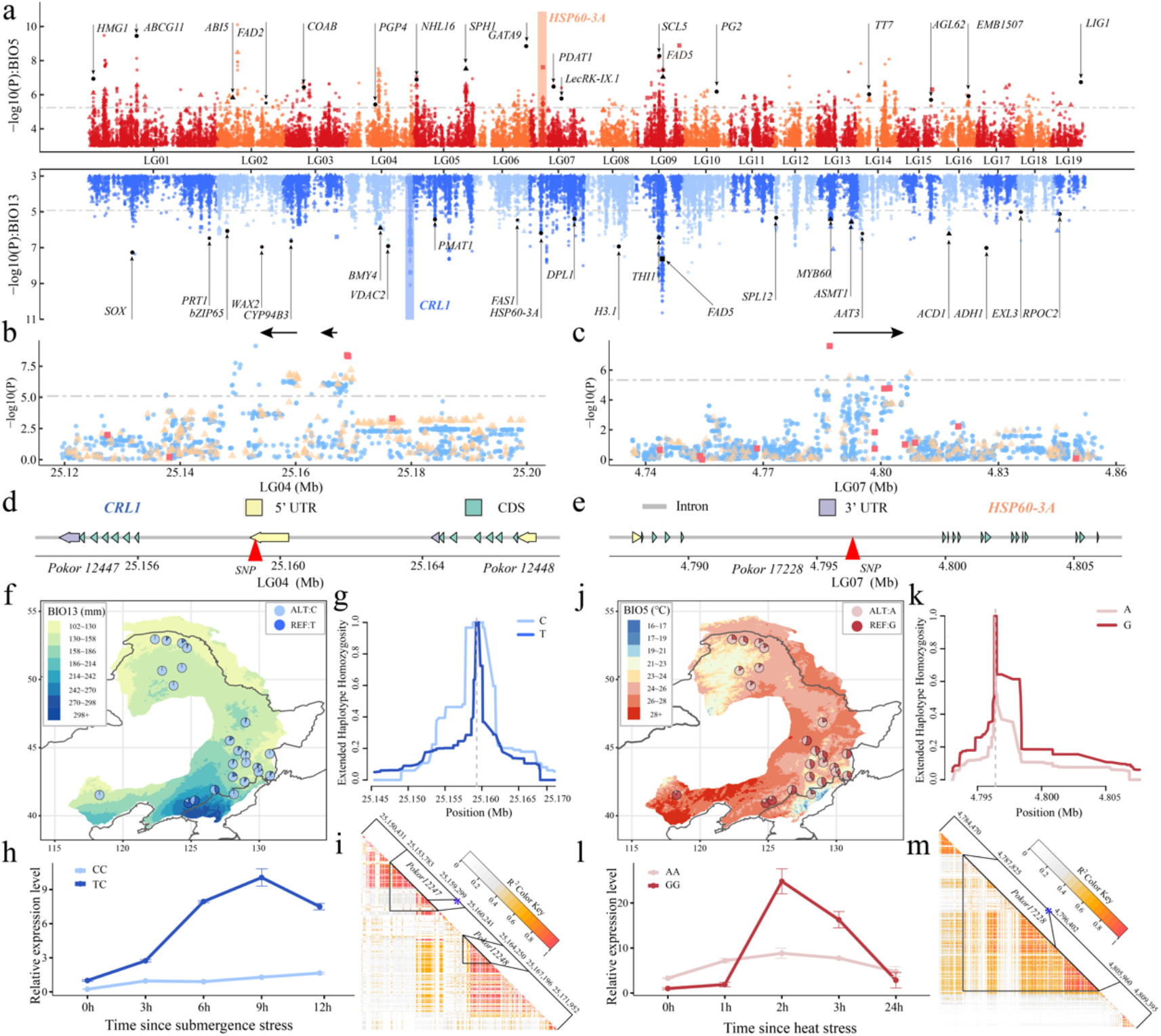
Genome-wide screening of the loci associated with local environmental adaptation. **a** Manhattan plots for variants associated with the Maximum Temperature of Warmest Month (BIO5) (red, upper panels) and the Precipitation of Wettest Month (BIO13) (blue, lower panels). Dashed horizontal lines represent significance thresholds. Different chromosomes are distinguished by different shades of the major color. Selected candidate genes are labeled in the plot at their respective genomic positions. **b**,**c** Local manhattan plots around two candidate genes (black arrows), *CRL1* (*Pokor12447* and *Pokor12448*) *and HSP60-3A* (*Pokor17228*) on chromosome 4 and 7, associated with BIO5 and BIO13 respectively. SNPs, Indels and SVs are represented by blue dots, yellow triangles and red squares, separately. **d**,**e** The gene structure of selected genes, with the two representative candidate SNPs corresponding to the sites shown in **f-m** are marked by red triangles, respectively. **f**,**j** Allele frequencies of the candidate SNPs associated with BIO5 (**f** LG04:25159299) or BIO13 (**j** LG07: 4796402). Colors on the map are based on variation in the relevant climate variables across the distribution range. **g**,**k** Decay of extended haplotype homozygosity (EHH) for the two alternative alleles at the two representative SNPs. **h**,**l** Comparison of the relative expression of *CRL1* (h) and *HSP60-3A* (l) genes between the two genotypes using qRT-PCR after submergence (h) and heat (l) treatment, respectively. **i**,**m** Heatmap of LD surrounding the two candidate regions show above. The blue stars indicate the two representative SNPs, and the black triangles mark the corresponding genic regions.

LFMM is a univariate approach that tests for associations between one variant and one environmental variable at a time and to alleviate these issues we also used a complementary multivariate landscape genomic method, redundancy analysis (RDA) ^29^, to identify covarying variants that are likely associated with multivariate environment predictors. To avoid issues due to multicollinearity, six uncorrelated environmental variables (Spearman’s r <0.6, Supplementary Fig. 13) were selected for the RDA analyses, including three temperature variables (Annual Mean Temperature (BIO1), Isothermality (BIO3), Maximum Temperature of Warmest Month (BIO5)) and three precipitation variables (Precipitation of Wettest Month (BIO13), Precipitation Seasonality (BIO15), Precipitation of Coldest Quarter (BIO19)). Of the 3,435 significant variants identified in our LFMM analyses, 1,779 (1,554 SNPs, 206 Indels and 19 SVs) were found to display extreme loadings (standard deviation >3) along one or multiple RDA axes (details in Materials and Methods). These shared variants were regarded as “core adaptive variants” for local climate adaptation and they were broadly distributed across the genome (Supplementary Fig. 14). Significantly stronger genetic differentiation (*F*_ST_) were observed at these adaptive variants (Supplementary Fig. 15), indicating that spatially varying selection has likely driven population differentiation at climate-associated adaptive variants compared to random neutral genetic markers ^30,31^. On average, we found that more adaptive variants were associated with precipitation-related compared to temperature-related variables (Supplementary Fig. 14).

Of the core adaptive variants, only 3.2% were non-synonymous and 2.0% were synonymous mutations, with all remaining variants being non-coding (Supplementary Table 15), indicating that adaptation to climate in *P. koreana* have primarily evolved as a result of selection acting on regulatory rather than on protein-coding changes ^32^. In particularly, we found a significant enrichment of climate adaptive variants located in the 5’ UTR of genes (Supplementary Fig. 16). Moreover, 9.7% of the adaptive variants were found to be located within the regions of accessible chromatin as identified by transposase-accessible chromatin sequencing (ATAC-seq) (Supplementary Table 14), again suggesting that changes in cis-regulatory elements may play important roles in driving environmental adaptation in natural populations of *P. koreana*. To further assess the selection pressures acting on the climate adaptive variants, we calculated the standardized integrated haplotype score (iHS) across all common variants to identify loci with signatures of selective sweeps ^33^. Our results show that climate-associated variants did not display stronger signatures of positive selection compared to randomly selected SNPs (Supplementary Fig. 17), suggesting that adaptation to local climate in *P. koreana* may largely arise by polygenic selection, characterized by subtle to moderate shifts in allele frequencies of many loci with small effect sizes ^34,35^.

Together, we identified many well-studied genes involved in climate adaptation in *P. koreana* (Supplementary Fig. 12; Supplementary Table 14 and 16), although no significant functional enrichment could be detected. For loci that are significantly involved in adaptation to precipitation-associated environmental variables, the distribution of allele frequencies in general showed similar patterns (Supplementary Fig. 18). A prime example of such a locus that is strongly associated with variation in precipitation during the wettest month is *CRL1* (Fig. 3a). It is a LOB-domain transcription factor that play an essential role in crown root formation and that has been shown to play a critical role in regulating root system architecture in response to flooding and drought stresses ^36,37^. We found two tandem duplicates homologous to *Arabidopsis CRL1* in *P. koreana* (Fig. 3b), and we identified a total of 104 candidate adaptive variants (83 SNPs, 19 Indels and 2 SVs) located around these two genes (*Pokor12247, Pokor12248*). We choose one candidate adaptive SNP located in 5’ UTR of *Pokor12247* (LG04:25159299) as an example to show the distribution pattern of allele frequencies (Fig. 3d). The T allele was mainly distributed in the southeast regions of the *P. koreana* distribution range that are characterized by heavy precipitation in the wettest month, whereas the C allele was almost fixed in areas experiencing low rainfall (Fig. 3f). To verify the potential function of *Pokor12247* in mediating adaptation to extreme precipitation, we performed qRT-PCR to profile its expression under submergence stress. Interestingly, we found that *Pokor12247* exhibited differential expression between genotypes in response to submergence stress treatment, with individuals carrying the TC genotype at LG04:25159299 displaying enhanced expression compared to individuals with the CC genotypes in response to submergence (Fig. 3h). This indicates that the haplotype carrying the T allele may be associated with increased tolerance to submergence in regions with high rainfall. Nevertheless, the relatively high degree of LD (Fig. 3i) at this region makes it hard to identify the true causal variant(s) that are involved in mediating environmental adaptation. Furthermore, we did not observe signals of strong recent selection at this locus ^38^. The extended haplotype homozygosity (EHH) did not exhibit significant differences between haplotypes carrying the T or the C allele at the focal SNP (Fig. 3g; the standardized |iHS| score =1.693), which again supports a polygenic pattern of adaptation ^39^. In addition, many other genes were also found to be involved in precipitation-associated adaptation (Supplementary Fig. 12, 18; Supplementary Table 14, 16), such as *Pokor27800*, which encodes a MYB transcription factor (orthologous to *MYB60*) that is essential for promoting stomata opening and closure in response to flooding and/or drought stresses ^40^; *Pokor18547* is orthologous to *Arabidopsis DPL1* and encodes a sphingoid long-chain base-1-phosphate lyase, and this gene has been shown to be involved in the dehydration stress response ^41^; Similarly, *Pokor25841*, encoding a SQUAMOSA promoter binding protein-like transcription factor orthologous to *Arabidopsis SPL12*, has been shown to be an important regulator of plant growth, development and stress responses ^42^.

We also identified a set of temperature-associated loci, including genes orthologous to *Arabidopsis HMG1, PGP4, FAD5, EMB1507* showing similar allele frequency distribution patterns as we saw for the precipitation associated genes (Fig. 3a; Supplementary Fig. 12, 19; Supplementary Table S14). A striking example of such a locus associated with variation in the maximum temperature of the warmest month was *Pokor17228*, which encodes a heat shock protein (HSP) orthologous to *Arabidopsis HSP60-3A* ^43^. The rapid synthesis of HSPs induced by the heat stress can protect cells from heat damage and enable plants to obtain thermotolerance by stabilizing and helping refold heat-inactivated proteins ^44^. Relatively high LD was found within the region surrounding this gene (Fig. 3m), including a total of 62 candidate adaptive variants (59 SNPs, 2 Indels and 1 SV). We chose one candidate adaptive SNP located in an intronic region of *Pokor17228* (LG07: 4796402) for further exploration of allele frequency distribution patterns (Fig. 3e). Populations located in areas with relatively higher temperature of the warmest month of the year were more likely to carry the G allele, while the A allele was more likely to be observed in regions with low temperatures (Fig. 3j). To further explore the role of *Pokor17228* in the response to heat stress, we examined the expression pattern of the two genotypes (GG vs AA) at the candidate SNP. The genotypes with the candidate warm-adapted allele (G) showed much higher expression than the A allele after two and three hours of heat stress treatment (Fig. 3l), indicating that *Pokor17228* is a likely candidate gene for heat stress tolerance in *P. koreana*. Similar to what is observed at most candidate adapted variants, we failed to detect signatures of strong recent selection signal at this locus (the standardized |iHS| score =1.661). Despite this, the haplotypes carrying the warm-adapted allele (G) had elevated EHH relative to the haplotypes carrying the other allele (A) (Fig. 3k), suggesting it might have experienced weak positive selection.

Taken together, our results support a polygenic model for local climate adaptation across natural populations of *P. koreana*. The thorough characterization of the genetic basis underlying ecological adaptation performed in this study offers promising information for predicting species response to future climate change ^12,14^.

### Genomic vulnerability prediction to future climate change

Based on the established contemporary genotype–environment relationships and the identified climate-associated genetic loci, we aim to make predictions of how populations of *P. koreana* will response to future climate change. To achieve this we used two complementary approaches to investigate the spatial pattern of maladaptation across the range of *P. koreana* and to identify populations that are most vulnerable to future climate shifts under four CMIP6 emission scenarios of shared socioeconomic pathway (SSP126, SSP245, SSP370 and SSP585) for two defined periods (2061-2080 and 2081-2100) ^45^. First, we calculated the risk of nonadaptedness (RONA) for each population based on the 19 environmental variables (Fig. 4; Supplementary Fig. 20). RONA measures the expected allele frequency shifts required to cope with future climate conditions after establishing a linear relationship between allele frequencies at environmentally associated variants and present climates ^16,46^. As expected, for most environmental variables, RONA increases under more severe climate change scenarios, with higher emissions leading to increased overall RONA values (i.e. SSP585 vs. SSP126, more details in Supplementary Table 17). Moreover, we found substantial variation in RONA estimates among different environmental variables, and for each variable, RONA values were also different across populations (Fig. 4; Supplementary Fig. 20; Supplementary Table 17). We choose predictions for two environmental variables (BIO5 and BIO13, described above) under future climate scenario SSP370 in 2061-2080 as representative outcome. Populations located in areas with more drastic environmental changes are anticipated to have greater RONA values. RONA estimates for temperature variables were substantially higher than those projected from precipitation-induced responses, indicating that substantial allele frequency shifts are needed at temperature-associated loci to cope with future temperature increases (Fig. 4 a,b; Supplementary Fig. 21) ^16^. In addition, we found that populations in both the northern and southern distributions of *P. koreana* had almost equally large values of RONA in face of temperature changes. In contrast, for precipitation changes southern populations displayed much higher genomic vulnerability compared to northern populations where RONA values were generally low, in particular for those populations near the Korean Peninsula that were predicted to experience severe rainfall and extreme precipitation events in the future (Fig. 4 c,d).

**Fig. 4.**
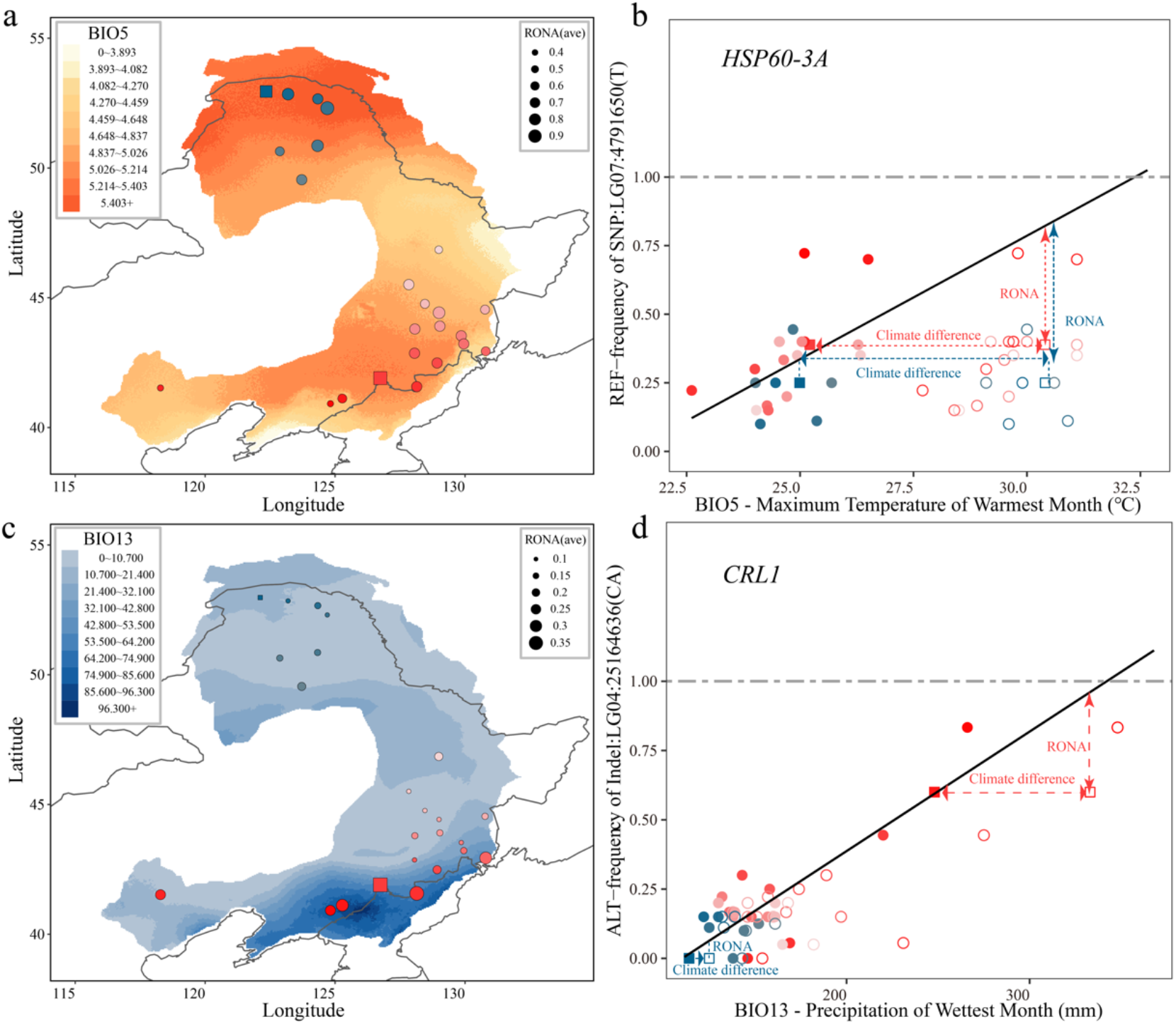
Risk of non-adaptedness (RONA) of *P. koreana* to future climatic conditions. **a**,**c** RONA estimates for two environmental variables (a: BIO5; c: BIO13) for populations under the climate scenarios of SSP370 in 2060-2080. The raster colors on the map represent the degree of projected future climate change (absolute change). Areas with darker red (a) or blue (c) are predicted to experience more dramatic change in the respective climate variables. Solid circles with different colors on the map reflect different natural populations, where red and blue represents the southern and northern groups of populations, respectively. Circle size represent average RONA values in the populations and squares (one southern and one northern) indicate the two example populations illustrated in b and d. **b**,**d** Example diagrams of RONA to future climatic conditions, presented on genotype-environment association plot, for two climatic-associated variants within *HSP60-3A* (b) and *CRL1* (d), respectively. Hollow circles represent future climate conditions for the populations and provide the basis for calculating the required allele frequency change (RONA) to track future climatic conditions. The two example populations in (a) and (c) are again highlighted by squares.

Second, we used the gradient forest (GF) approach to model the turnover in allele frequencies along present environmental gradients and predict genetic offset to a projected future climate ^10^. We first performed GF analyses to determine the relative importance of various environmental variables based on the putatively environmental-associated variants. Of the 19 environmental variables tested, the top explanatory variables were mostly precipitation related, again suggesting that adaptation to precipitation is likely the most important environmental driver shaping the spatial patterns of adaptive genetic variation (Fig. 5b). To avoid multicollinearity issues and to simultaneously consider the ranked importance by GF, we used the same six uncorrelated environmental variables that were used in the RDA analyses (BIO15, BIO19, BIO13, BIO1, BIO3, BIO5) to estimate genomic vulnerability across the geographic distribution of *P. koreana*. By visualizing climate-associated genetic variation across the natural distribution of *P. koreana*, we found that adaptive genetic variation could be largely explained by these six climatic variables (Supplementary Fig. 22). Moreover, we observed that the use of the six uncorrelated climatic variables or all the nineteen climatic variables had no major impact on the results (Supplementary Fig. 23, 24). Overall, genomic offset was found to be highest in southeastern populations near the Korean Peninsula (Fig. 5a), where also high RONA values for both precipitation and temperature-related variables were observed (Fig. 4). Therefore, all these findings demonstrate that southeastern populations of *P. koreana* near the Korean Peninsula are expected to experience higher magnitudes of environmental change in the future, from both warmer temperatures and more extreme summer rainfall conditions, and are therefore likely to be more vulnerable to climate change ^14,17^.

**Fig. 5.**
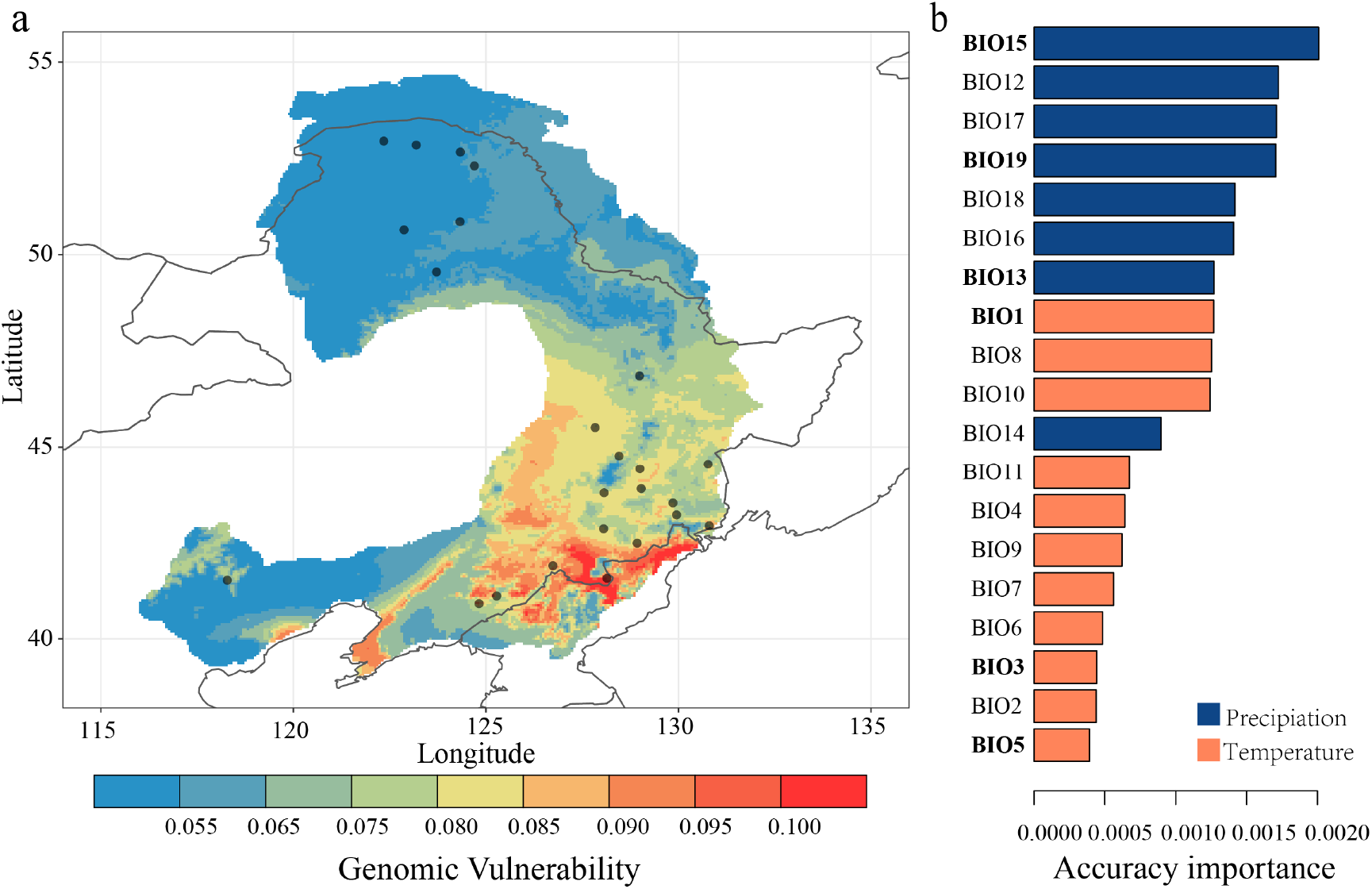
Gradient-forest modelling and predicted genomic vulnerability. **a** Map of genetic offset across the natural distribution of *P. koreana* for the period 2060-2080 under the scenario SSP370. The color scale from blue to red refers to increasing genetic offset and points on map reflect sampled populations. **b** Ranked importance of 19 environmental variables based on the gradient forest analysis shows that precipitation-related environmental factors strongly explain spatial genomic variation in *P. koreana*. The six uncorrelated environmental variables selected for calculation of genetic offset are highlighted in bold text.

Although genomic information shows great promise for predicting future vulnerability of species to climate change, recent simulation studies revealed that the measures of potential genomic offset could be artificially inflated by other neutral processes such as population structure and effective populations sizes ^47^. However, in our study, both RONA and genomic offset estimated here are all based on candidate climatic adaptive variants that were identified by genome scan procedures after accounting for the effects of neutral population structure. In addition, compared to the expectation that populations with small *N*_e_ would exhibit greater signatures of genetic drift that further leads to greater turnover of allele frequencies and cause false-positive signals of increased estimates of offsets ^47^, we did not find a relationship between the level of nucleotide diversity, which is proportion to *N*_e_, and the estimated genomic offsets across populations (Supplementary Fig. 25a). Furthermore, as higher genetic drift in small populations would limit the efficacy of purifying selection and result in higher genetic load ^48^, we further estimated and compared genetic load using a measure that compare the proportion of 0-fold nonsynonymous to 4-fold synonymous SNPs among populations. In line with the results on nucleotide diversity, there was no relationship between the estimated genetic load and offsets (Supplementary Fig. 25b). Together, all results suggest that neutral evolutionary processes should not have much impact on our estimates of genetic offsets and the vulnerability assessment across populations to future climate change.

The metrics of genomic vulnerability estimated here are therefore reliable and have clear implications for not only delineating future conservation units but also informing management decisions of this key long-lived tree species ^49^. For instance, the southeastern populations nearby the Korean Peninsula are inferred to be most at risk from future higher temperatures and more intense precipitation. Considering that these populations contain many unique, climate-adaptive germplasms where a set of adaptive alleles for warmer and wetter climates have been identified in multiple functional important genes, *ex situ* conservation efforts may be appropriate and necessary in this area ^50^.

## Conclusion

Ongoing climate change is predicted to threaten populations for numerous species, and despite the importance of intraspecific adaptive variation in determining responses, predictions of vulnerability to climate change usually lack a component of evolutionary responses. In this study, we first assembled a highly continuous, accurate, and complete genome of *P. koreana* using Nanopore long reads and Hi-C interaction maps. The high-quality reference genome enables us to perform comprehensive population genomic analyses, which are fundamental for an accurate characterization of the spatial patterns of genomic variation and for gaining unique insights into the genetic architecture of climatic adaptation. We further combine genomics, space-for-time and machine-learning approaches to predict broad spatiotemporal responses to future climate change in this species. Most notably, we identify a set of populations located in southeastern part of the current distribution range as being potentially most vulnerable under future climate scenarios, information which is invaluable for developing conservation and management strategies. To summarize, our results demonstrate how genomic data can be used to assess climate change vulnerability in an ecologically important non-model species, showing great promise as the first step in the design of applied conservation efforts in response to a rapidly changing climate.

## Materials and Methods

### Plant materials and genome sequencing

Fresh leaf tissues were sampled from a wild *P. koreana* plant growing in Changbai Mountain of Jilin province in China, and immediately stored in liquid nitrogen. Total genomic DNA was extracted using the CTAB method. For the Illumina short-read sequencing, paired-end libraries with insert sizes of 350bp were constructed and sequenced using an Illumina HiSeq X Ten platform. For the long-read sequencing, the genomic libraries with 20 Kbp insertions were constructed and sequenced utilizing the PromethION platform of Oxford Nanopore technologies. For the Hi-C experiment, about 3g of fresh young leaves of the same *P. koreana* accession was ground to powder in liquid nitrogen. A sequencing library was then constructed by chromatin extraction and digestion, DNA ligation, purification and fragmentation ^51^, and was subsequent sequenced on an Illumina HiSeq X Ten platform.

### Genome assembly and scaffolding

The quality-controlled reads were firstly corrected via a self-align method using the NextCorrect module in the software NextDenovo v2.0-beta.1 (https://github.com/Nextomics/NextDenovo) with parameters “reads_cutoff=1k, seed_cutoff=32k”. Smartdenovo v1.0.0 (https://github.com/ruanjue/smartdenovo) was then used to assemble the draft genome with the options -k 21 -J 3000 -t 16. To improve the accuracy of the draft assembly, two-step polishing strategies were applied: the first step included three rounds of polishing by Racon v1.3.1 ^52^ based on the corrected ONT long reads. The second step includes four rounds of polishing by Nextpolish v1.0.5 ^53^ based on cleaned Illumina short reads after removing adapters and low-quality reads using fastp v0.20.0 ^54^ with parameters ‘-f 5 -F 5 -t 5 -T 5 -n 0 -q 20 -u 20’. Finally, allelic haplotigs were removed using the purge_haplotigs v1.1.1 ^55^ software with the options ‘-l 5 -m66 -h 170’ to obtain the final contig-level assembly.

For chromosome-level scaffolding, the Hi-C reads were first filtered by fastp v0.20.0 with parameters described above. Each pair of the clean reads were then aligned onto the contig-level assembly by bowtie2 v2.3.2 ^56^ with parameters ‘-end-to-end, - very-sensitive -L 30’. The quality of Hi-C data was evaluated by HiC-Pro v2.11.4 ^57^, which further classified read-pairs as valid or invalid interaction pairs. Only valid interaction pairs were retained for further analysis. Finally, scaffolds were clustered, ordered and oriented onto chromosomes using LACHESIS ^58^ with parameters: CLUSTER MIN RE SITES = 100; CLUSTER MAX LINK DENSITY=2.5; CLUSTER NONINFORMATIVE RATIO = 1.4; ORDER MIN N RES IN TRUNK=60; ORDER MIN N RES IN SHREDS=60. The placement and orientation errors that exhibit obvious discrete chromosome interaction patterns were then manually adjusted.

The completeness of the genome assembly was assessed by both the representation of Illumina whole-genome sequencing short reads from mapping back reads to the assembly using bwa v0.7.12 ^59^, and by Benchmarking Universal Single-Copy Orthologs (BUSCO) v4.0.5 ^60^ with the searching database of “embryophyte_odb10”.

### Repeat and gene annotation

For repeat annotation, we used the Extensive de-novo TE Annotator (EDTA v1.9.3) ^61^, which incorporates well performed structure- and homology-based programs (including LTRharvest, LTR_FINDER, LTR_retriever, TIR-learner, HelitronScanner and RepeatModeler) and subsequent filtering scripts, for a comprehensive repeat detection. Subsequently, TEsorter (v1.2.5, https://github.com/zhangrengang/TEsorter/) ^62^ was used to reclassify those TEs that were annotated as “LTR/unknown” by EDTA.

For gene annotation, we first used RepeatMasker v4.1.0 ^63^ to mask the whole genome sequences with the TE library constructed using EDTA. An integrated strategy that combined homology-based prediction, transcriptome-based prediction and *ab initio* prediction was used to predict the protein-coding genes. For homology-based gene prediction, published protein sequences of six plant species, including *Populus euphratica, Salix brachista, Salix purpurea, Populus trichocarpa, Arabidopsis thaliana* and *Vitis vinifera* were downloaded and aligned onto the repat-masked genome by using TBLASTN (ncbi-BLAST v2.2.28 ^64^) program with E-value cutoff setting 1e^-5^, and GeneWise v2.4.1 ^65^ was then used to predict gene models with default settings. For transcriptome-based gene prediction, trimmed RNA-seq reads from leaf, stem and bud tissues were mapped to the reference genome using HISAT v2.2.1 ^66^ with parameters “--max-intronlen 20000 --dta --score-min L, 0.0, −0.4”, and Trinity v2.8.4 ^67^ was used for transcripts assembly with default parameters. Assembled transcripts were subsequently aligned to the corresponding genome to predict gene structure using PASA v2.4.1 ^68^. For the *ab initio* prediction, Augustus v3.3.2 ^69^ was employed using default parameters after incorporating the transcriptome-based and homology-based evidence for gene model training. Finally, all predictions of gene models generated from these approaches were integrated into the final consensus gene set using EvidenceModelerv1.1.1 ^68^. After prediction, PASA was again used to update alternatively spliced isoforms to gene models and to produce a final gff3 file with three rounds of iteration.

In addition, we also performed noncoding RNAs (ncRNAs) annotation. Transfer RNAs (tRNAs) were identified using tRNAscan-SE v2.0.7 ^70^ with default parameters. Ribosomal RNAs (rRNAs) were identified by aligning rRNA genes of *P. trichocarpa*_v3.1 to the assembly using blast. The other three types of ncRNA (microRNA, small nuclear RNA and small nucleolar RNA) were identified using Infernal v1.1.4 ^71^ by searching Rfam database v12.0 ^72^.

For functional annotation, our predicted protein-coding genes were aligned to multiple public databases including NR, Swiss-Prot, TrEMBL ^73^, COG and KOG using NCBI BLAST+ v.2.2.31 with E-value of 1e-5 as cutoff ^64^. Motifs and domains were annotated by searching against InterProScan (release 5.32-71.0) ^74^. Gene ontology (GO) terms and KEGG pathways of predicted sequences were assigned by InterProScan and KEGG Automatic Annotation Server, respectively ^75^.

### Gene family clustering and phylogenetic analysis

Protein sequences from 13 plant species, including *Populus koreana, Populus euphratica, Populus pruinosa, Populus trichocarpa, Populus deltoides, Populus tremula, Populus alba, Salix suchowensis, Salix pruinosa, Ricinus communis, Arabidopsis thaliana, Vitis vinifera and Oryza sativa*, were selected for gene family clustering. Genes with premature stop codons or encoding proteins shorter than 50 amino acids were removed. For genes with alternative splicing variants, the longest transcript was selected to represent the gene. An all-against-all comparison was performed using BLASTP v2.5.0+ with e-value setting 1e^-5^, and OrthoFinder v2.5.2 ^76^ was used to further cluster gene families.

A total of 905 single-copy orthologous genes were extracted. The coding DNA sequence (CDS) alignments of each single-copy gene family were generated based on protein sequences aligned with MAFFT v7.475 ^77^ and poorly conserved blocks and gaps were trimmed by trimAl v1.4 ^78^ with default settings. Then, the consensus sequences were concatenated into a ‘super gene’ for each species, and RAxML v8.2.8 ^79^ was used to construct a phylogenetic tree under the GTRGAMMA model with 1000 bootstrap replicates, which was visualized by FigTree v1.4.4. Molecular dating was carried out using the MCMCTree program implemented in the PAML package v4.10.0 ^80^ based on the calibration time for divergence between *O. sativa* and *A. thaliana* (mean: 152 Mya) and between *A. thaliana* and *V. vinifera* (mean: 117 Mya) obtained from the TimeTree database (http://www.timetree.org) ^81^. Finally, we applied CAFE v4.2.1 ^82^ to compute changes in gene families along each lineage of the phylogenetic tree under a random birth-and-death model. The expanded and contracted gene families in *P. koreana* relative to other species were subjected to functional analysis using GO enrichment.

### Genome synteny and whole-genome duplication (WGD) analysis

We selected four species (*P. euphratica, P. trichocarpa, P. tremula, S. purpurea*) from Salicaceae to determine whether *P. koreana* shared the same whole-genome duplication events as other Salicaceae species. Colinear genes and syntenic blocks within each genome and between genomes were inferred using all-versus-all BLASTP and MCscan ^83^, with syntenic blocks being defined as those with at least five syntenic genes. Synonymous substitutions per synonymous site (*K*s) between colinear blocks was calculated for each pair of homologous genes using WGDI v0.4.5 ^84^. The median *K*s values of each syntenic block were then selected and used for the distribution analysis after performing the evolutionary rate correction.

### Genome resequencing, read mapping and variant calling

A total of 230 individuals were collected from 24 natural populations, representing most natural habitats of *P. koreana*. Within each population, individuals were sampled after ensuring that sampled individuals were at least 100m apart from each other. Genomic DNA was extracted from leaf samples with Qiagen DNeasy plant kit. Whole genome paired-end sequencing was generated using the Illumina NovaSeq 6000 platform with a target coverage of 20× per individual.

For raw resequencing reads, we used Trimmomatic v0.36 ^85^ to remove adapters and cut off bases from either the start or the end of reads if the base quality was < 20. Trimmed reads shorter than 36 bases were further discarded. After quality control, all high-quality reads were mapped to our *de novo* assembled *P. koreana* genome using the BWA-MEM algorithm of bwa v.0.7.17 ^59^ with default parameters. The alignment results were then processed by sorting and PCR duplicate marking using SAMtools v.1.9 ^86^ and Picard v.2.18.11 (http://broadinstitute.github.io/picard/). For genetic variant identification, SNP and Indel calling was performed using Genome Analysis Toolkit (GATK v.4.0.5.1) ^87^ and its subcomponents HaplotypeCaller, CombineGVCFs and GenotypeGVCFs to form a merged VCF file with “all sites” (including nonvariant sites) included using the ‘EMIT_ALL_SITES’ flag. SV calling was performed using the software DELLY v0.8.3 ^88^ with default parameters. We further performed multiple filtering steps to only retain high-quality variants for downstream analysis. For SNPs, SNPs with multi-alleles (>2) and those located at or within 5 bp from any indels were removed. In addition, after treating genotypes with read depth (DP) < 5 and genotype quality (GQ) < 10 as missing, SNPs with missing rate higher than 20% were filtered; for indels, those with muti-alleles (>2) and with QD < 2.0, FS > 200.0, SOR > 10.0, MQRankSum < −12.5, ReadPosRankSum < −8.0 were removed. Indels with missing rate >20% after treating genotype with DP<5 and GQ<10 as missing were further filtered out; for SVs, those with length < 50bp and with imprecise breakpoints (flag IMPRECISE) were removed. After treating genotypes with GQ<10 as missing, we further filtered SVs with missing rate >20%. Finally, we implemented the software SNPable (http://lh3lh3.users.sourceforge.net/snpable.shtml) to mask genomic regions where reads were not uniquely mapped and filtered out variants located in these regions. After these filtering steps, 16,619,620 SNPs, 2,663,202 indels and 90,357 SVs were remained for subsequent analyses. The filtered variants were further phased and imputed using Beagle v4.1 ^89^ and the effects of individual variants were annotated using SnpEff v.4.3 ^90^ with “-ud 2000” and other parameters set to default.

### Population structure analysis

We first used PLINK v1.90 ^91^ with the parameters “indep-pairwise 50 10 0.2” to extract a LD pruned SNP set with minor allele frequency (MAF) > 5%, which yielded 535,191 independent SNPs to be used in the population structure analysis. First, we used ADMIXTURE v.1.3.0 ^92^ with default parameters to investigate population genetic structure across all individuals, with the number of clusters (K) being set from 1 to 8. Second, to quantify the relatedness between individuals, the identify-by-state (IBS) genetic distance matrix was calculated using “-distance 1-ibs” parameter in PLINK v1.90. We constructed a neighbor-joining (NJ) phylogenetic tree based on the distance matrix using MEGAX ^93^ and displayed the tree using FigTree v.1.4.4. Third, for the isolation-by-distance (IBD) analysis, we first used VCFtools v0.1.15 ^94^ to calculate the population differentiation coefficient (*F*_ST_). The matrix of *F*_ST_ (*F*_ST_ (1− *F*_ST_)) and the matrix of geographic distance (km) among different groups of populations were then used for performing the Mantel tests using the R package “vegan” ^95^, with the significance being determined based on 999 permutations.

### Genetic diversity, linkage disequilibrium and demographic history analysis

To estimate and compare genetic diversity across populations of *P. koreana*, we calculated both intra-population (π) and inter-population (d_xy_) nucleotide diversity after taking into account both polymorphic and monomorphic sites using the program pixy v0.95.0 ^96^ over 100 Kbp nonoverlapping windows. In addition, Tajima’s D statistics were calculated using VCFtools v0.1.15 in 100 Kbp non-overlapping windows for the northern and southern groups of populations, respectively. To further estimate and compare the pattern of LD among different groups of populations, PopLDdecay v.3.40 ^97^ was used to calculate the squared correlation coefficient (*r*^2^) between pairwise SNPs with MAF >0.1 in a 100-kb window and then averaged across the whole genome.

PSMC ^98^ was used to infer historical changes in effective population size (*N*_e_) of *P. koreana* using parameters of -N25 -t15 -r5 -p “4+25*2+4+6”. We selected seven individuals from both the northern and southern groups of populations to run the PSMC analyses, and 100 bootstrap estimates were performed per individual. Assuming a generation time of 15 years and a mutation rate of 3.75×10^−8^ mutations per generation, we converted the scaled population parameters into *N*_e_ and years.

### Identification of environment-associated genetic variants

We used two different approaches to identify environment-associated variants (SNPs, indels, and SVs) across the whole genome. We only kept common variants with MAF >10%, including a total of 5,182,474 SNPs, 736,051 indels and 30,934 SVs, for these analyses. First, we used a univariate latent-factor linear mixed model (LFMM) implemented in the R package LEA v3.3.2 ^99^ to search for associations between allele frequencies and the 19 BIOCLIM environmental variables ^100^. Based on the number of ancestry clusters inferred with ADMIXTURE v.1.3.0, we ran LFMM with three latent factors to account for population structure in the genotype data. For each environmental variable, we ran five independent MCMC runs using 5000 iterations as burn-in followed by 10,000 iterations. *P*-values from all five runs were then averaged for each variant and adjusted for multiple tests using a false discovery rate (FDR) correction of 5% as the significance cutoff. Second, we performed a redundancy analysis (RDA) to identify genetic variants showing especially strong relationship with multivariate environmental axes ^29,101^. RDA has been demonstrated to be one of the best-performing multivariant genotype-environmental association approaches and which exhibits low false-positive rates ^29^. Six uncorrelated environmental variables (BIO1, BIO3, BIO5, BIO13, BIO15 and BIO19) with pairwise correlation coefficients <0.6 were selected for the RDA analyses using the R package vegan v2.5-7. Significant environment-associated variants were defined as those having loadings in the tails of the distribution using a standard deviation cut-off of 3 along one or more RDA axes.

To further assess selection pressures acting on climate adaptive variants, we assessed the extended haplotype homozygosity (EHH) pattern for a selected set of strongly associated variants using the R package “rehh” ^102^, and calculated the standardized integrated haplotype score (iHS) across the genome for common variants using the software selscan v.1.3.0 ^103^.

### Stress treatment and expression analysis by qRT-PCR

Stem segments from wild genotypes of *P. koreana* were surface sterilized by soaking in 10% sodium hypochlorite solution and 70% Ethyl alcohol for 5 minutes, and then thoroughly washed five times with distilled water. The stem segments were inserted into MS medium (0.05mg/L NAA) for 30 d at 25/20 °C (day 16 h/night 8 h) and after rooting, the stem segments were transplanted to soil for 40 d at 25/20 °C (day 16 h/night 8 h). To explore the effect of different genotypes of one candidate adaptive SNP located in the 5’ UTR of *Pokor12247* (LG04:25159299) in mediating adaptation to extreme precipitation, we carried out a submergence treatment. For the submergence treatment, water was maintained at 2 cm above the soil surface and plants were maintained in the growth chamber providing 25 °C/20 °C (day 16 h/night 8 h) for 0h, 3h, 6h, 9h and 12h. In addition, we also carried out a heat stress treatment to explore the effect of one candidate adaptive SNP located in intronic region of *Pokor17228* (LG07: 4796402) in response to heat stress. For the heat stress treatment, plants were placed into a plant incubator at 42 °C/20 °C (day/night) with an illumination of 16 h/8 h (day/night) for 0 h, 1 h, 2 h, 3 h and 24 h. At each time point, leaf tissues were collected from each plant at the same place and frozen immediately in liquid nitrogen for expression analyses.

Quantitative Reverse Transcription PCR (qRT-PCR) ^104^ was used to investigate the expression levels of selected genes in the abiotic treatments (*Pokor12247* for submergence stress; *Pokor17228* for heat stress). Total RNA was extracted from pooled leaf materials using a Plant RNA extract kit (Biofit, Chengdu, China), and the HiScript II RT SuperMix for qPCR kit (+gDNA wiper) (Vazyme, Nanjing, China) was used to obtain cDNA. qPCR was performed with gene-specific primers (Supplementary Table 18) using the Taq Pro Universal SYBR qPCR Master Mix (Vazyme, Nanjing, China) reaction system on the CFX96 Real-Time detection system (Bio-Rad, CA, USA). Each experiment was performed with three technical replicates and the *UBQ10* was used as the endogenous control for data analysis.

### ATAC-seq analysis

For the ATAC experiment, fresh leaf tissue were collected from the same individual used for the genome-assembly of *P. koreana* and prepared according to the experimental protocol following ^105^. In brief, approximately 500mg of flash-frozen leaves were immediately chopped and processed for ATAC-seq, followed by library construction and were then subjected to sequencing on the Illumina HiSeq X-Ten platform (San Diego, CA, USA). The raw reads generated were first trimmed using Trimmomatic v.0.36 ^85^ with a maximum of two seed mismatches, and the adapters were trimmed by NexteraPE. Then the clean reads were aligned to the reference genome using Bowtie v.2.3.2 ^56^ using the following parameters: ‘bowtie2 --very-sensitive -N 1 -p 4 -X 2000 -q’. Aligned reads were sorted using SAMtools v.1.1.1 ^86^. The redundant reads from PCR amplification and reads that mapped to either chloroplast or mitochondria were removed using Picard v.2.18.11 (http://broadinstitute.github.io/picard/). Finally, only high quality properly paired reads were retained for further analysis. ATAC-seq peak calling was done by MACS2 ^106^ with the ‘-keep dup all’ function.

### Genomic vulnerability assessment

For each sampling location, we downloaded future (2061-2080 and 2081-2100) environmental data for the 19 BIOCLIM variables from WorldClim CMIP6 dataset (BCC-CSM2-MR model; resolution 2.5 arcmin) ^100^. Each of the two future environmental datasets consists of four Shared Socio-economic Pathways (SSPs): SSP126, SSP245, SSP370 and SSP585. We used two different approaches to evaluate the genomic vulnerability to future climate change. First, we calculated the risk of nonadaptedness (RONA) ^16^, which quantifies the theoretical average change in allele frequency needed to cope with climate change, under projected future climate scenarios. Following the method used in ^107^, a linear relationship between allele frequencies at significantly associated loci (detected by both LFMM and RDA) and environmental variables was first established using linear regressions. For each locus, population and environmental variable, the theoretical allele frequency change needed to cope with future climate conditions (RONA) were calculated, and the average RONA values were further weighted by the *R*^2^ for each linear regression following ^46^. Second, as a complementary approach to RONA, we used a nonparametric, machine-learning gradient forest analysis to calculate genomic vulnerability across the range of *P. koreana* using ‘gradientForest’ in R ^10,108^. We first built a GF model with 500 trees on the 19 BIOCLIM variables using the environmental-associated variants detected by both LFMM and RDA, which provided a ranked list of the relative importance of all environmental variables. Based on the ranked importance and pairwise correlation coefficients of the nineteen variables, we selected six unrelated environmental variables (BIO1, BIO3, BIO5, BIO13, BIO15 and BIO19, identical to the RDA analyses) to build a second gradient forest model for estimating the genetic offset under the different future scenarios. The genetic offset was calculated as a metric for the Euclidean distance of the genomic composition between the current and future projected climates, and then mapped with ArcGIS 10.2 to display its’ geographical distribution.

## Funding

This project was supported by National Natural Science Foundation of China (31971567) and Fundamental Research Funds for the Central Universities (YJ201936, SCU2020D003, SCU2021D006).

## Author contributions

J.W., K.M. and J.L. conceived the research. J.W. supervised the study. Y.S., H.Z., K.M. performed the sampling and collected the materials. Y.S., Z.L., T.S., C.J. X.Z., Q.L., G.Y., X.X. conducted all bioinformatics analyses. X.D., J.F., H.L. and Y.J. performed the experiment. Y.S. Z.L. and J.W. wrote the manuscript, with the input from P.K.I and J.L. All authors approved the final manuscript.

## Competing interests

The authors declare that they have no competing interests.

## Data availability

All data needed to evaluate the conclusions in the paper are present in the paper and/or the Supplementary Materials. All sequencing data in this study will be deposited in National Genomics Data Center (NGDC) and/or NCBI during reviewing process. All scripts used in this study will be available at https://github.com/jingwanglab/Populus_genomic_prediction_climate_vulnerability upon publication.

## Notes

### Competing Interest Statement

The authors have declared no competing interest.

