## Supplemental figures and tables for "Genomic insights into present local adaptation and future climate change vulnerability of a keystone forest tree species in East Asian"

**This PDF file includes:**

Figs. S1 to S25
Tables S1 to S10, S13, S15 to S18

**Other Supplementary Materials for this manuscript include the following:**

Table S11, S12, S14 and Table S17 as separate EXCEL files

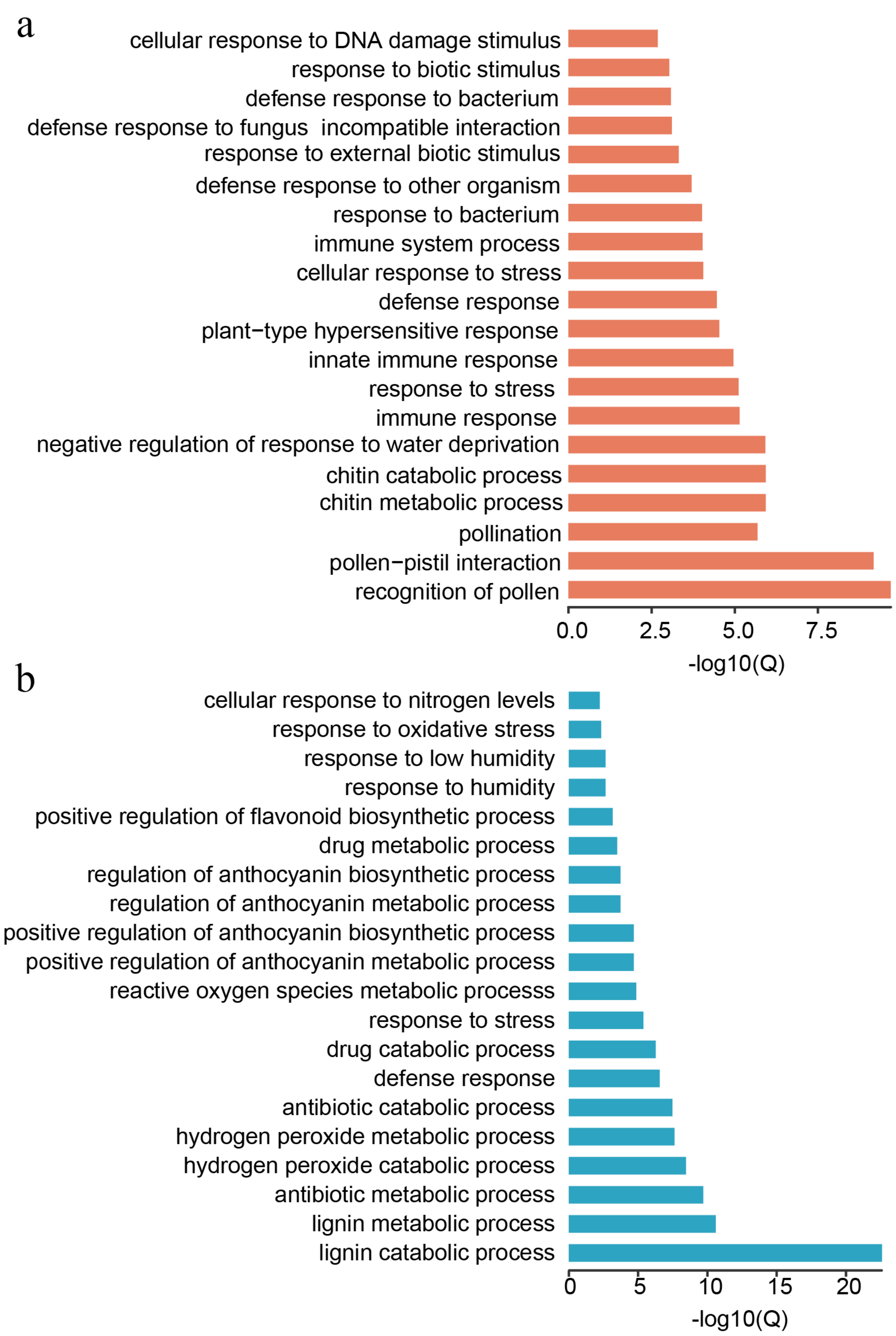


**Supplementary Fig. 1.** Gene Ontology (GO) enrichment analysis for expanded (a) and contracted (b) gene families in *P. koreana* genome.

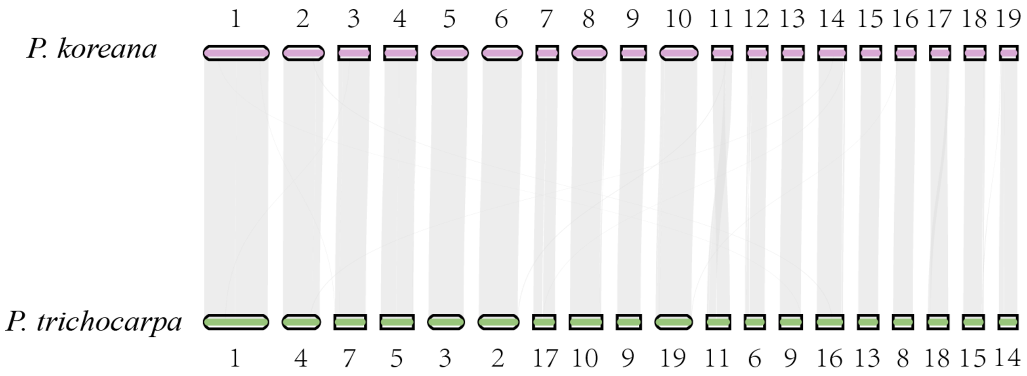


**Supplementary Fig. 2.** Genomic collinearity between *P. koreana* and *P. trichocarpa*.

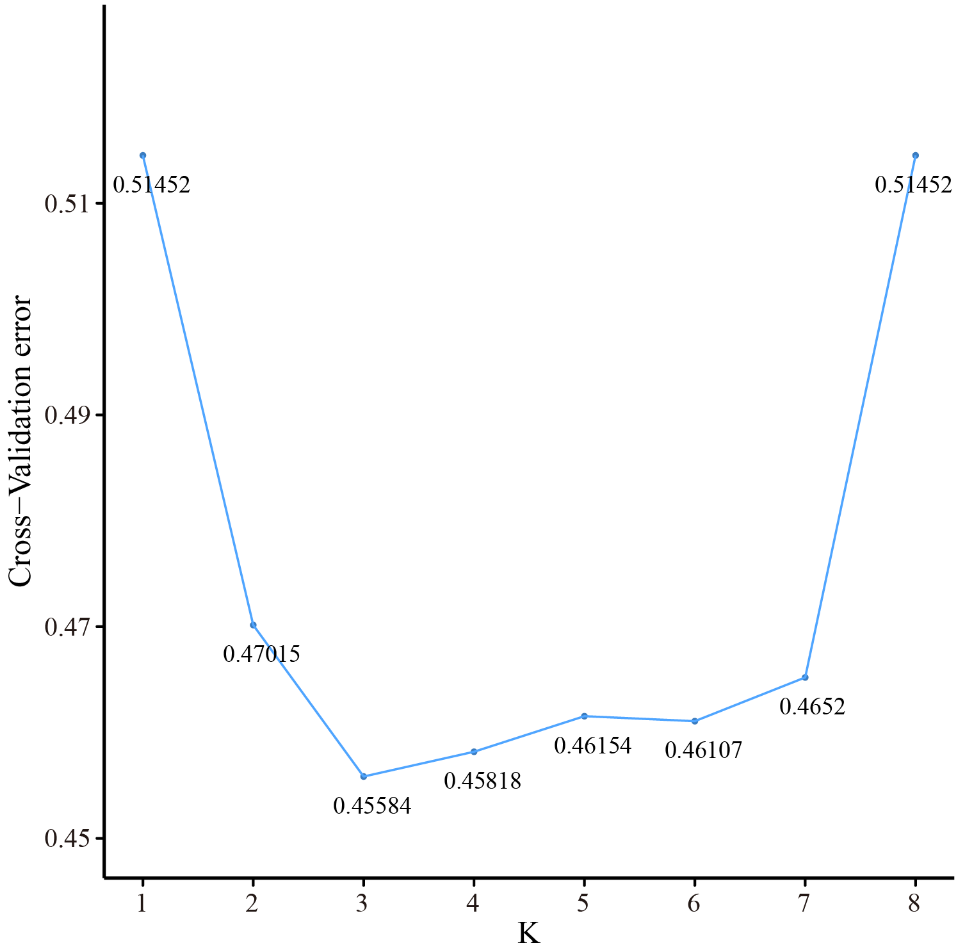


**Supplementary Fig. 3.** The Cross-Validation error distribution according to the number of clusters (K) by ADMIXTURE.

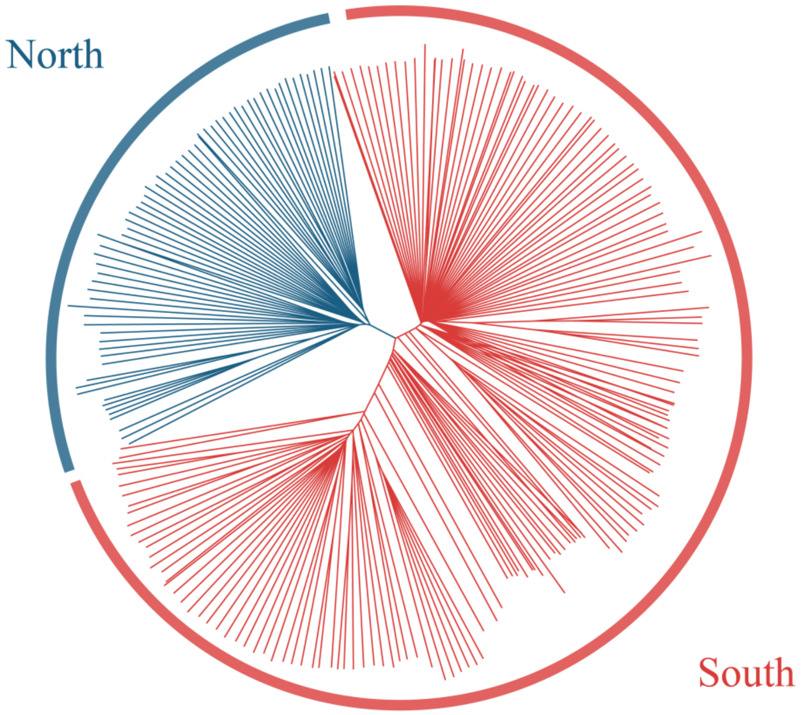


**Supplementary Fig. 4.** Neighbor-joining (NJ) phylogenetic tree of all 230 *P. koreana* individuals constructed using a whole-genome pruned SNP dataset. Two major clades (North and South) are indicated.

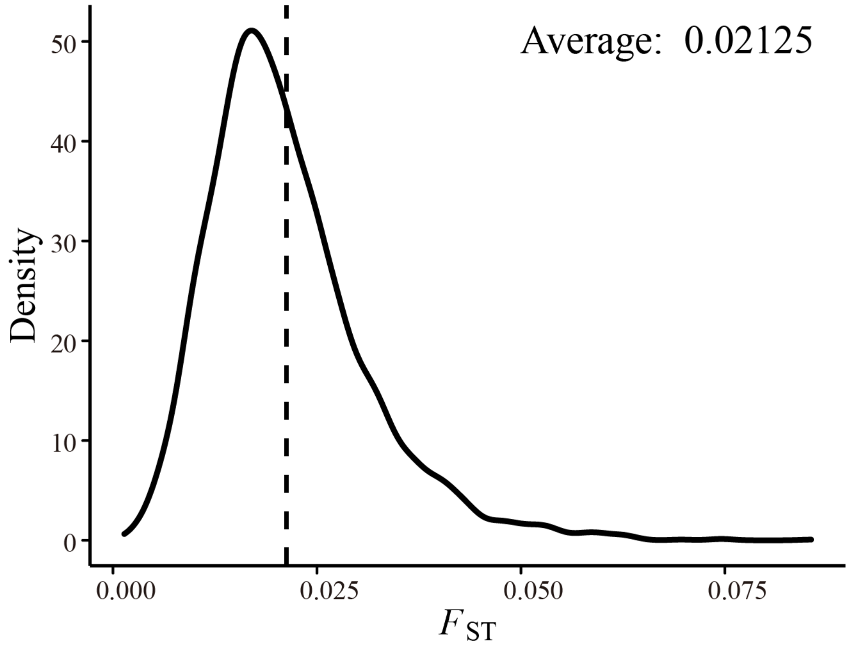


**Supplementary Fig. 5.** The distribution of $F_{ST}$ values between north and south groups of populations over 100 Kbp non-overlapping windows across the genome. The dashed line indicates the average $F_{ST}$ value.

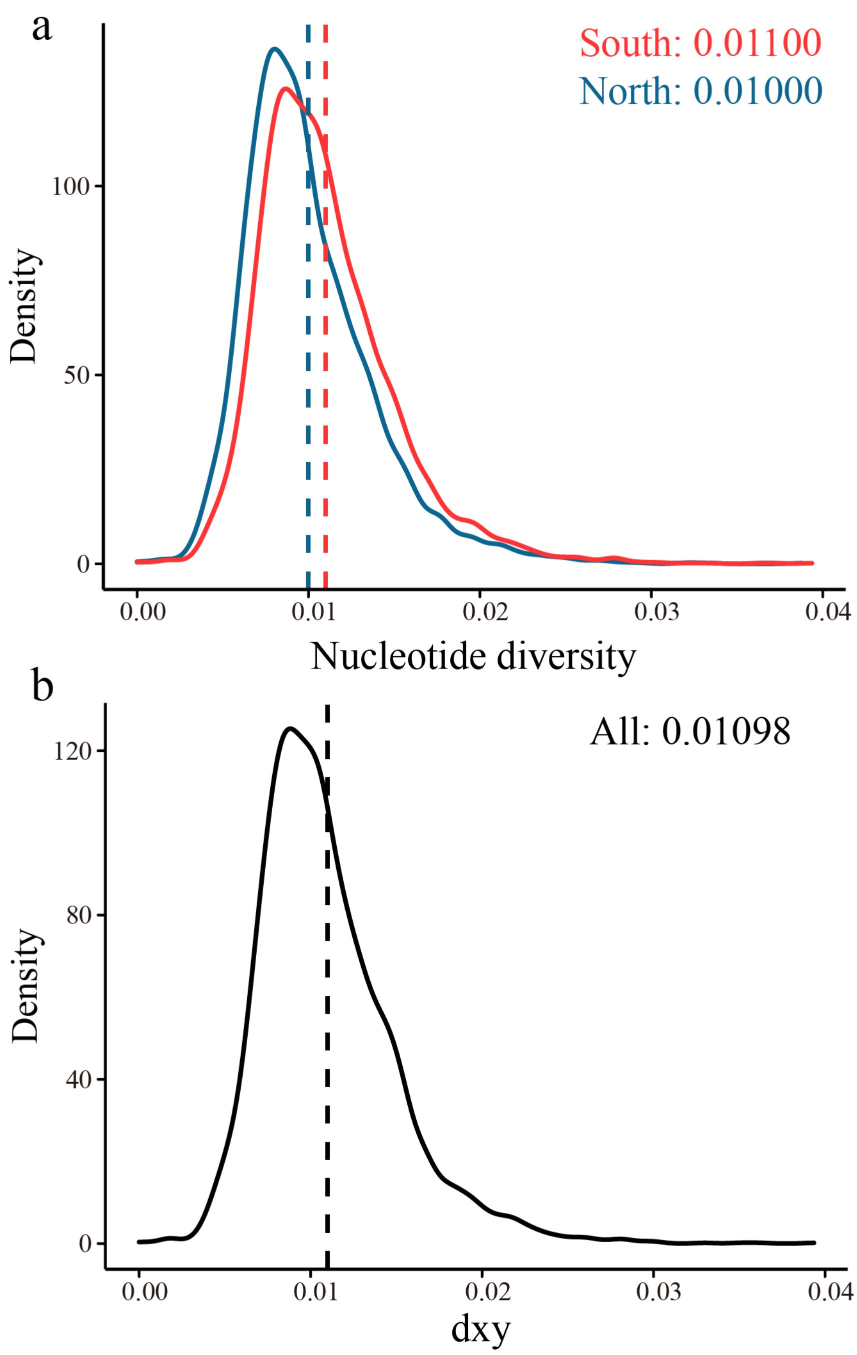


**Supplementary Fig. 6.** (a) The nucleotide diversity (π) estimates of south (red) and north (blue) groups of populations over 100 Kbp non-overlapping windows across the genome. (b) The nucleotide divergence (dxy) between south and north groups of populations over 100 Kbp non-overlapping windows across the genome. The dashed lines indicate the average estimates.

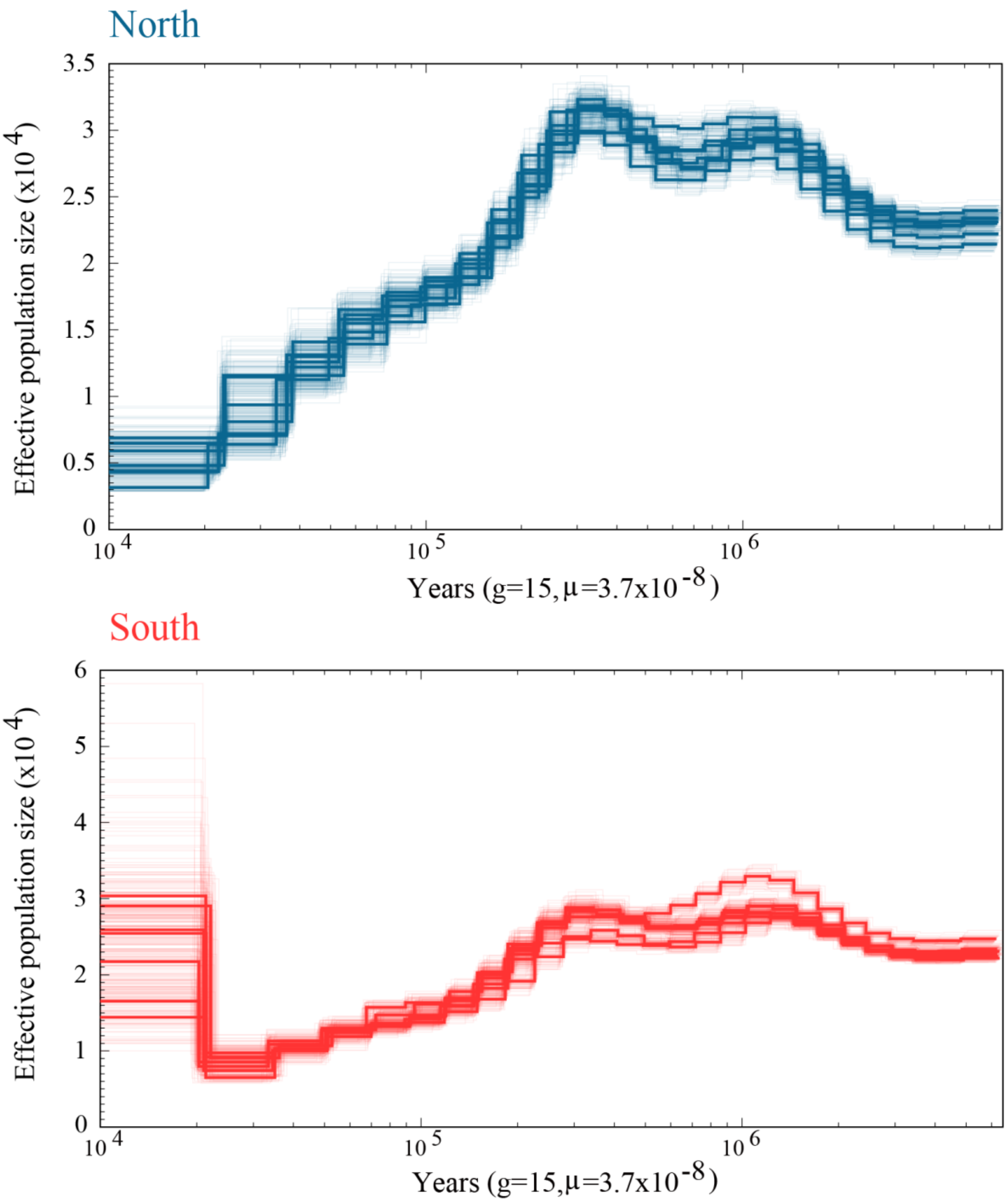


**Supplementary Fig. 7.** Demographic analysis for two groups (South and North) of populations of *P. koreana*. Thick lines are the estimate values of 7 selected individuals from each group, whereas faint lines are individual bootstrap replicates, 50 replicates were conducted per individual.

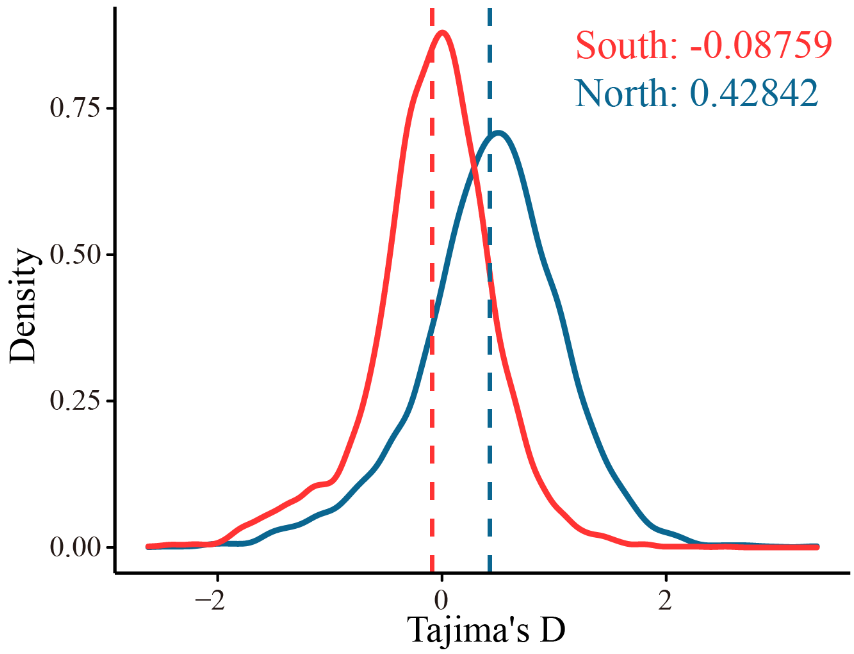


**Supplementary Fig. 8.** The Tajima's D statistics of south (red) and north (blue) groups of populations over 100 Kbp non-overlapping windows across the genome. The dashed lines indicate the average estimates.

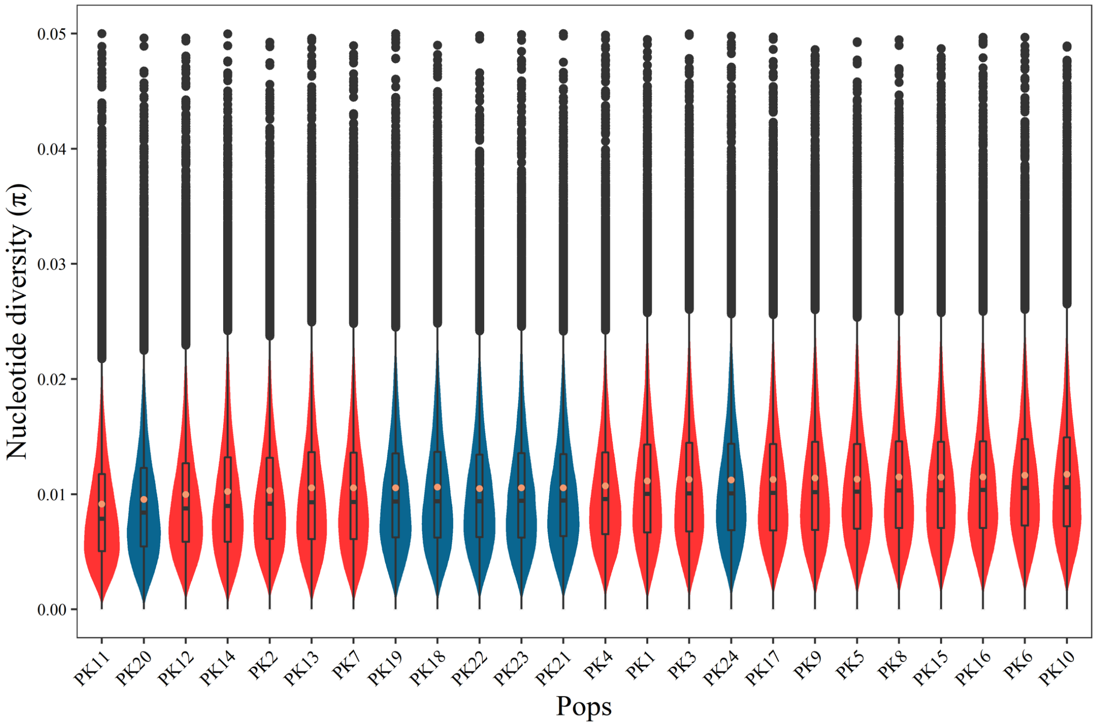


**Supplementary Fig. 9.** The distribution of nucleotide diversity over 10 Kbp non-overlapping windows across the 24 *P. koreana* populations.

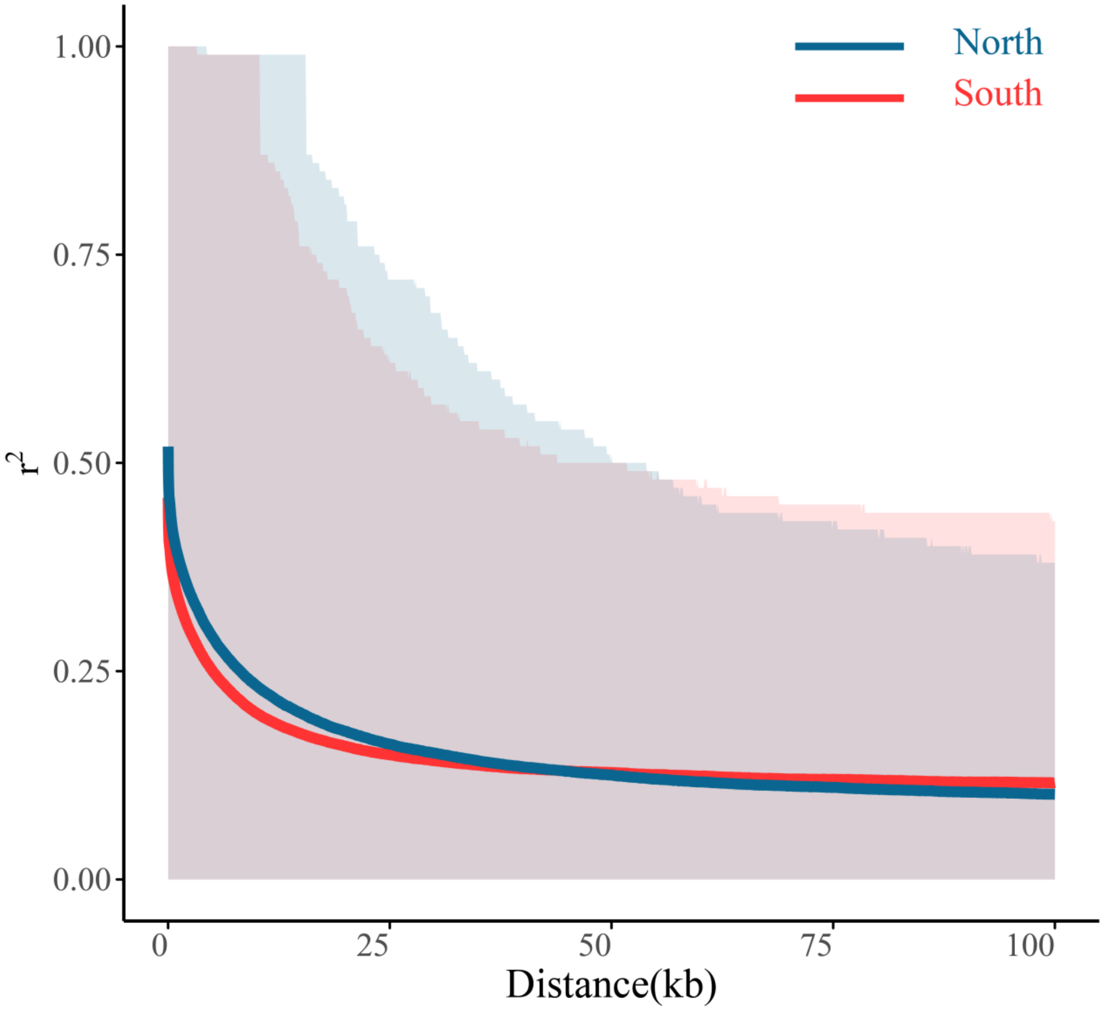


**Supplementary Fig. 10.** Linkage disequilibrium (LD) decay estimated by PopLDdecay for two groups of *P. koreana* (thick lines) with the 90% ranges (shadows, 5% to 95% percentiles) of $r^2$ values.

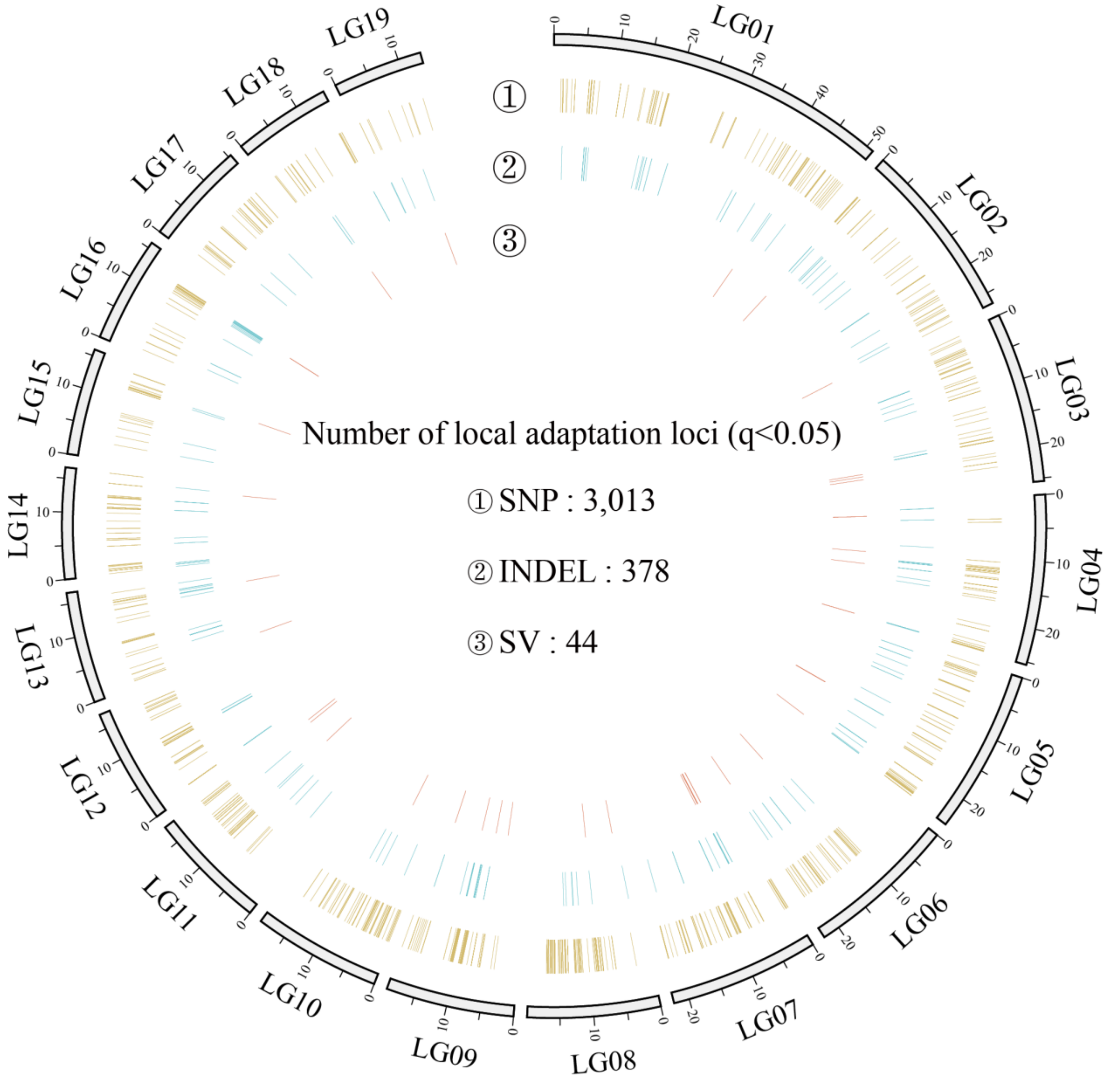


**Supplementary Fig. 11.** Distribution of 3,435 environmental-associated variants identified by LFMM across the genome.

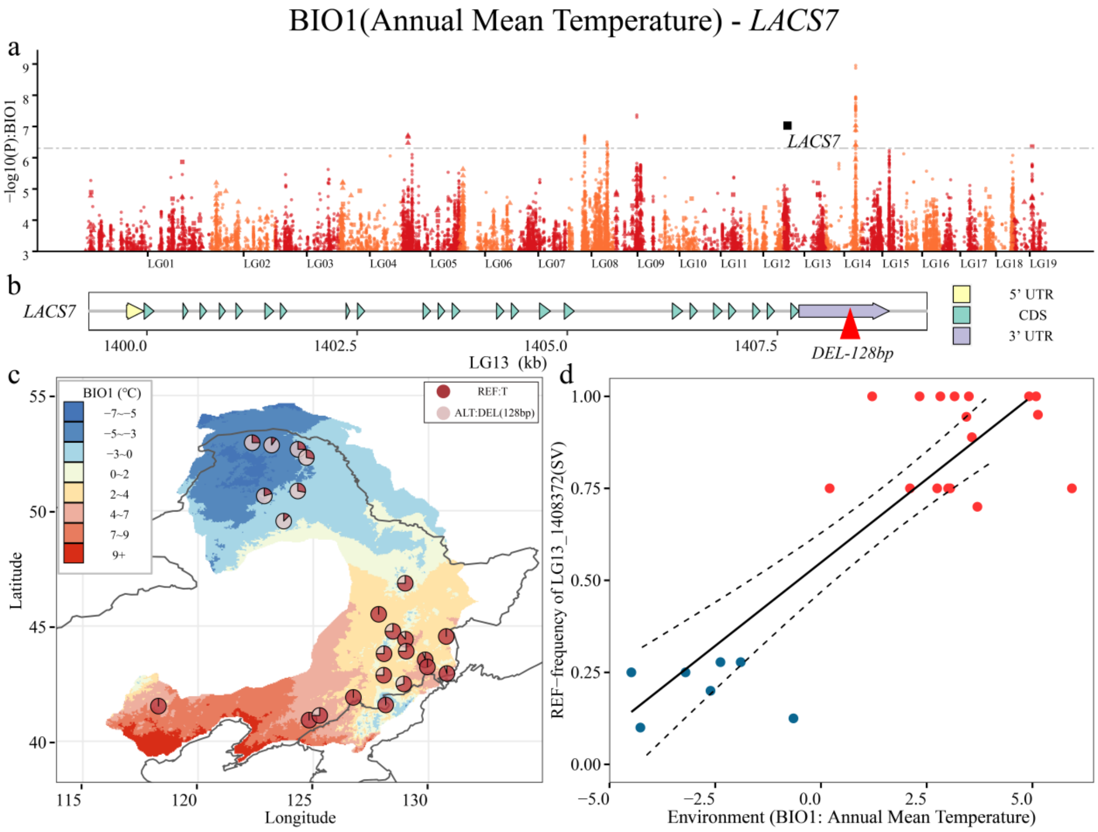


**Supplementary Fig. 12. Genome-wide variation associated with different environmental variables (BIO5 and BIO13 are not included). a** Manhattan plot shows the genotype-environment association estimated with LFMM. SNPs, Indels and SVs are represented by points, triangles and squares, respectively. The grey dashed line represents q = 0.05. Colors distinguish different chromosomes. **b** Schematic diagram of the structure of the gene associated with the variant marked in **a**. **c** Distribution of allele frequencies among 24 populations of the significant variant marked in (b). Colors of raster on map represent the environmental variable under current scenario. **d** Linear regression of allele frequencies and environment variables for the selected variant.

**Continuation Supplementary Fig. 12.**

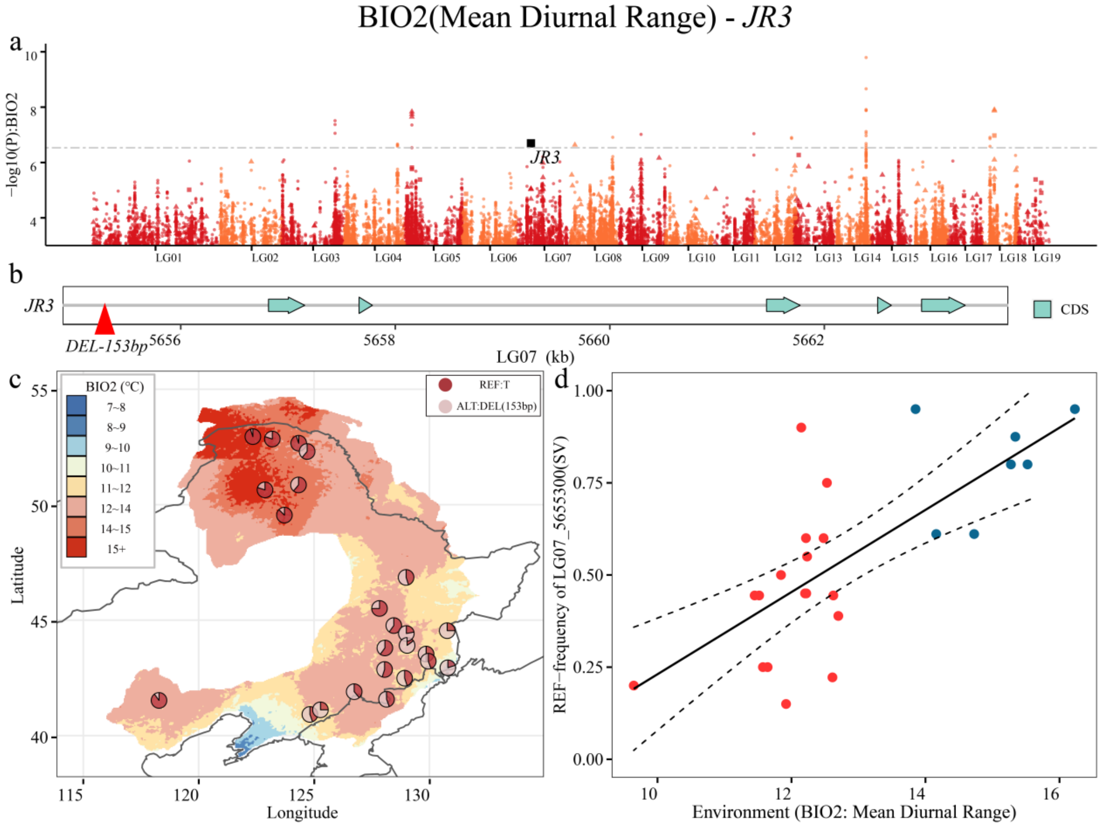

**Continuation Supplementary Fig. 12.**

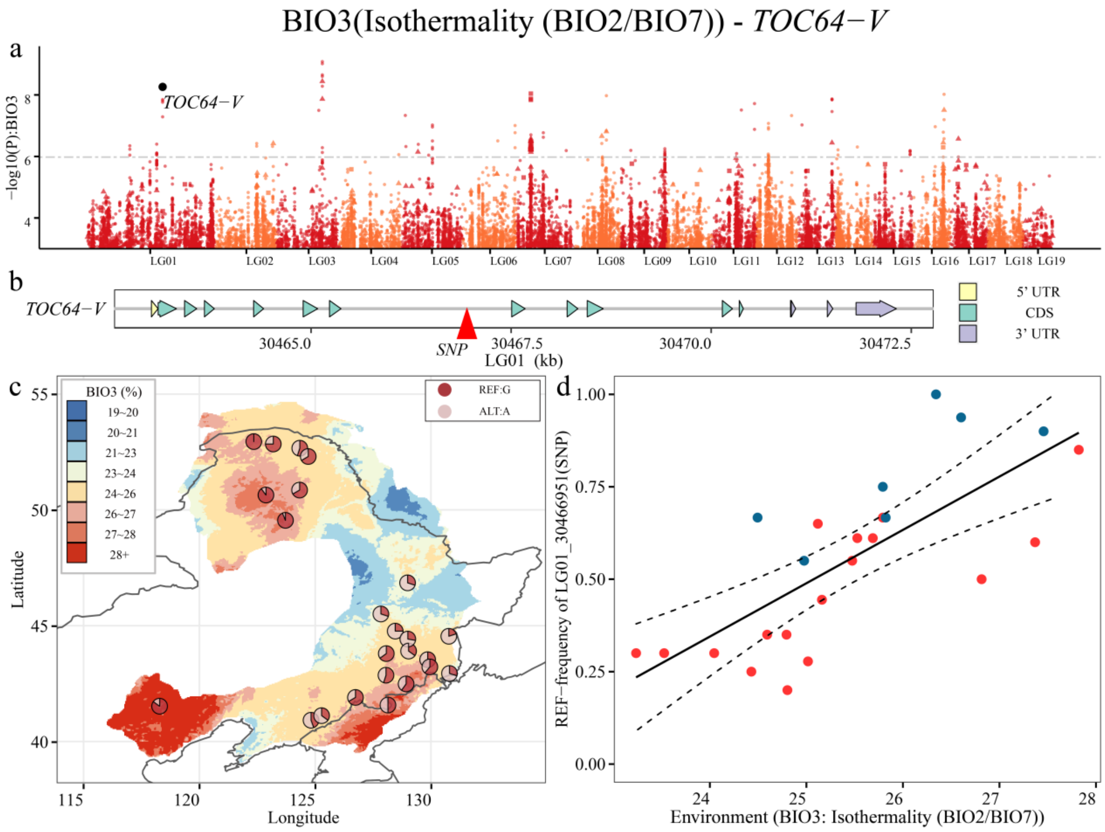

**Continuation Supplementary Fig. 12.**

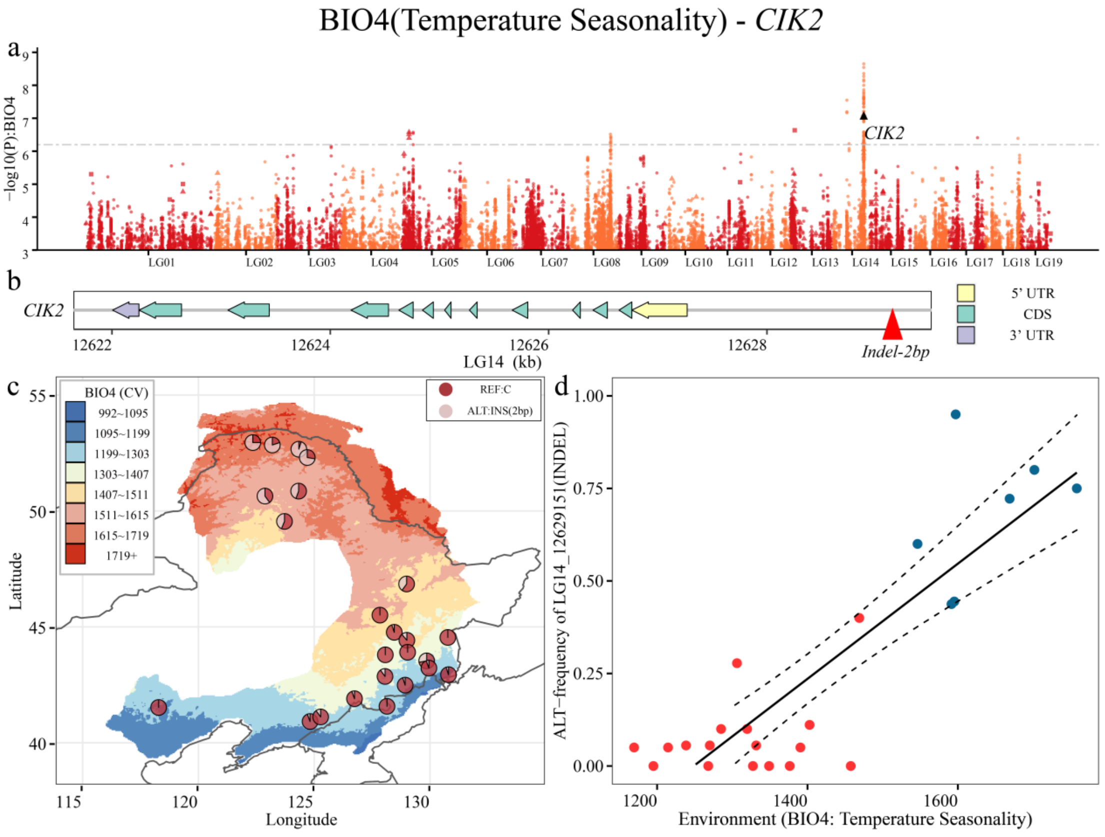

**Continuation Supplementary Fig. 12.**

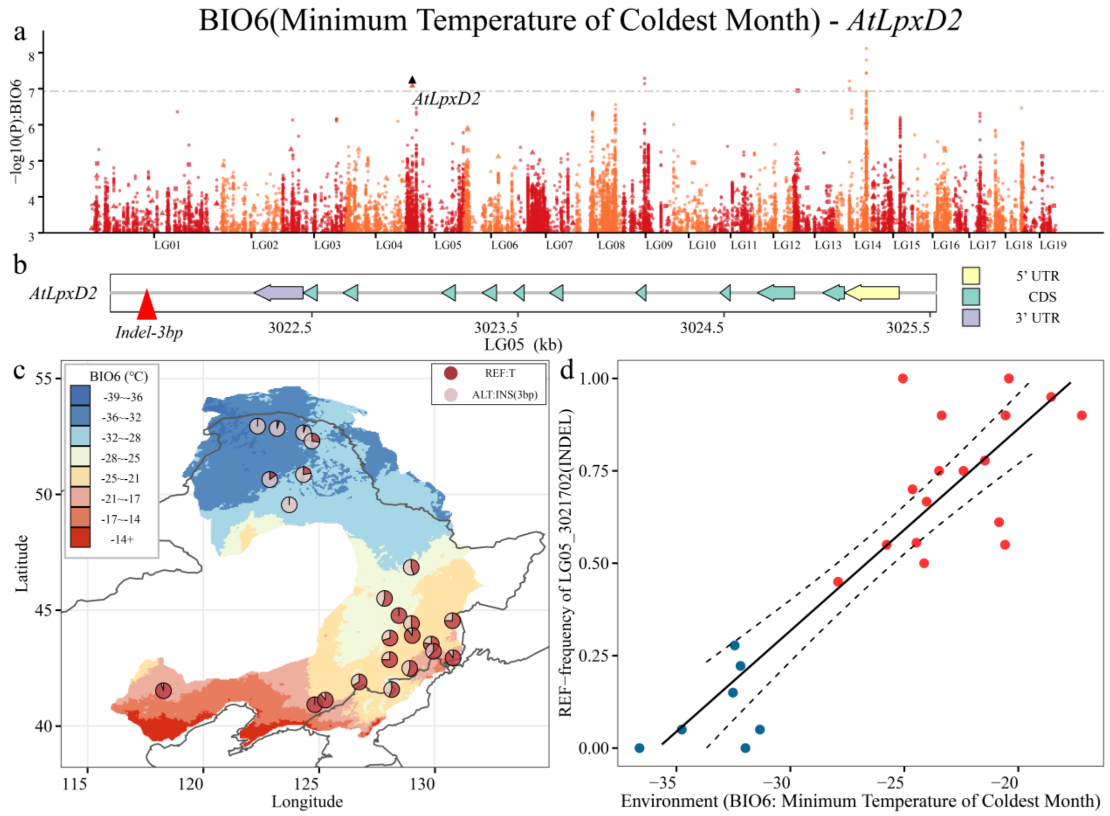

**Continuation Supplementary Fig. 12.**

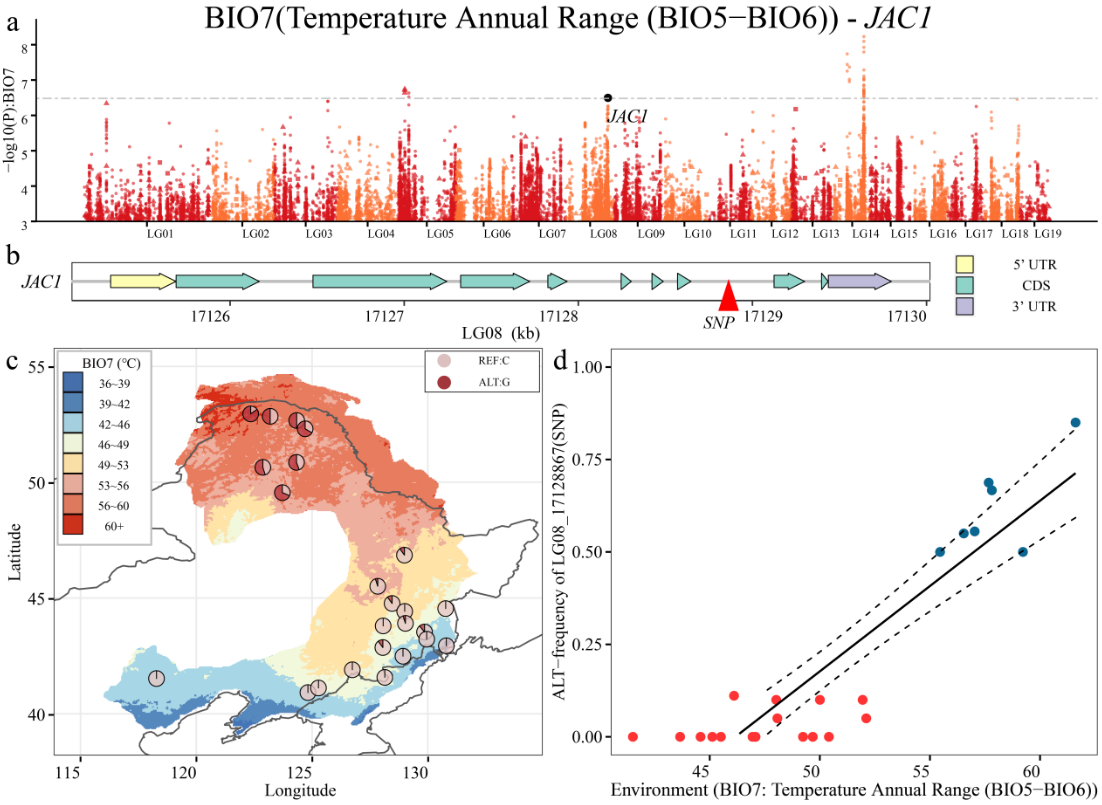

**Continuation Supplementary Fig. 12.**

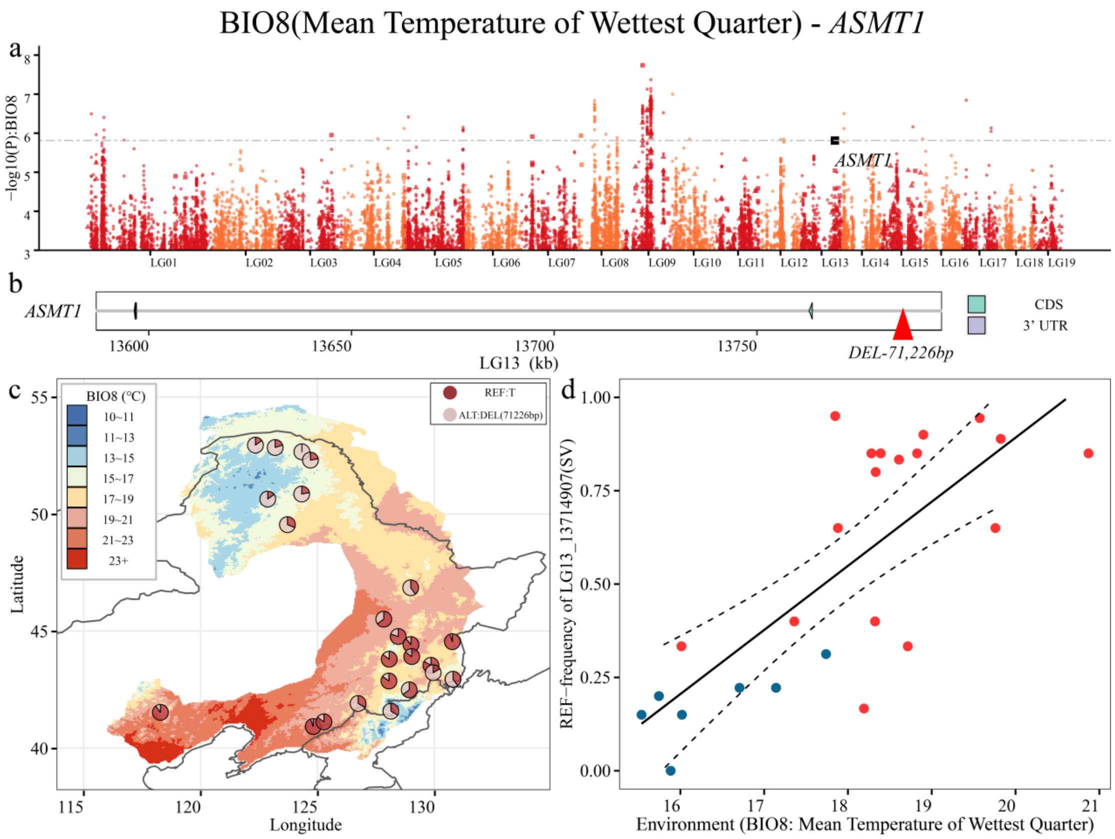

**Continuation Supplementary Fig. 12.**

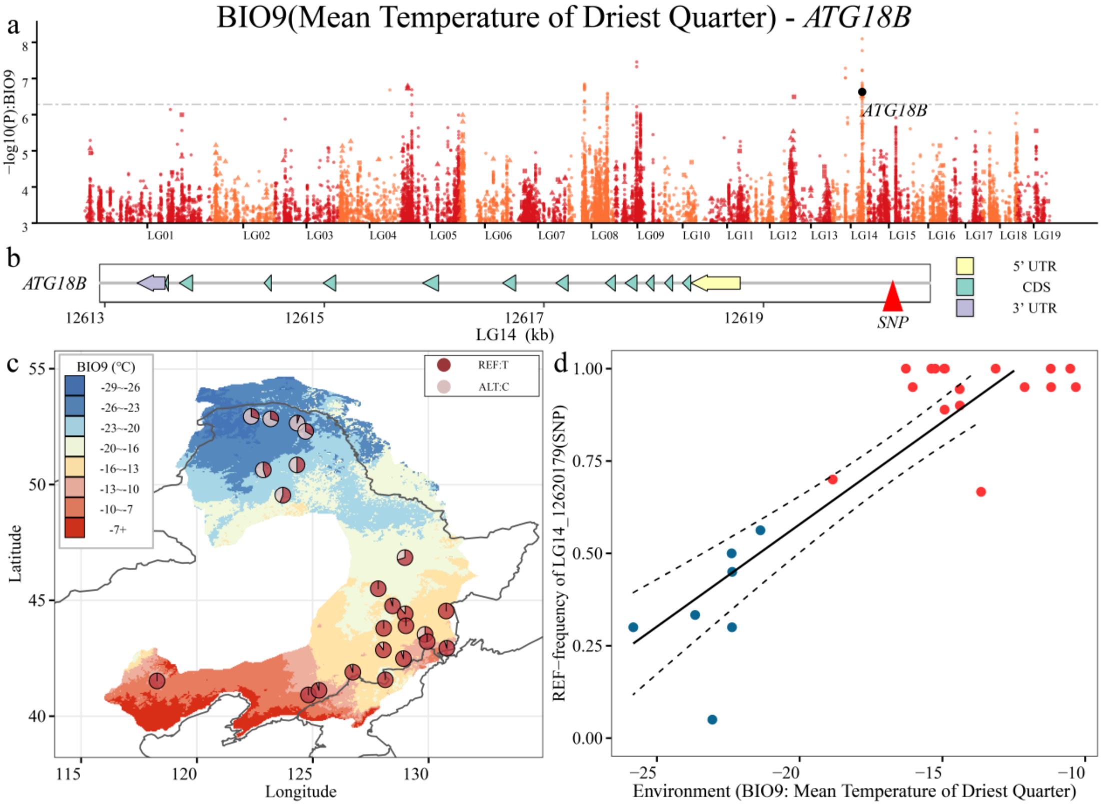

**Continuation Supplementary Fig. 12.**

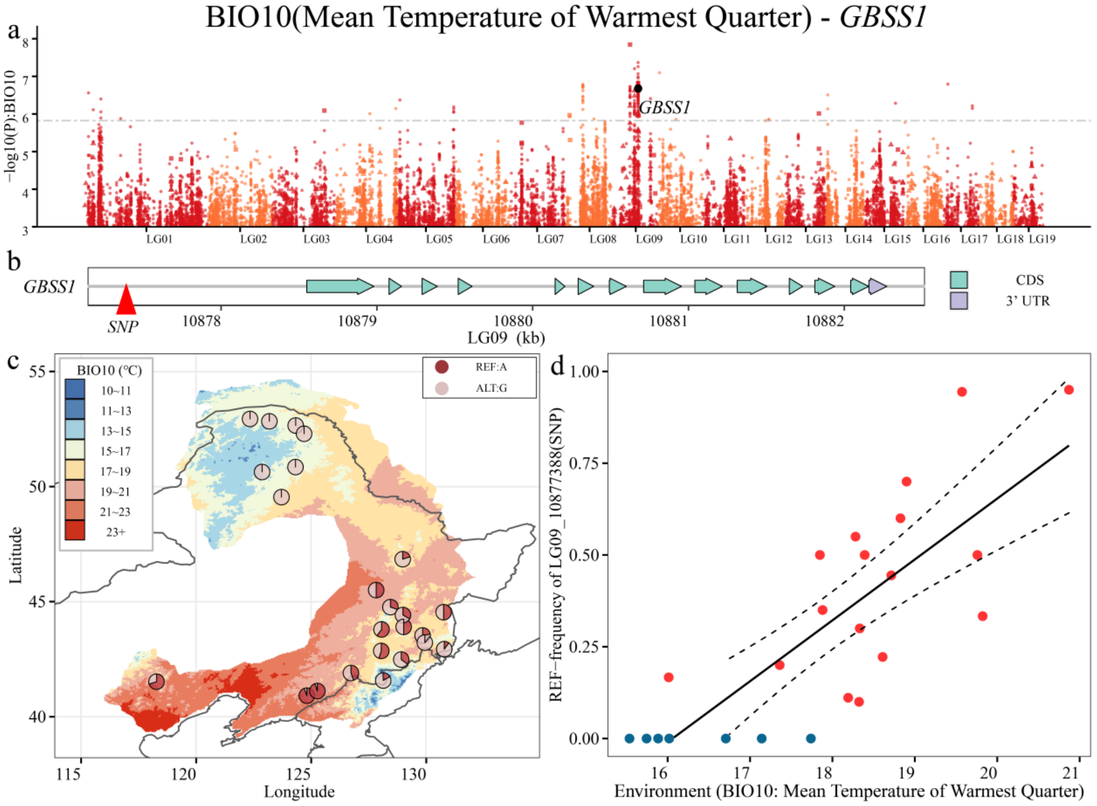

**Continuation Supplementary Fig. 12.**

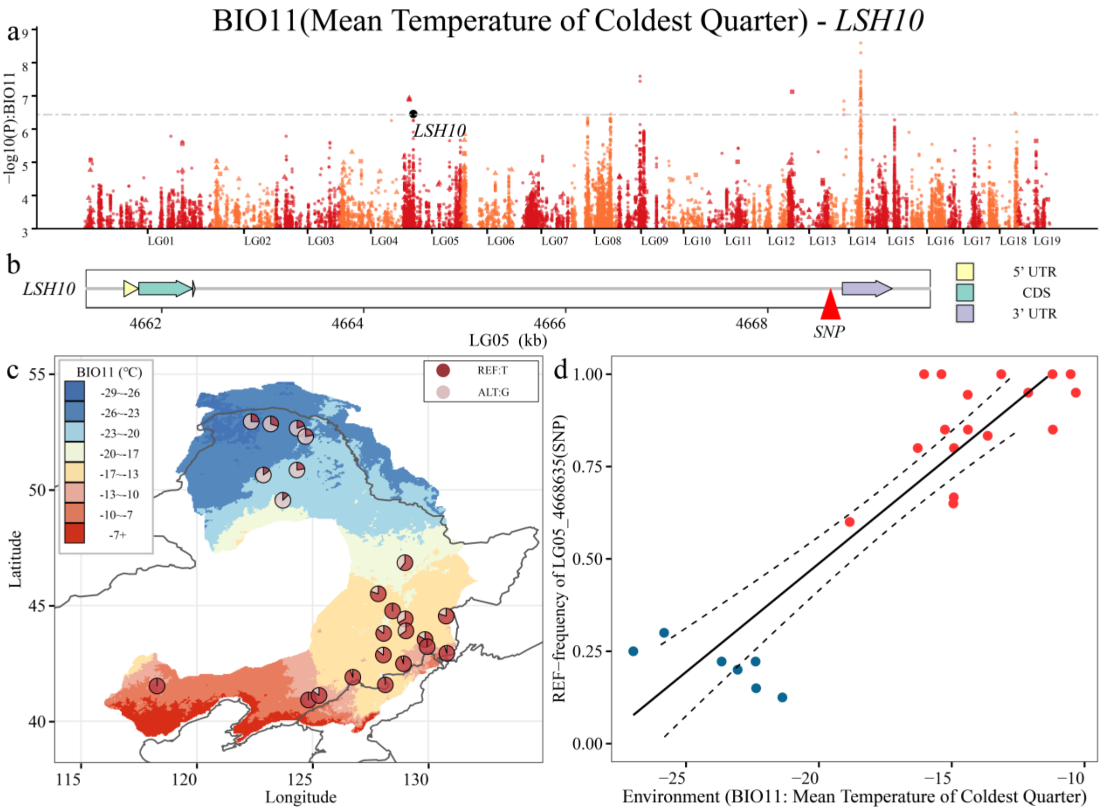

**Continuation Supplementary Fig. 12.**

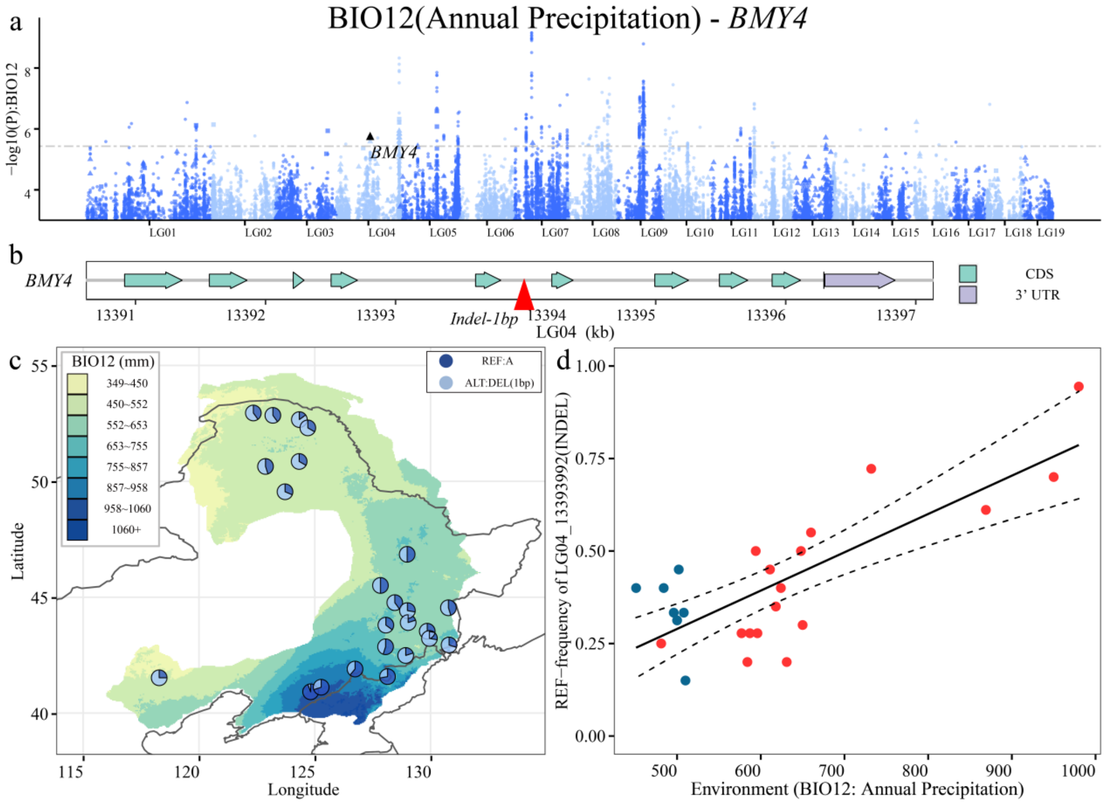

**Continuation Supplementary Fig. 12.**

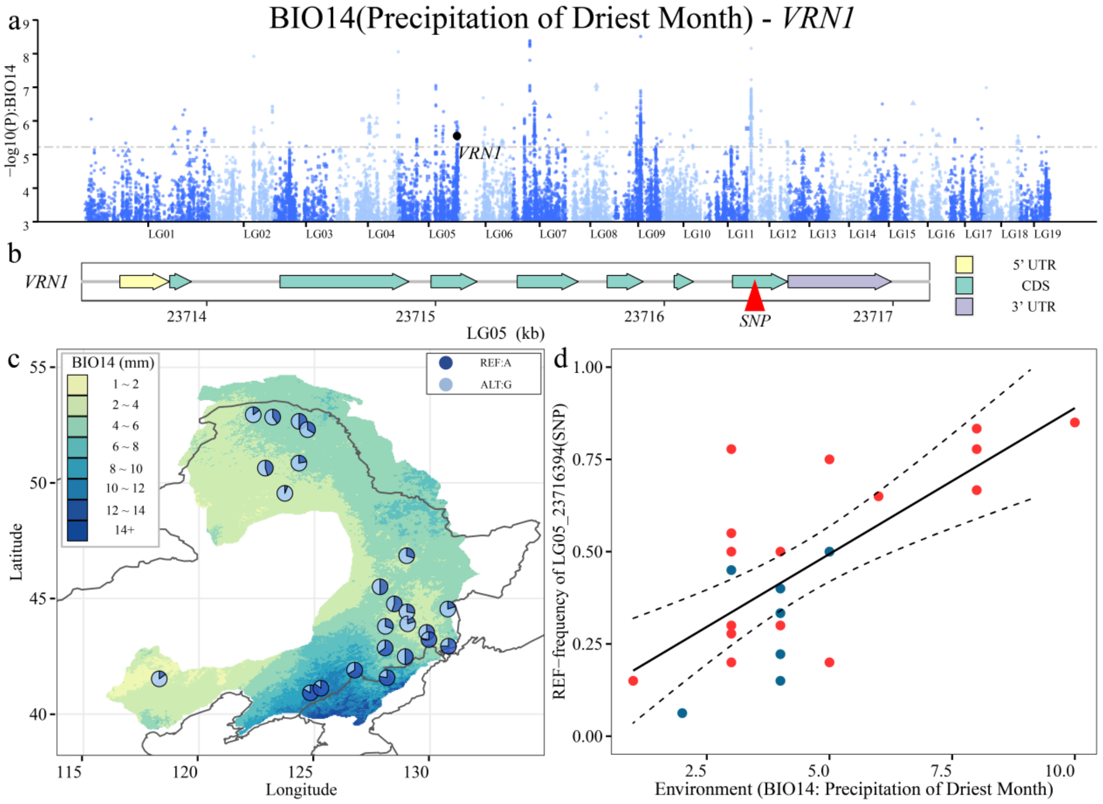

**Continuation Supplementary Fig. 12.**

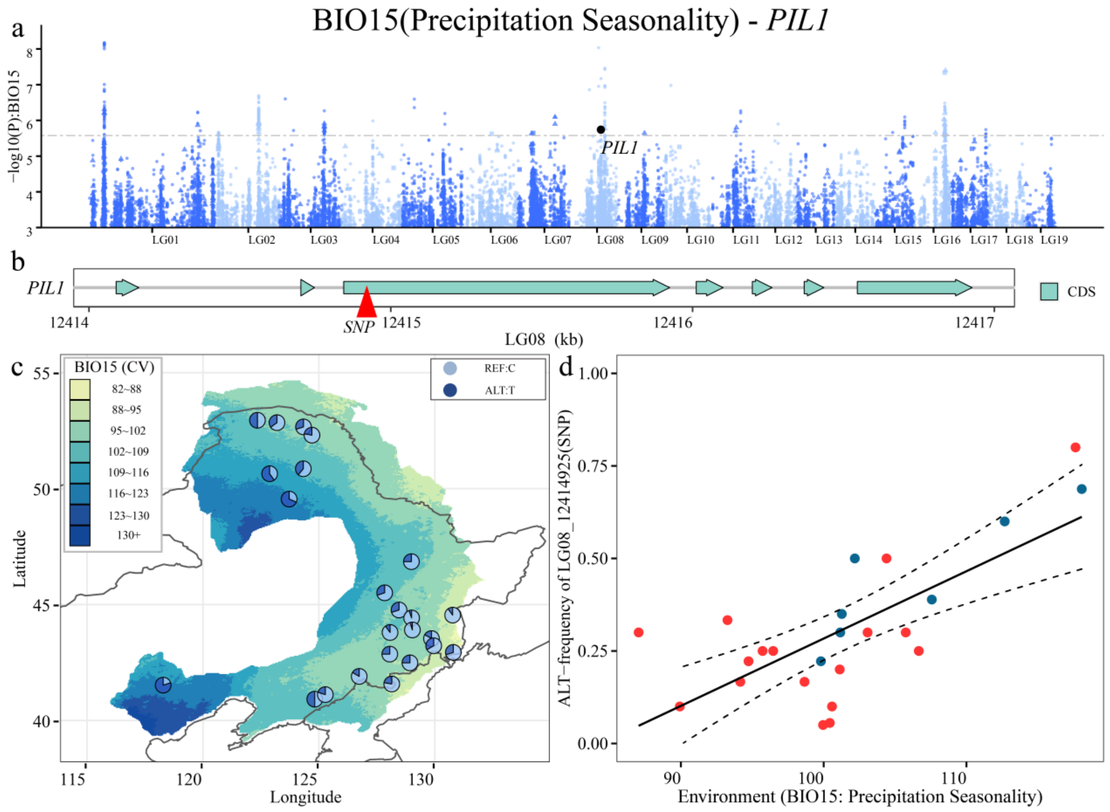

**Continuation Supplementary Fig. 12.**

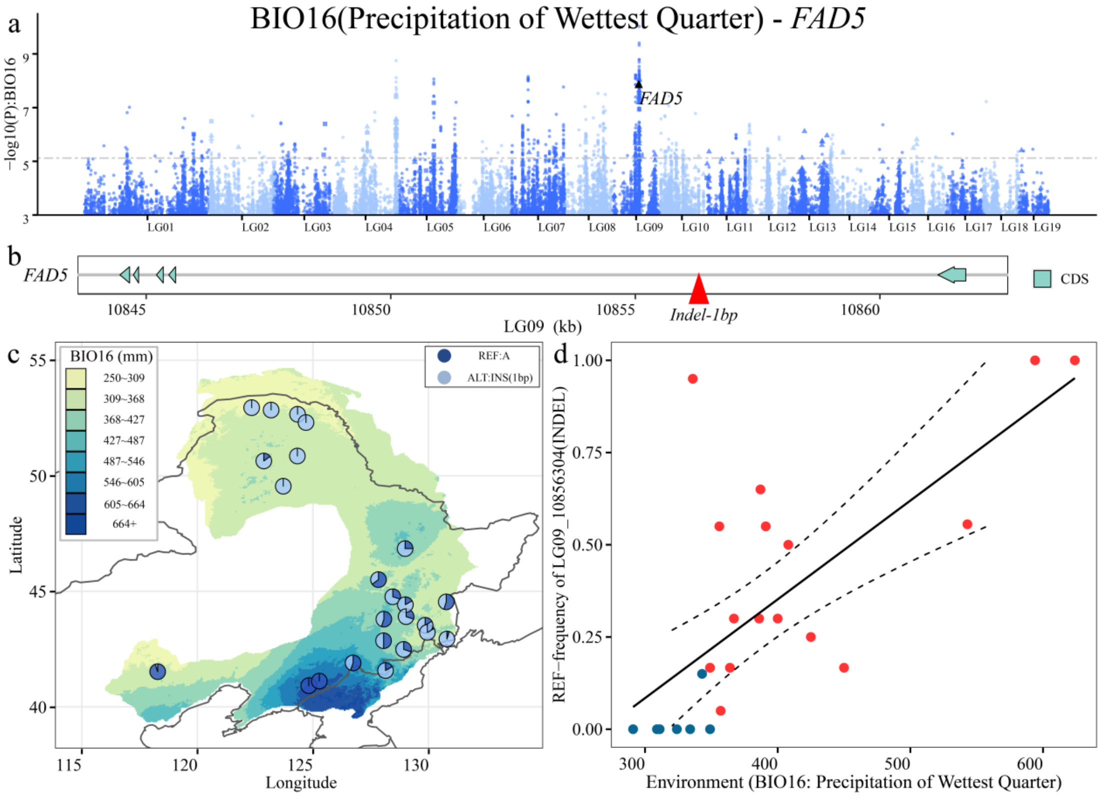

**Continuation Supplementary Fig. 12.**

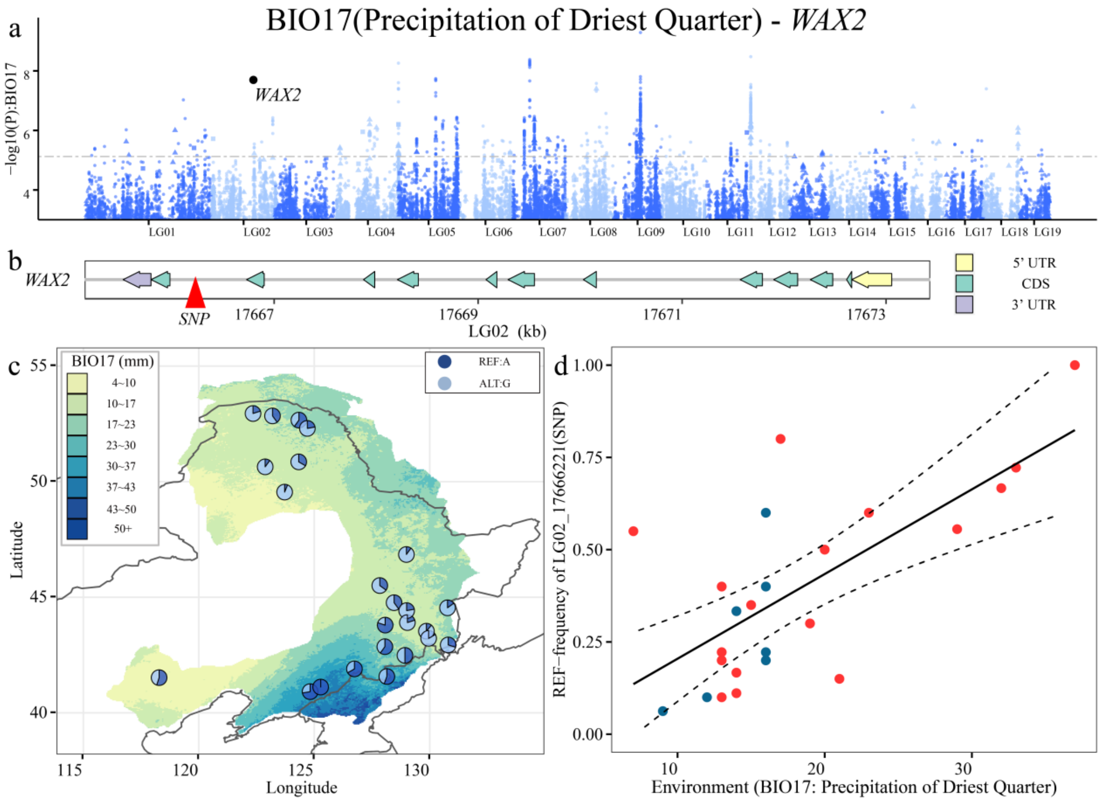

**Continuation Supplementary Fig. 12.**

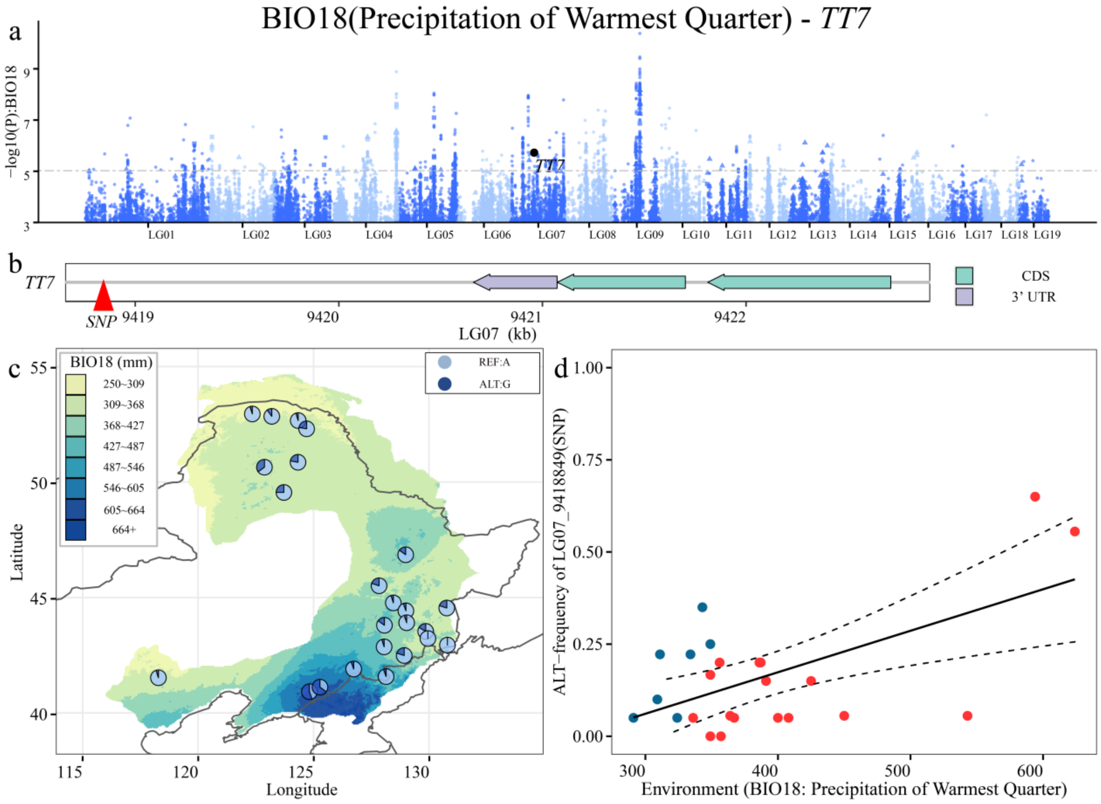

**Continuation Supplementary Fig. 12.**

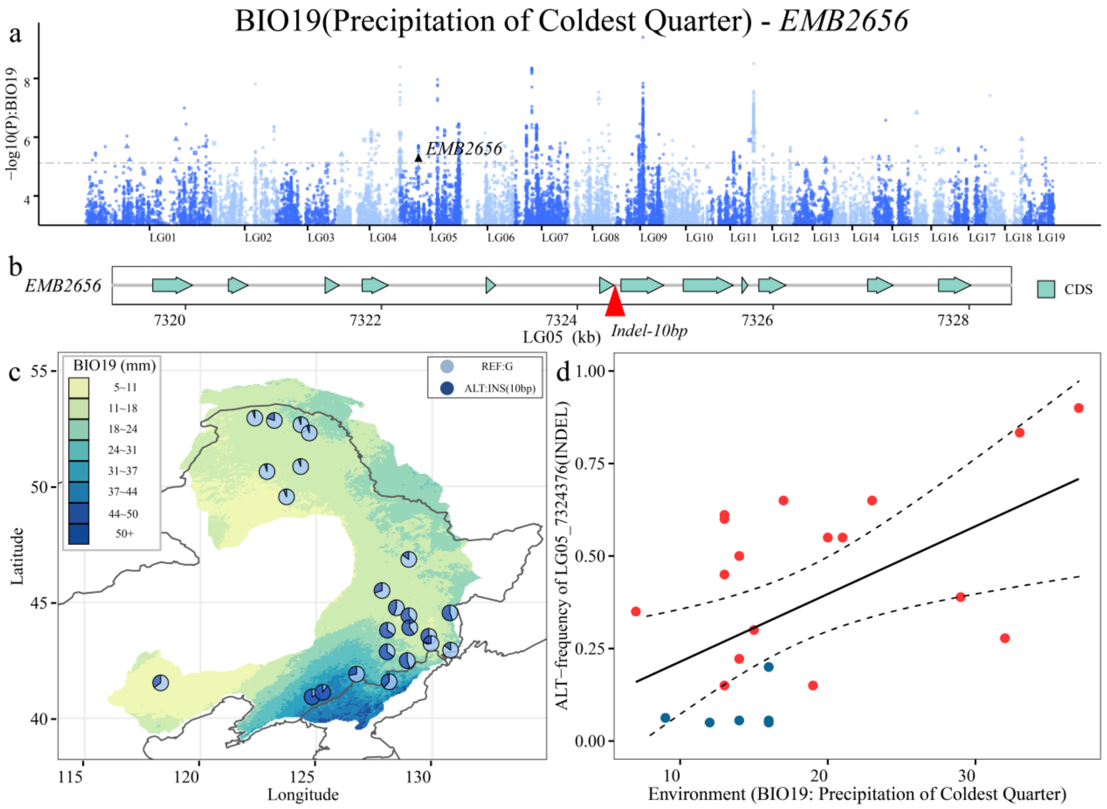

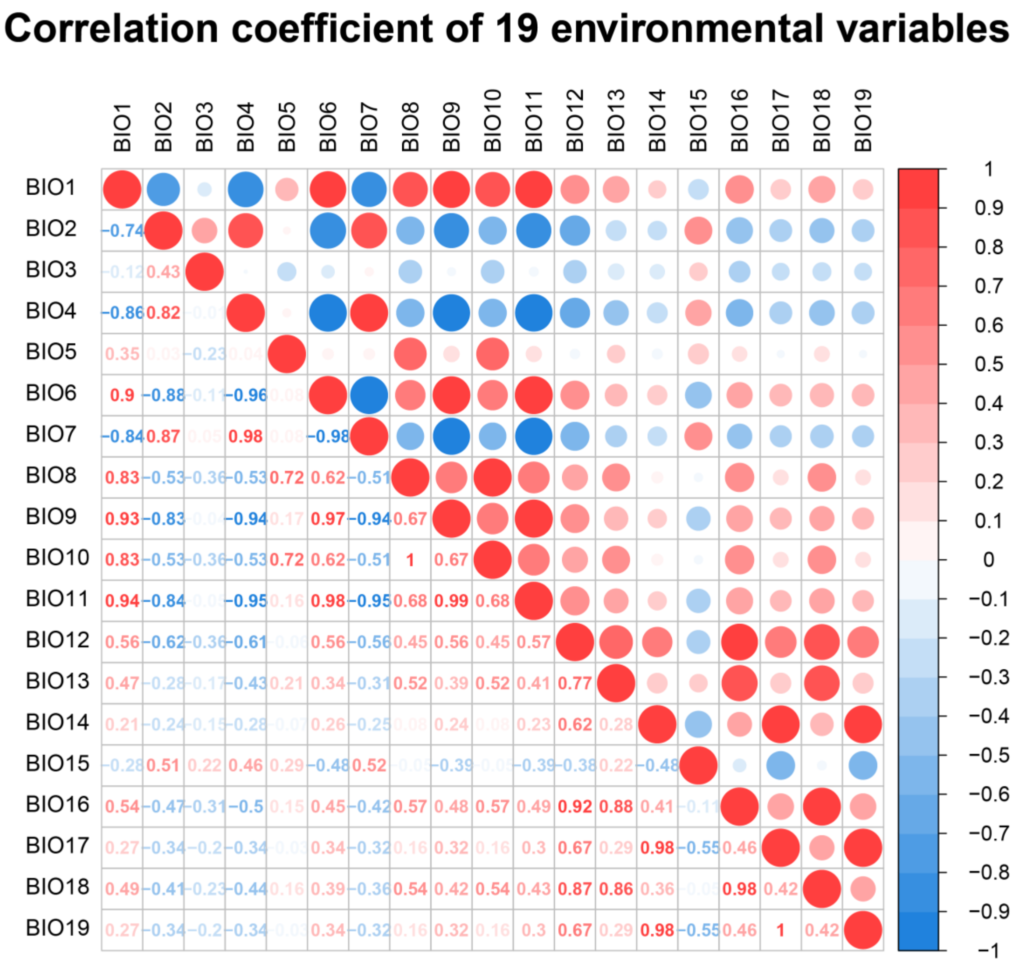


**Supplementary Fig. 13.** Spearman's correlation coefficient of 19 environmental variables.

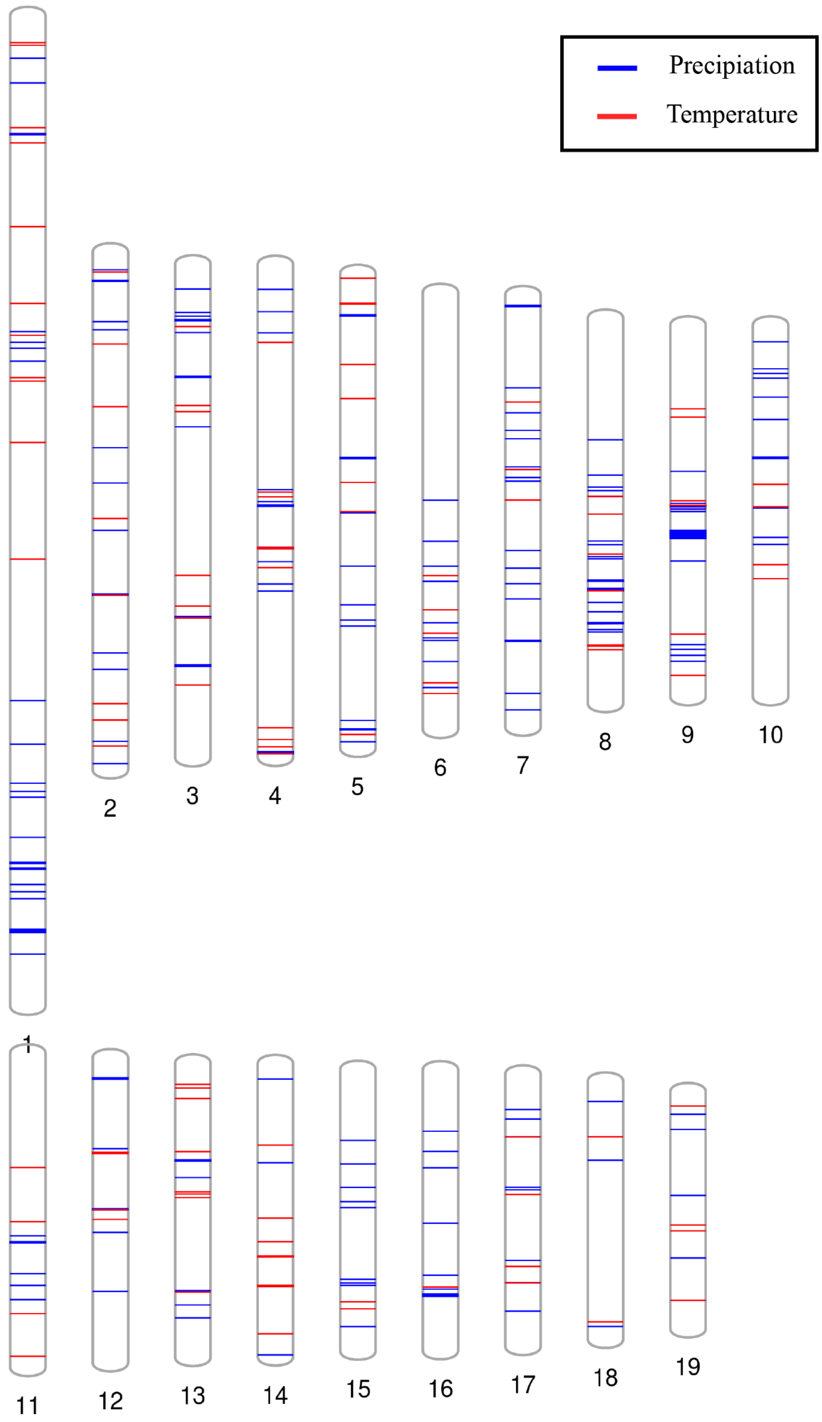


**Supplementary Fig. 14.** Mapping the location of the 1,779 core adaptive variants detected by both the approach of LFMM and RDA across the whole-genome. The blue and red bars indicate the adaptive variants associated with precipitation and temperature-related environmental variables, respectively.

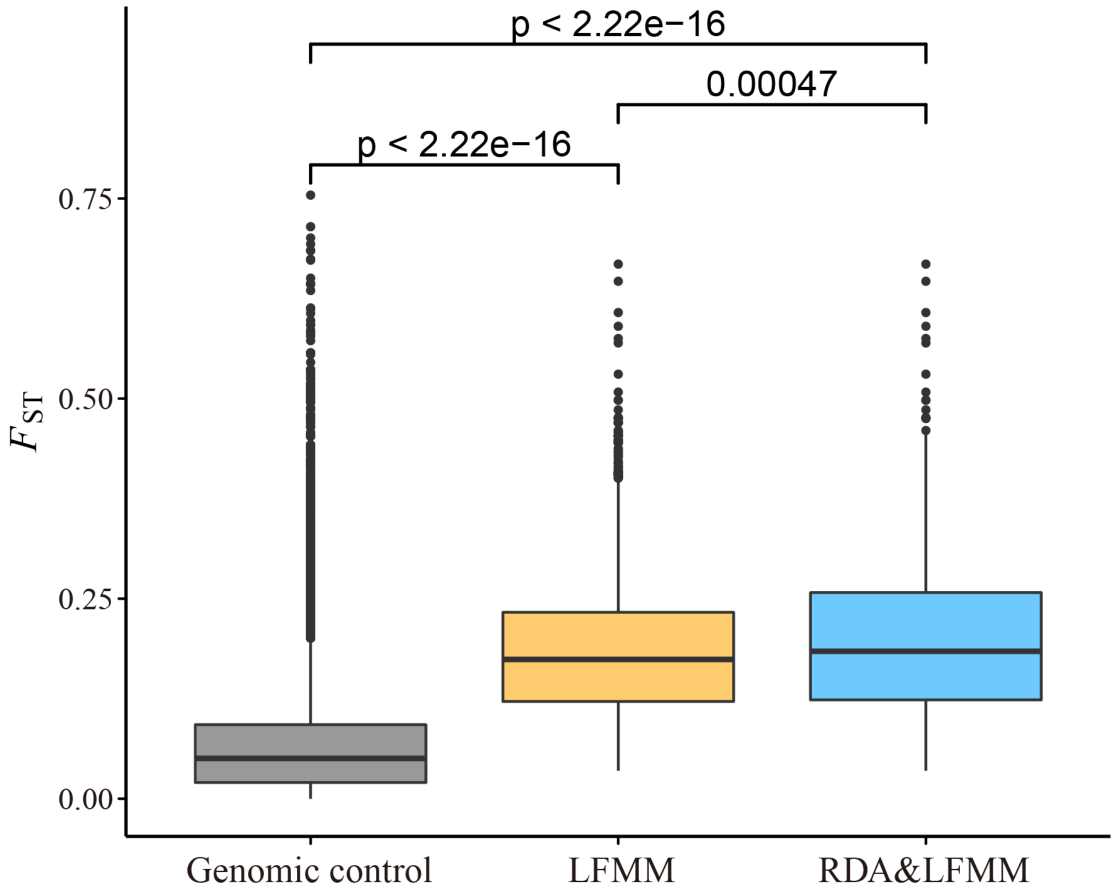


**Supplementary Fig. 15.** Comparison of genetic differentiation ($F_{ST}$) among pairs of populations calculated by using 100,000 random variants (genomic control), 3,435 environmental-associated variants detected by LFMM (LFMM) and 1,779 core adaptive variants detected by both LFMM and RDA (RDA&LFMM), respectively.

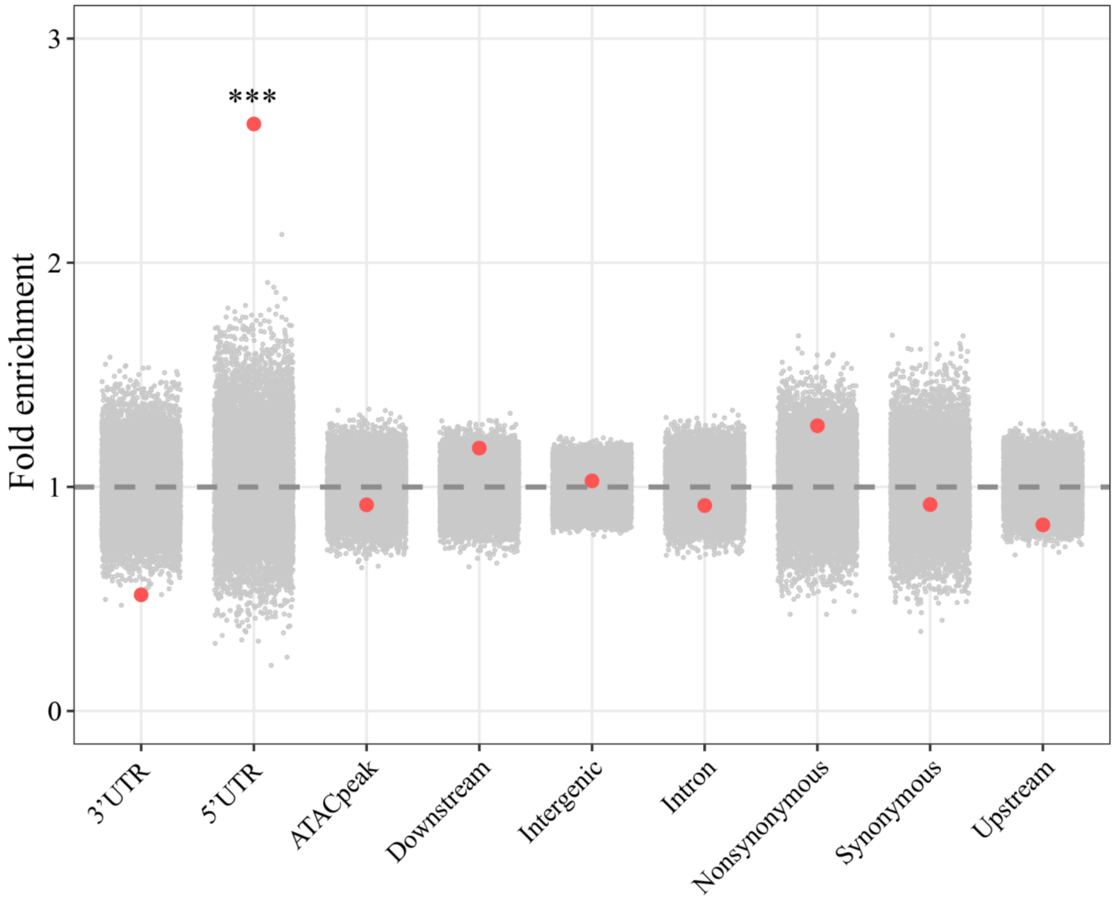


**Supplementary Fig. 16.** Enrichment of various functional categories in 1,779 core adaptive variants detected by both LFMM and RDA (red points). Grey dots show the distribution of results with 10,000 bootstrap replicates. The dashed line shows the expected enrichment under the null hypothesis of no enrichment. Enrichments that is significant relative to the bootstrap method are denoted by asterisks.

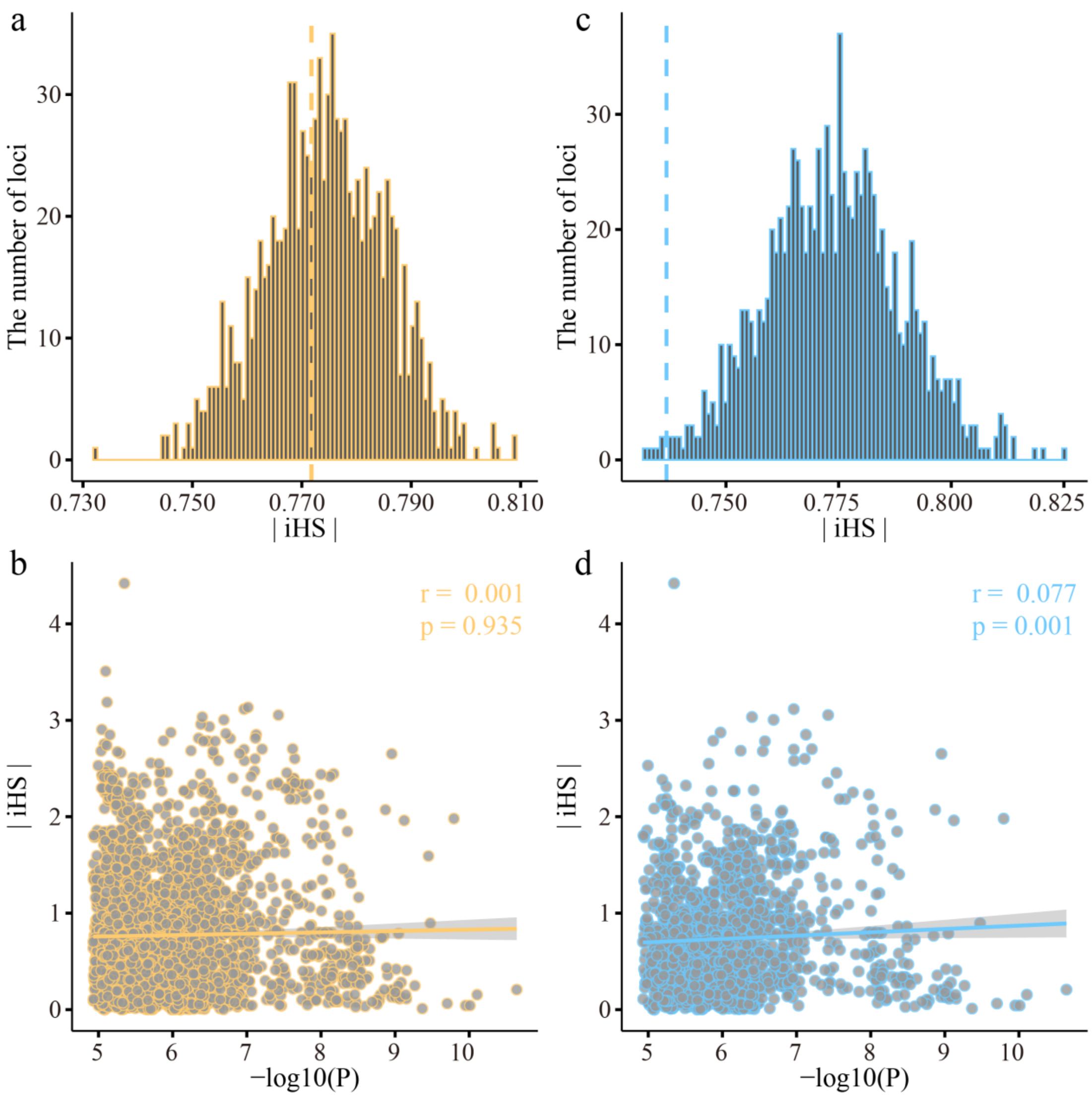


**Supplementary Fig. 17.** Weak selection signals for the environmental-associated variants. **a,c** The observed average values of |iHS| scores (dashed lines) relative to the genome-wide distribution (derived from 1000 bootstrap replicates) for the 3,435 variants detected by LFMM (a, yellow lines) and 1,779 adaptive variants detected by both LFMM and RDA methods (c, blue lines). **b,d** No significant relationship between signals of selection (|iHS|) and environmental associations (-log10P) for the 3,435 variants detected by LFMM (b, yellow dots) and 1,779 adaptive variants detected by both LFMM and RDA methods (d, blue dots).

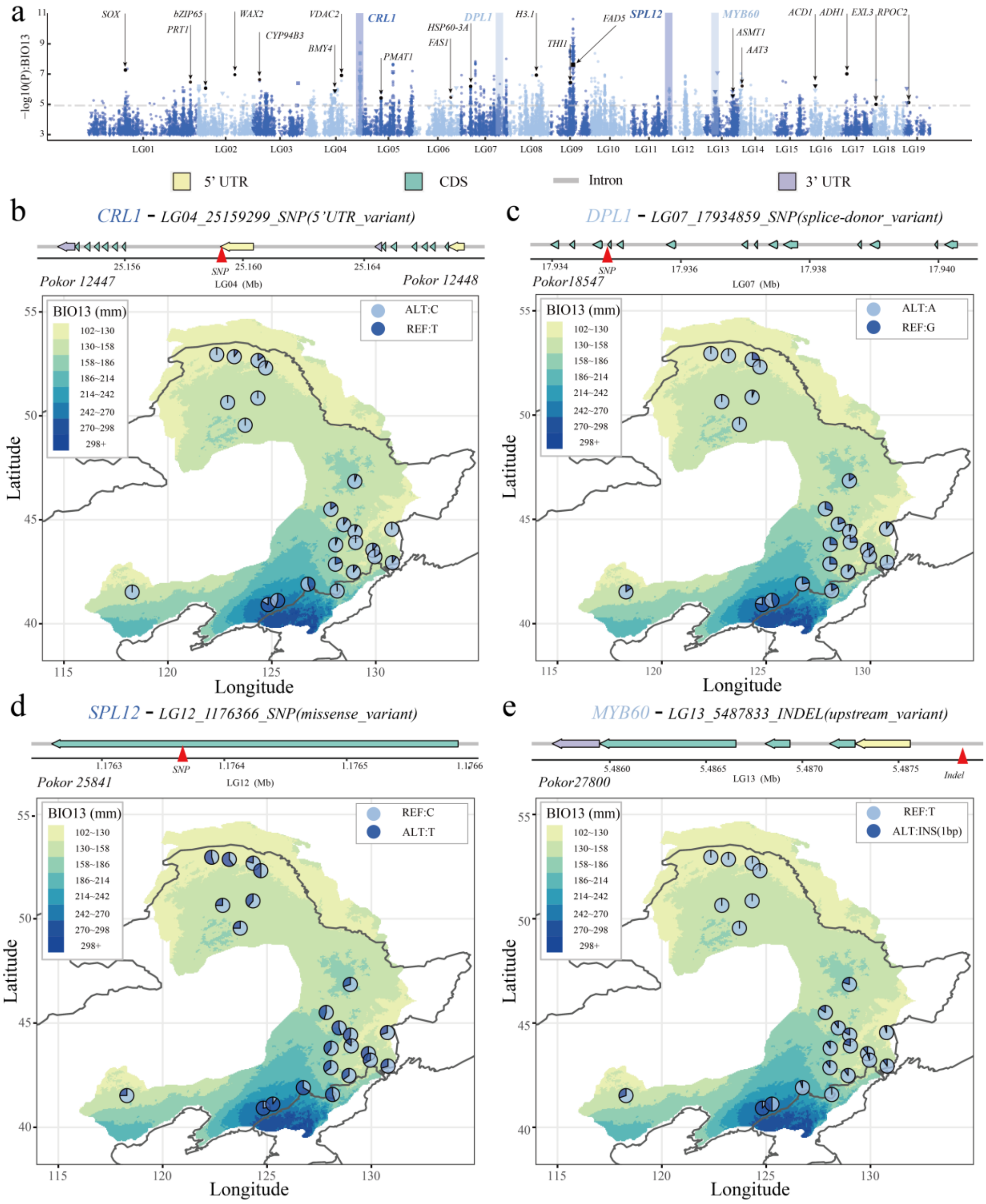

**Supplementary Fig. 18.** Display of some well-studied genes significantly related to BIO13 (precipitation of the wettest month). **a** Manhattan plot shows the genotype-environment association estimated with LFMM. SNPs, Indels and SVs are represented by points, triangles and squares, respectively. The grey dashed line represents q = 0.05. Colors distinguish different chromosomes. **b,c,d,e** The gene structure of selected genes, marked in **a** (upper panels). The distribution of allele frequencies of an example variation of the corresponding genes across 24 populations. Colors of raster on map represent the environmental variable under current scenario.

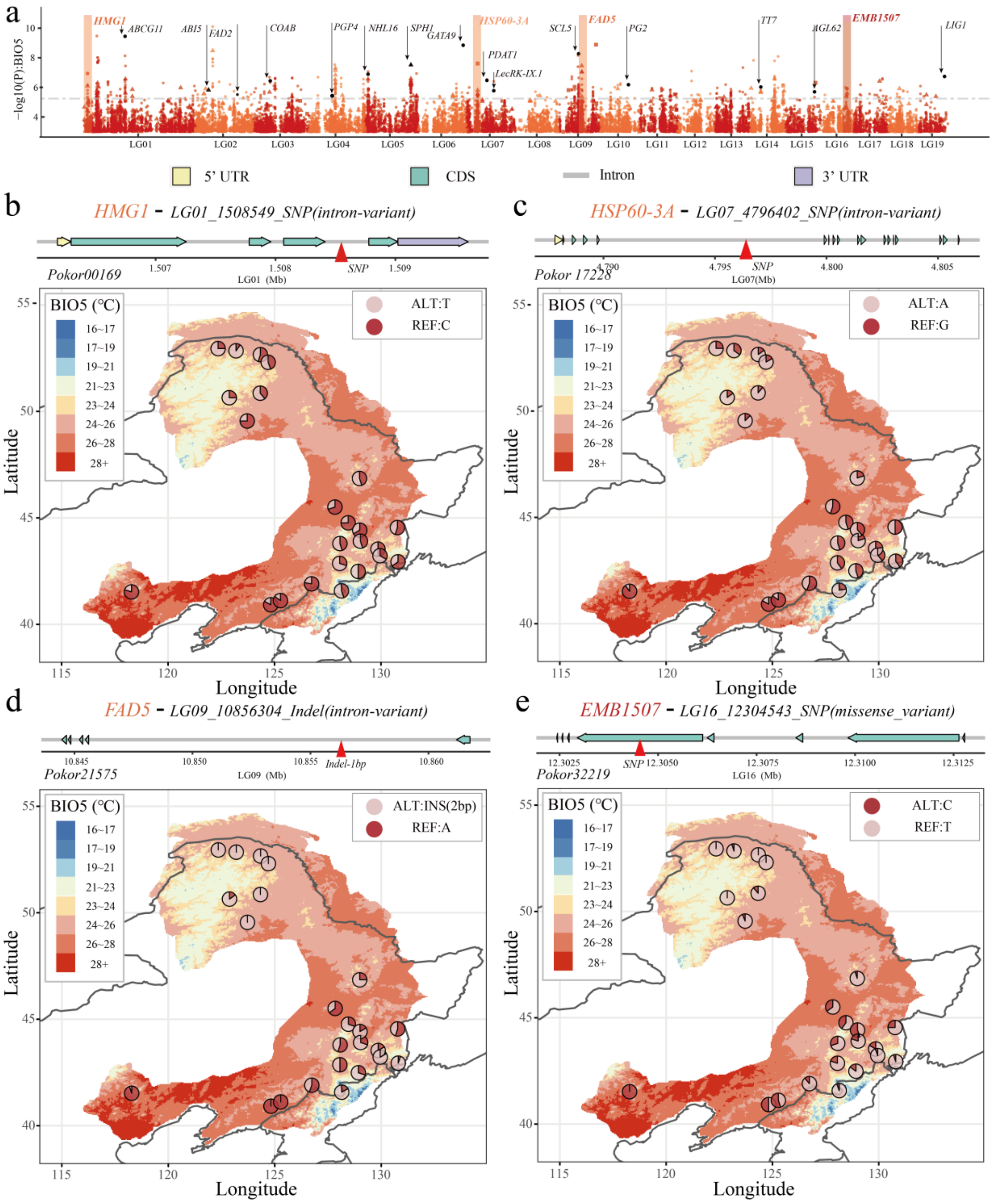


**Supplementary Fig. 19.** Display of some well-studied genes significantly related to BIO5 (maximum temperature of warmest month). **a** Manhattan plot shows the genotype-environment association estimated with LFMM. SNPs, Indels and SVs are represented by points, triangles and squares, respectively. The grey dashed line represents q = 0.05. Colors distinguish different chromosomes. **b,c,d,e** The gene structure of selected genes, marked in **a** (upper panels). The distribution of allele frequencies of an example variation of corresponding gene across 24 populations. Colors of raster on map represent the environmental variable under current scenario.

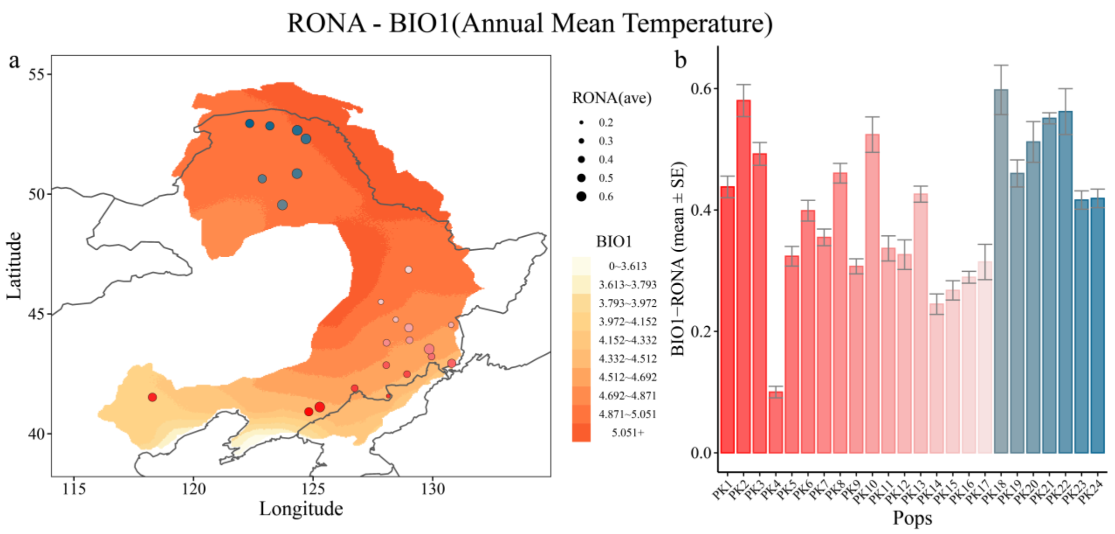


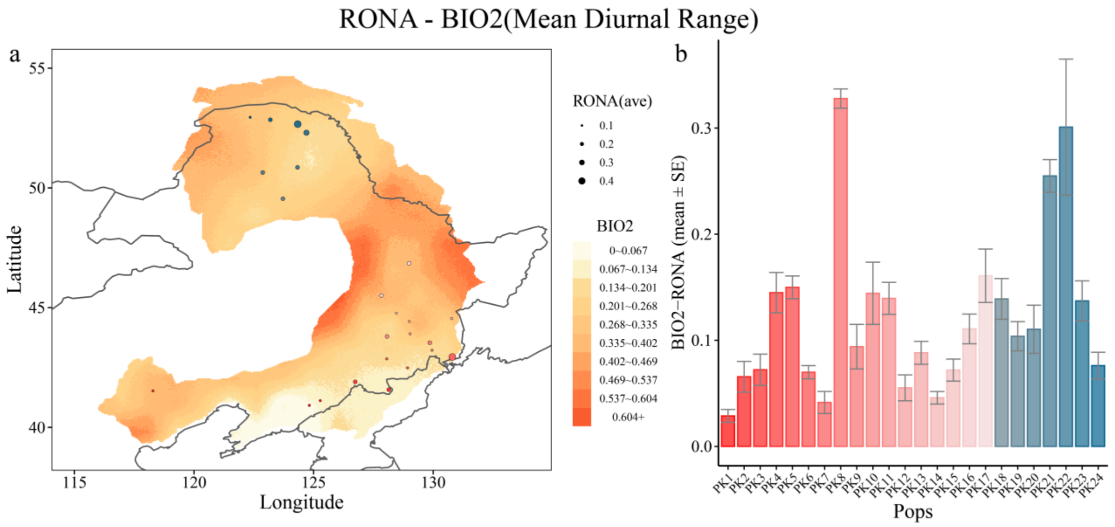


**Supplementary Fig. 20.** Risk of nonadaptedness (RONA) for the 19 environmental variables (BIO1-BIO19). **a** Solid circle with different colors (red and blue represents south and north groups of populations) on map reflects different natural populations, which also correspond to the populations shown in (b). The circle size represents the average RONA values (weighted means by $R^2$ value). The raster of map represents the magnitude of climate change (absolute value between SSP370 in 2080 and current). Areas with darker red or blue indicate more dramatic predicted climate change for temperature-related and precipitation-related variables, respectively. **b** Bars represent weighted means of RONA, and lines represent standard errors of RONA for each population.

**Continuation Supplementary Fig. 20.**

RONA - BIO3(Isothermality (BIO2/BIO7))

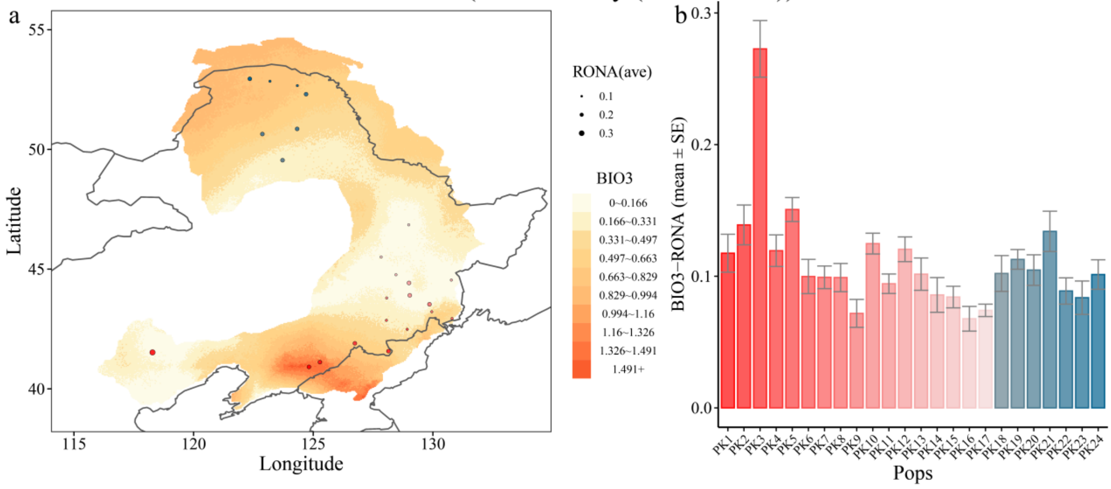


RONA - BIO4(Temperature Seasonality)

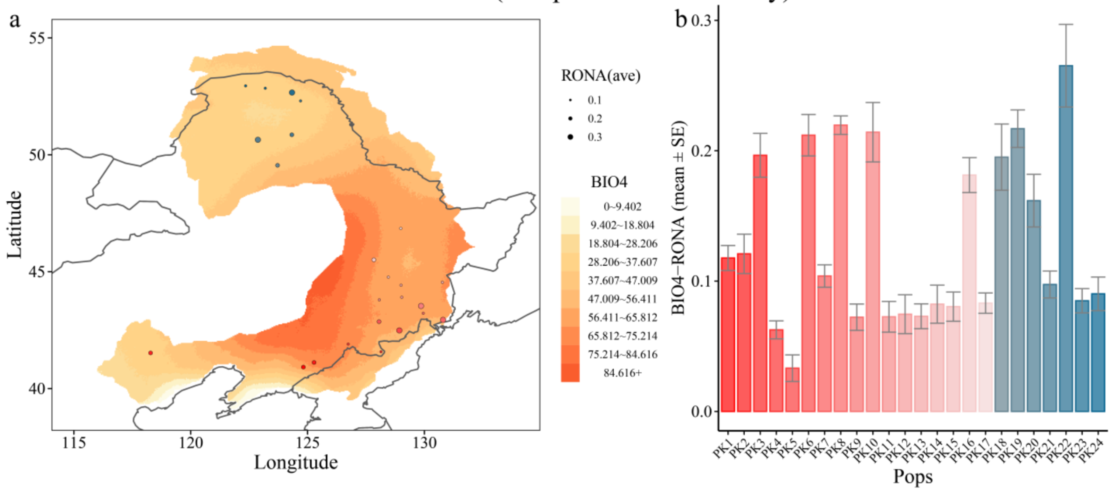


RONA - BIO5(Maximum Temperature of Warmest Month)

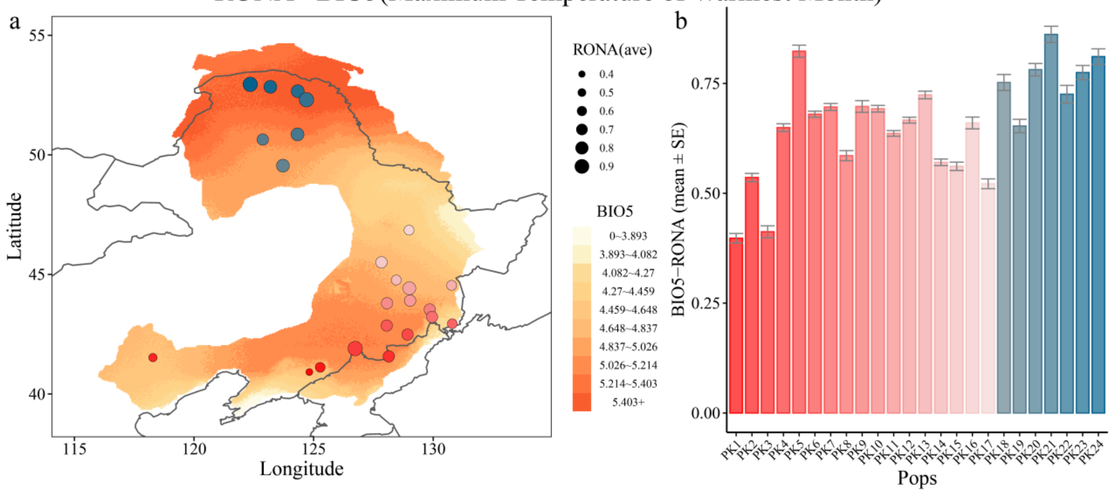

### Continuation Supplementary Fig. 20.

RONA - BIO6(Minimum Temperature of Coldest Month)

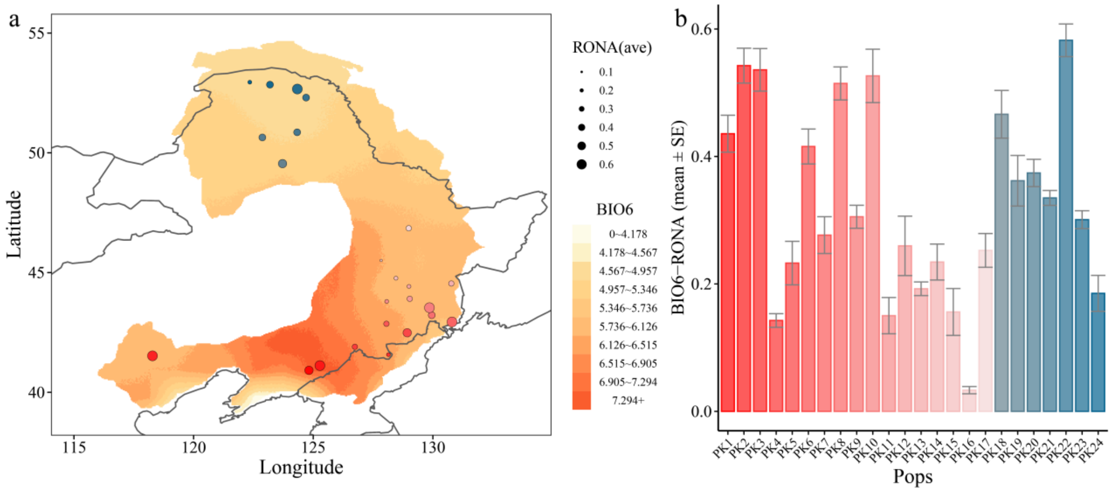


RONA - BIO7(Temperature Annual Range (BIO5-BIO6))

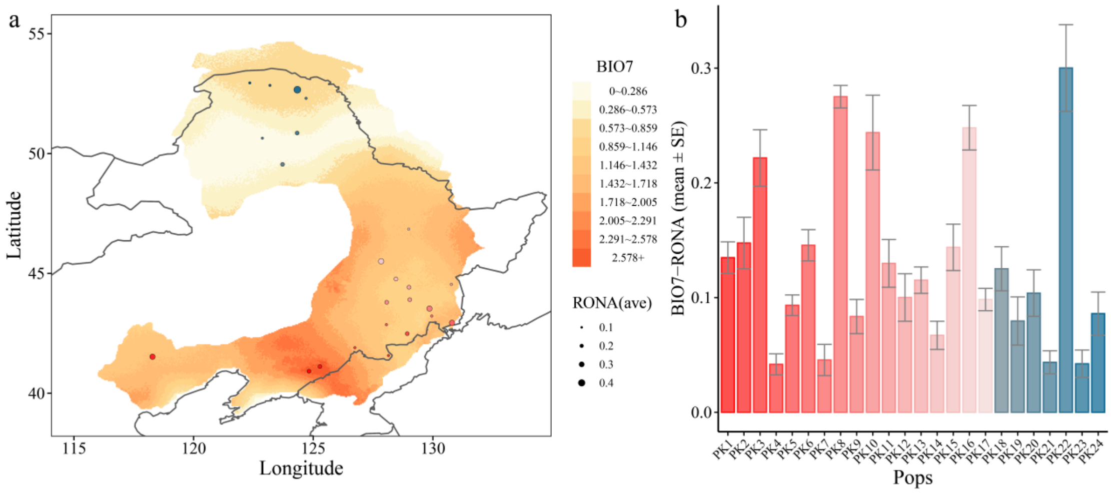


RONA - BIO8(Mean Temperature of Wettest Quarter)

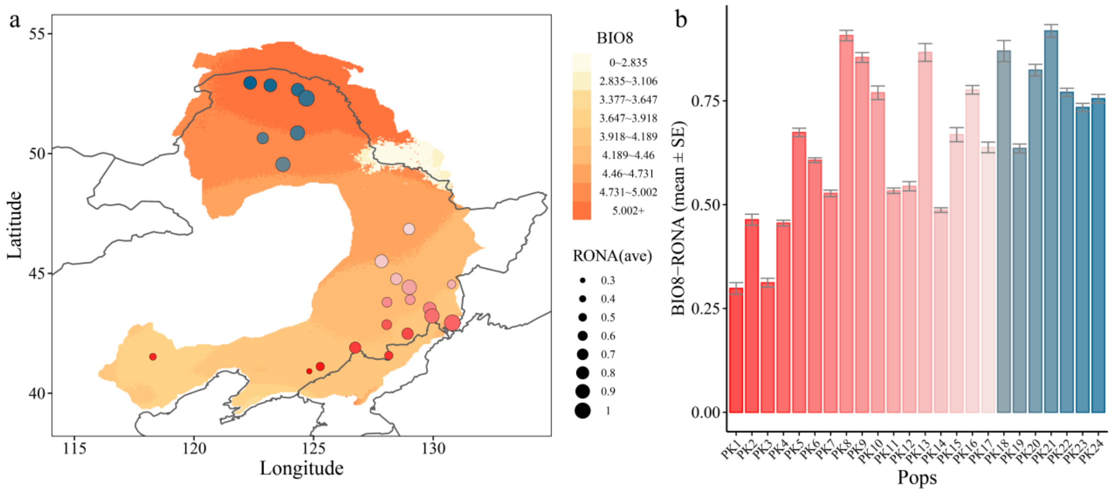

### Continuation Supplementary Fig. 20.

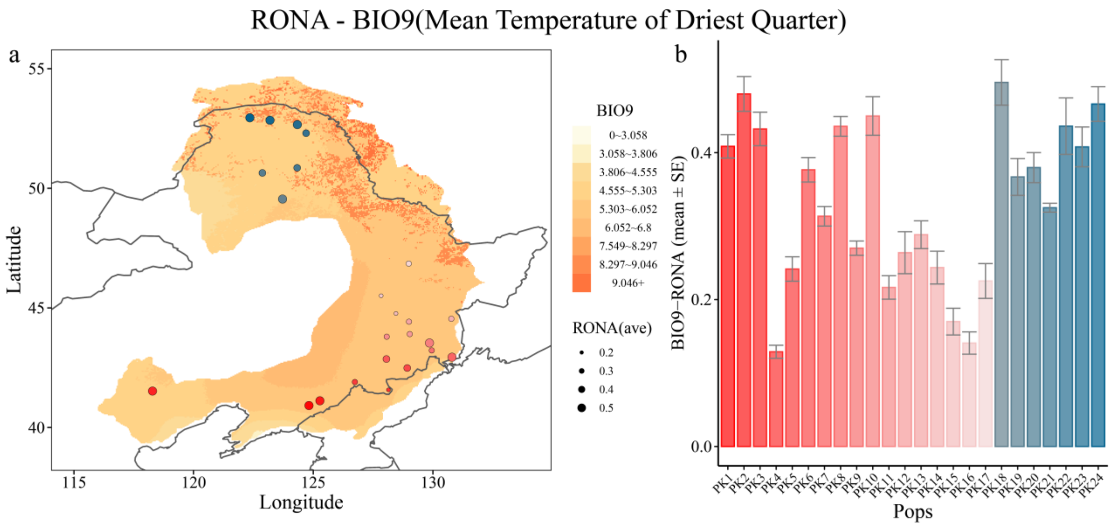


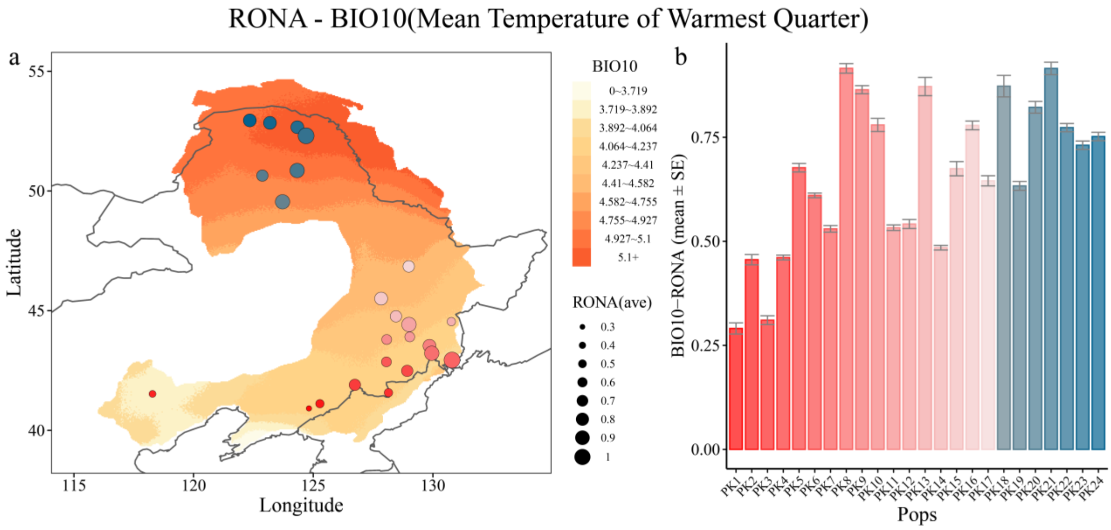


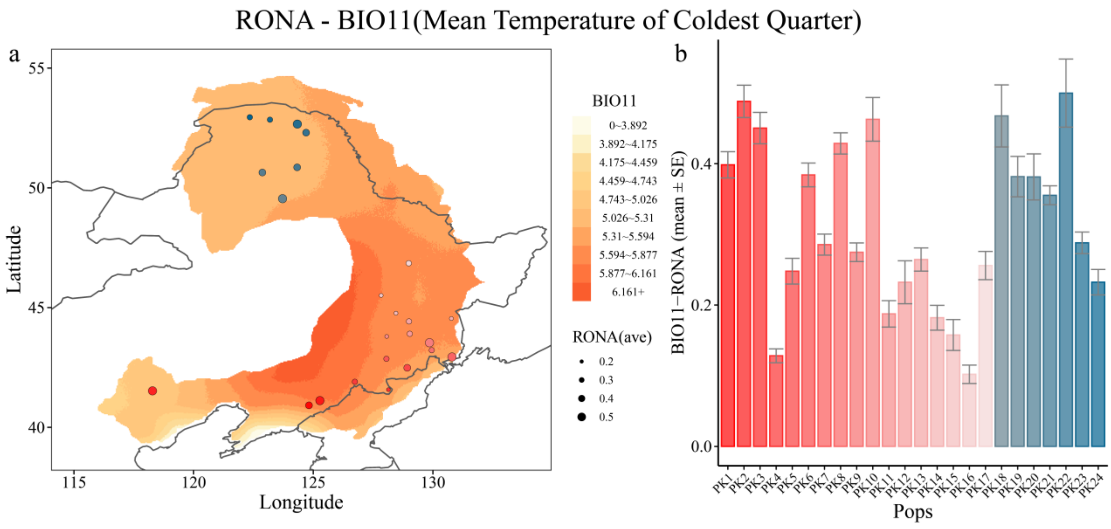

**Continuation Supplementary Fig. 20.**

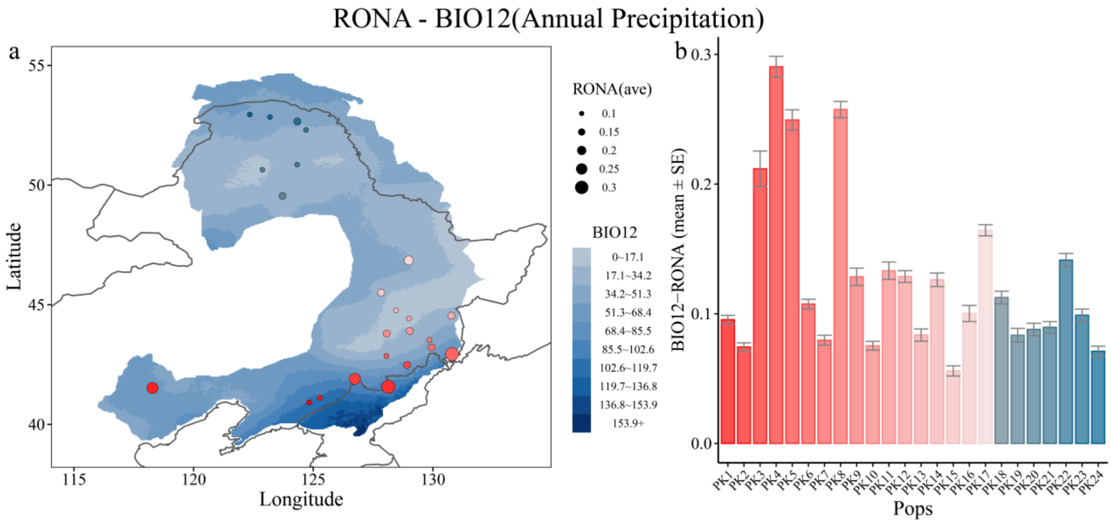


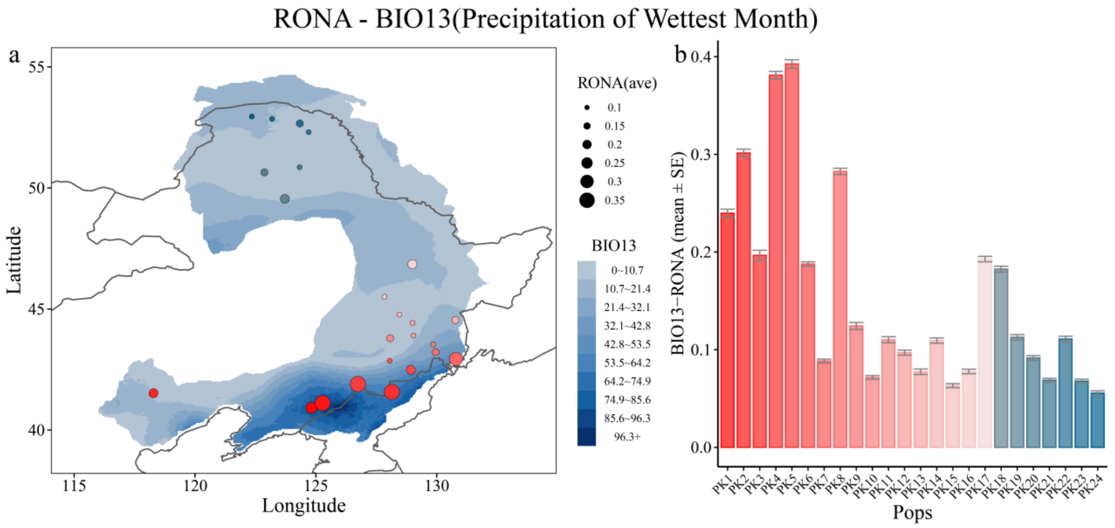


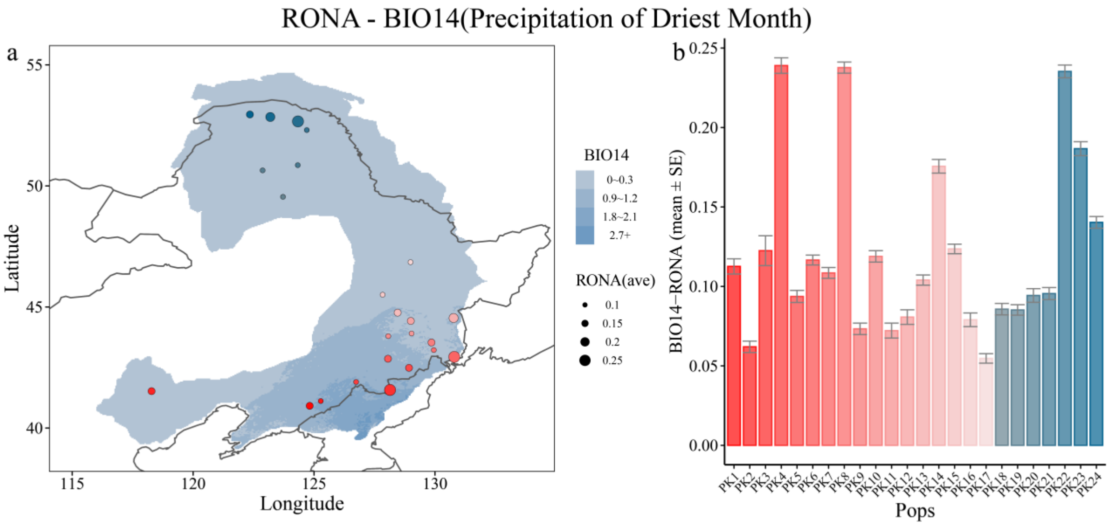

### Continuation Supplementary Fig. 20.

RONA - BIO15(Precipitation Seasonality)

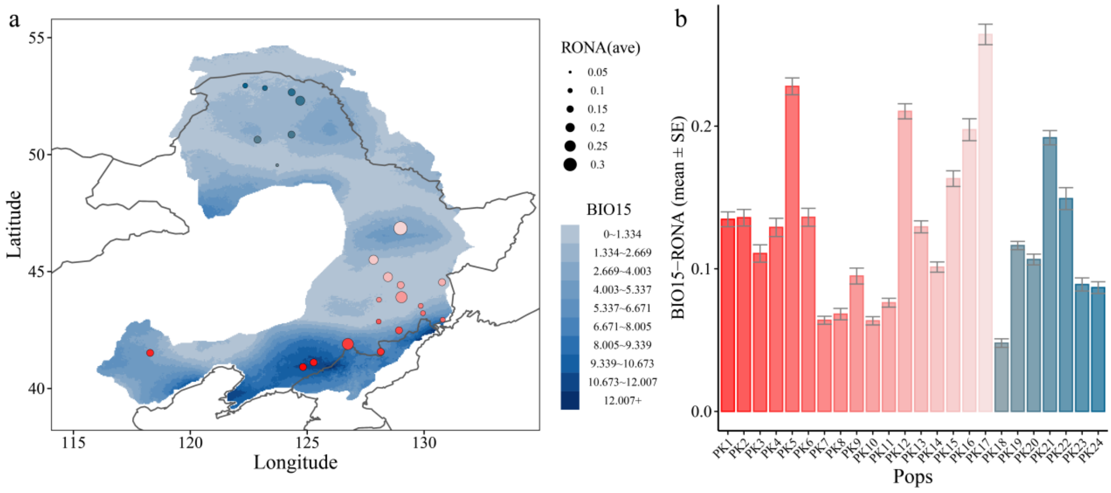


RONA - BIO16(Precipitation of Wettest Quarter)

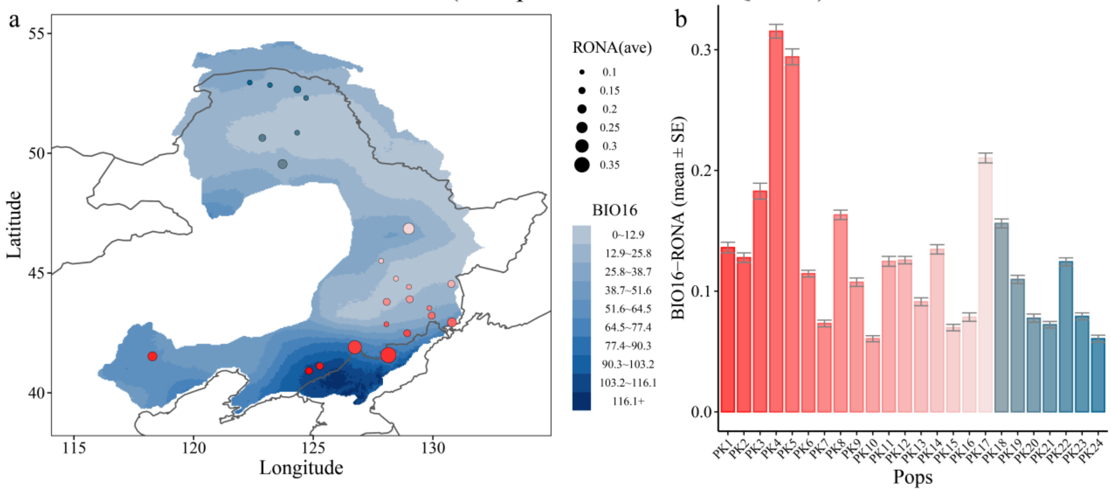


RONA - BIO17(Precipitation of Driest Quarter)

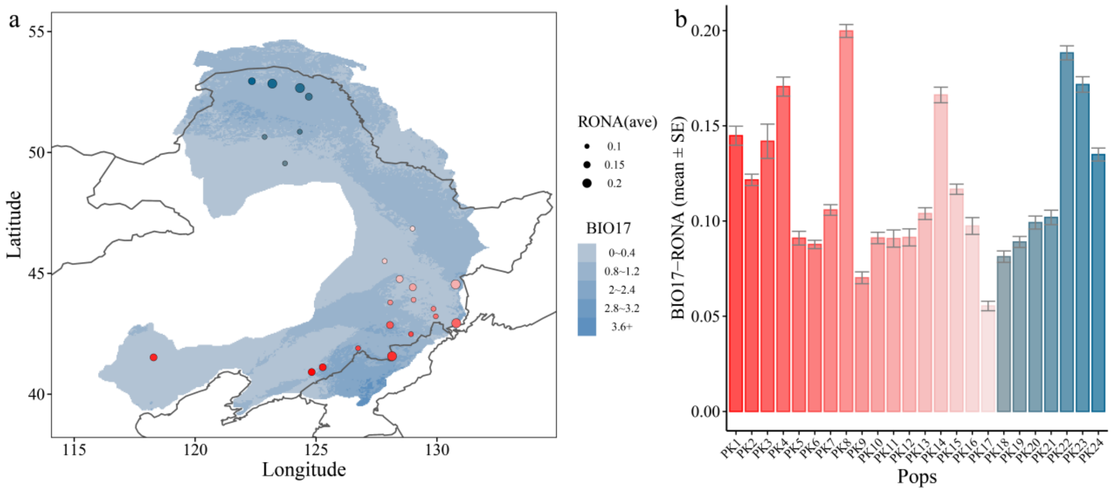

**Continuation Supplementary Fig. 20.**

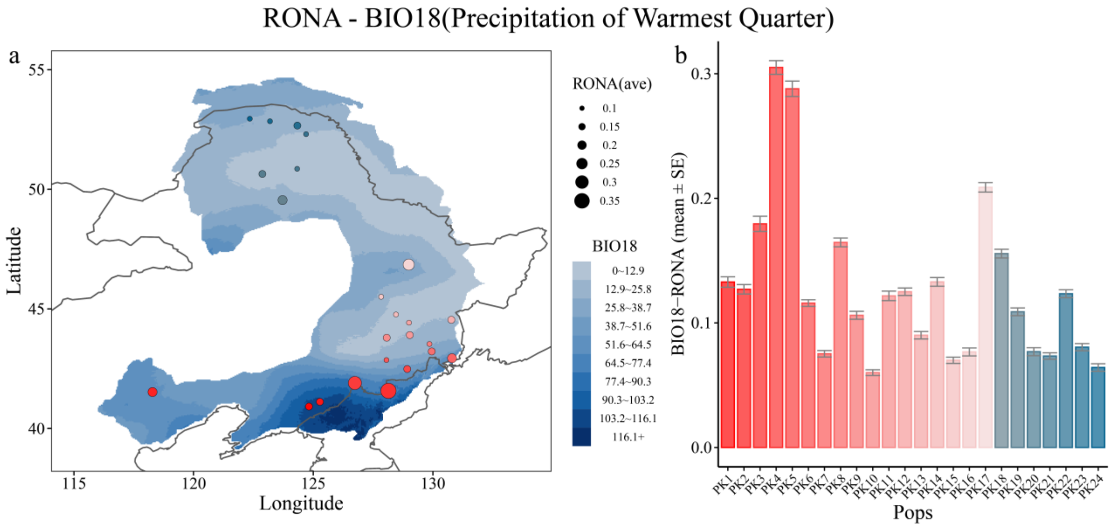


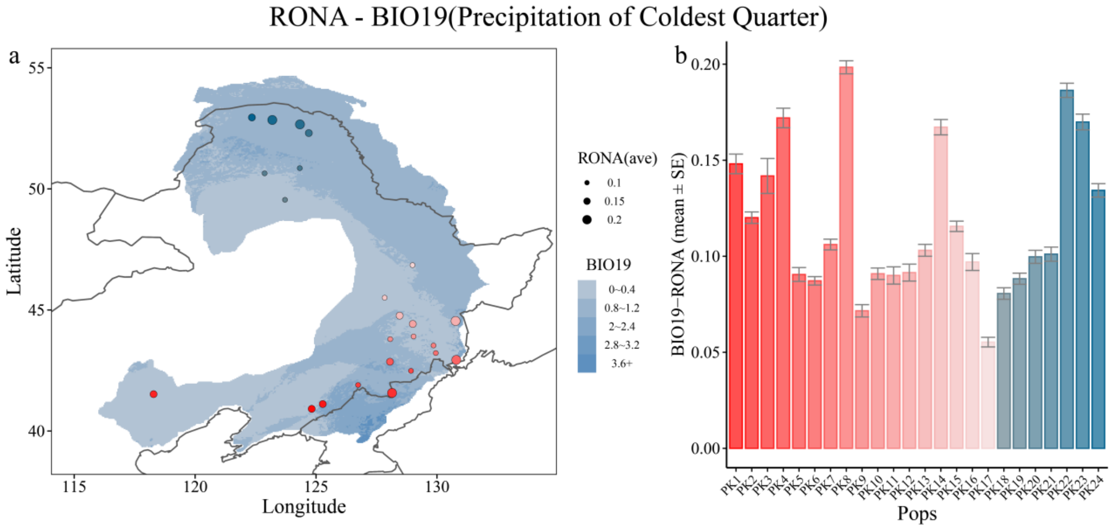

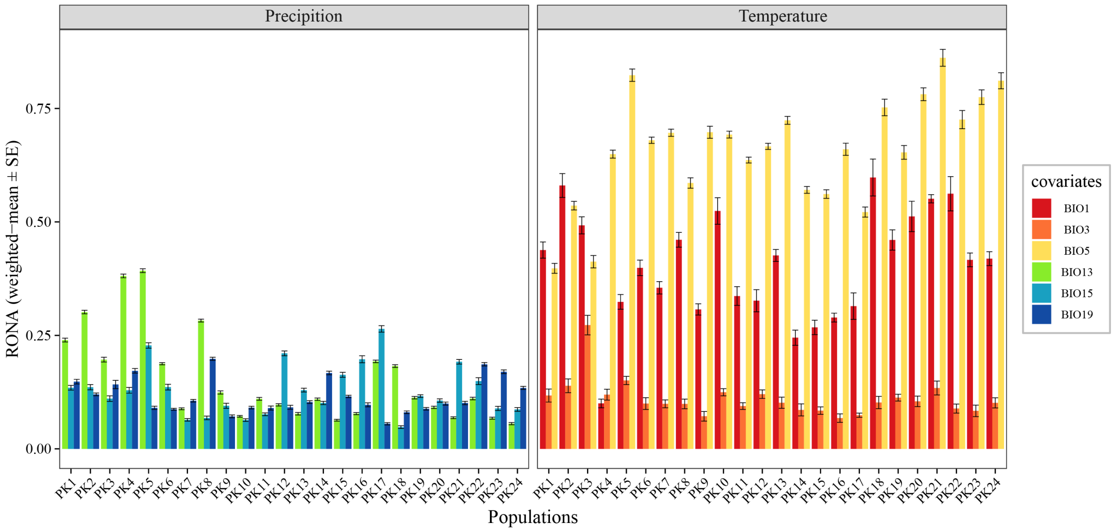


**Supplementary Fig. 21.** Risk of nonadaptedness (RONA) for six uncorrelated precipitation and temperature-related variables under SSP370 in 2080. Bars represent weighted averages of RONA, and lines represent standard error of RONA for each population.

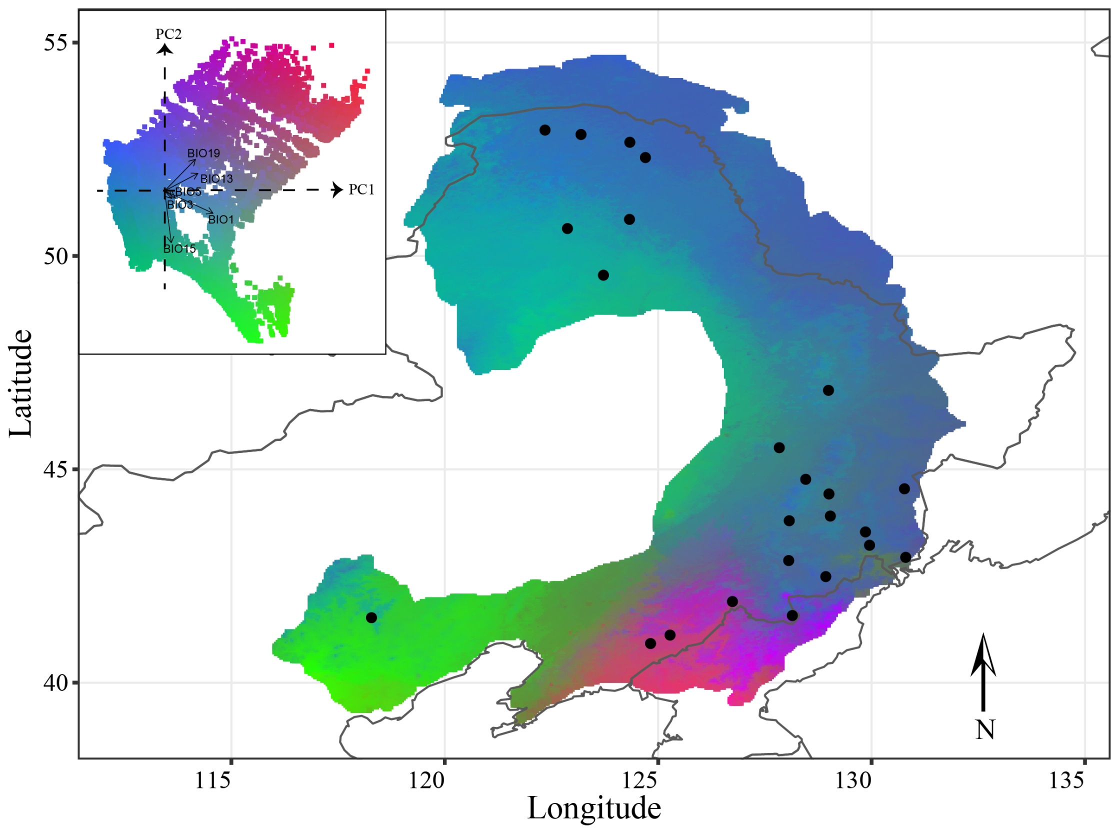


**Supplementary Fig. 22.** Gradient forest-transformed climate variables show climate adaptation across the distribution of *P. koreana*. Inset: Colors are based upon modelled genetic-environment associations from the 60,000 random points generated in the distribution range. Arrows show the loadings of the six uncorrelated environmental variables.

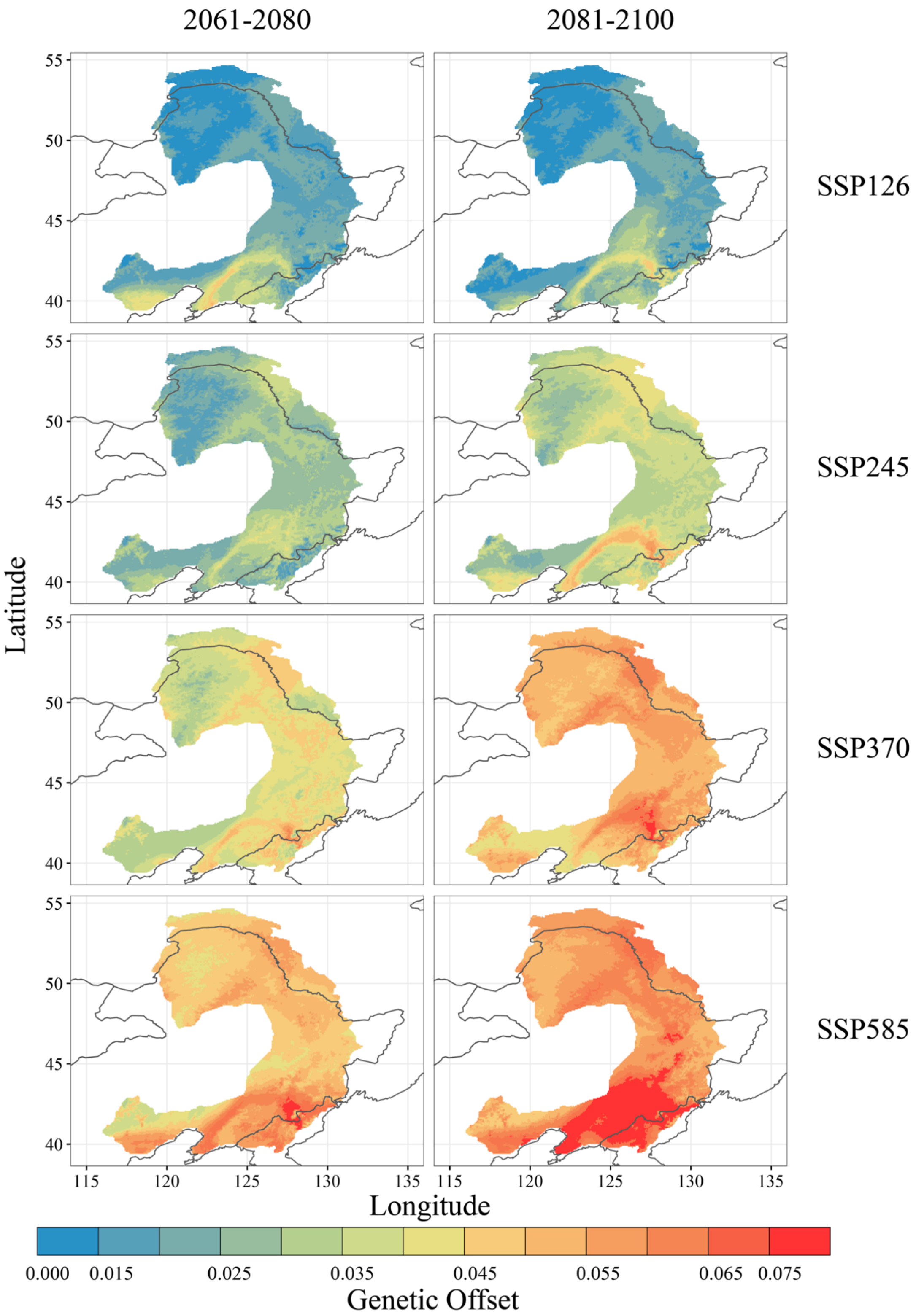


**Supplementary Fig. 23.** Map of genetic offset based on the estimates using 19 climate variables across the natural distribution of *P. koreana* for eight different scenarios of Shared Socio-economic Pathways (SSPs) at 2061-2080 and 2081-2100. The color scale from blue to red refers to the increasing of genetic offset.

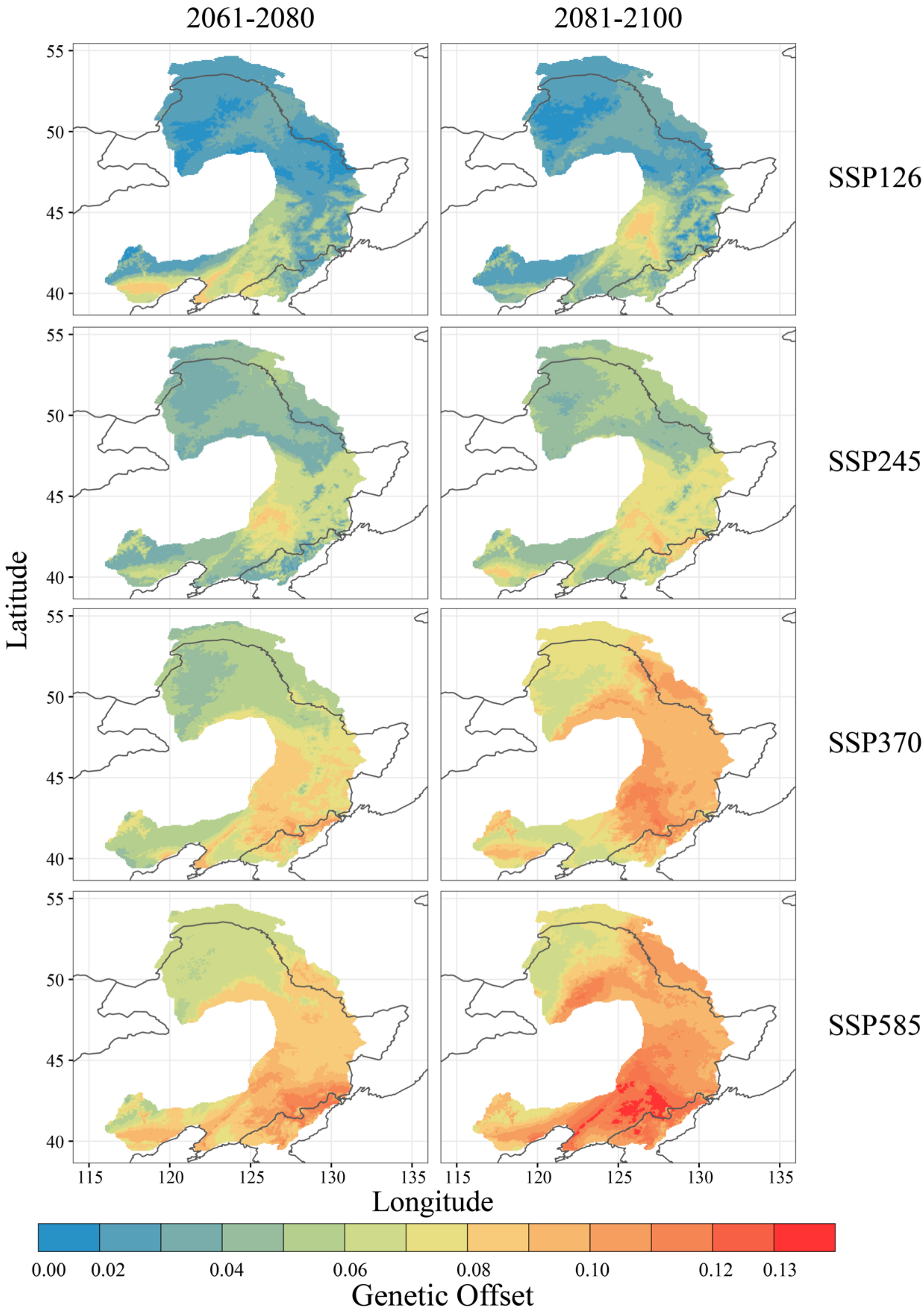


**Supplementary Fig. 24.** Map of genetic offset based on the estimates using six uncorrelated climate variables across the natural distribution of *P. koreana* for eight different scenarios of Shared Socio-economic Pathways (SSPs) at 2061-2080 and 2081-2100. The color scale from blue to red refers to the increasing of genetic offset.

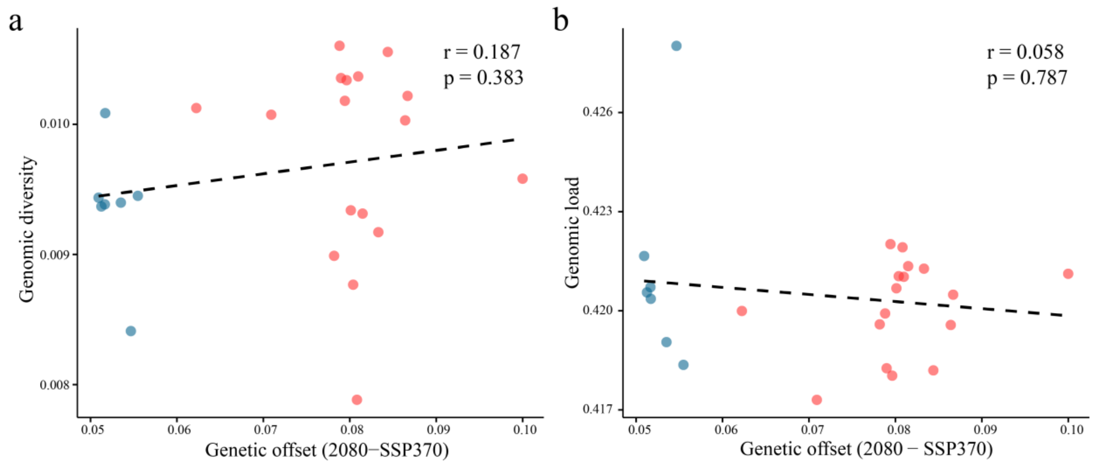


**Supplementary Fig. 25. a** The relationship (Pearson's test) between the genomic diversity (y axis) and prediction of genetic offset (x axis) under 2080-SSP370 scenario across the 24 natural populations of *P. koreana*. **b** The relationship (Pearson's test) between the estimated genomic load (y axis, measured as the proportion of 0-fold non-synonymous and 4-fold synonymous SNPs) and prediction of genetic offset (x axis) under 2080-SSP370 scenario.

**Supplementary Table 1.** Statistics of sequencing reads.

| Platform | Total reads | Total bases | Coverage (x) |
|---|---|---|---|
| Nanopore | 1,938,650 | 42,419,739,676 | 105.68 |
| Illumina | 198,806,712 | 2,982,1006,800 | 74.29 |
| Hi-C | 366,626,810 | 54,994,021,500 | 137.00 |

**Supplementary Table 2.** Illumina sequencing data statistics.

| **Sequencing data statistics** | |
|---|---|
| Total reads | 198,806,712 |
| Total bases | 29,821,006,800 |
| Clean reads | 196,862,480 |
| Clean bases | 27,377,255,469 |
| Q20 rate (%) | 96.65 |
| Q30 rate (%) | 90.79 |
| GC (%) | 35.12 |

**Supplementary Table 3.** Nanopore sequencing data statistics.

| **Sequencing data statistics** | |
|---|---|
| Reads data (bp) | 42,419,739,676 |
| Reads number | 1,938,650 |
| Reads mean length (bp) | 21,881 |
| Reads max length (bp) | 214,618 |
| Reads N50 (bp) | 28,921 |
| >10kb (%) | 80.08 |
| >20kb (%) | 47.70 |
| >40kb (%) | 11.06 |

**Supplementary Table 4.** Mapping summary of Hi-C data.

| **Hi-C data** | |
|---|---|
| Raw Paired-end Reads | 366,626,810 |
| Clean Paired-end Reads | 361,786,732 (98.68%) |
| Clean Base(bp) | 54,223,719,643 |
| Clean Q30 Base Rate (%) | 92.00 |
| Clean Pair-end Reads | 180,893,366 |
| Unmapped Paired-end Reads | 4,371,524 (2.42%) |
| Paired-end Reads with Singleton | 32,010,746 (17.70%) |
| Multi Mapped Paired-end Reads | 30,347,997 (16.78%) |
| Unique Mapped Paired-end Reads | 114,163,099 (63.11%) |
| Valid reads of unique mapping reads (%) | 86.05 |
| Valid reads of clean reads (%) | 54.31 |

**Supplementary Table 5.** Scaffolding of contigs based on Hi-C data.

| Pseudo-chromosome | Size (Contig number) |
|---|---|
| **LG01** | 51,125,891 (15) |
| **LG02** | 26,717,773 (7) |
| **LG03** | 25,475,328 (5) |
| **LG04** | 25,429,882 (7) |
| **LG05** | 24,496,347 (10) |
| **LG06** | 22,536,972 (5) |
| **LG07** | 22,306,471 (9) |
| **LG08** | 19,875,049 (6) |
| **LG09** | 19,170,182 (8) |
| **LG10** | 19,169,738 (6) |
| **LG11** | 17,803,069 (11) |
| **LG12** | 17,266,452 (8) |
| **LG13** | 16,661,543 (5) |
| **LG14** | 16,606,995 (5) |
| **LG15** | 15,942,094 (7) |
| **LG16** | 15,894,631 (8) |
| **LG17** | 15,420,218 (3) |
| **LG18** | 14,612,902 (5) |
| **LG19** | 13,424,669 (3) |
| **Total** | 399,936,206 (133) |

**Supplementary Table 6.** Evaluation of assembly (Nonredundant and Noncontaminated Genome) completeness with respect to gene space using BUSCO.

| BUSCO | Number (Percentage) |
|---|---|
| Complete BUSCOs (C) | 1,579 (97.83%) |
| Complete and single-copy BUSCOs(S) | 1,310 (81.16%) |
| Complete and duplicated BUSCOs(D) | 269 (16.67%) |
| Fragmented BUSCOs(F) | 13 (0.81%) |
| Missing BUSCOs(M) | 22 (1.36%) |
| Total BUSCO groups searched | 1,614 (100%) |

**Supplementary Table 7.** Transposon elements (TE) annotation of *P. koreana* genome.

| **Classification** | **Count** | **Length(bp)** | **Percentage of genome (%)** |
|---|---|---|---|
| **LTR** | | | |
| Copia | 17,024 | 12,757,941 | 3.18 |
| Gypsy | 47,866 | 35,192,597 | 8.77 |
| unknown | 39,979 | 15,189,643 | 3.78 |
| **nonLTR** | | | |
| DIRS_YR | | | |
| LINE_element | 1,091 | 772,846 | 0.19 |
| unknown | 191 | 109,452 | 0.03 |
| **TIR** | | | |
| CACTA | 17,212 | 6,832,438 | 1.70 |
| Mutator | 21,654 | 7,879,959 | 1.96 |
| PIF_Harbinger | 8,225 | 2,464,697 | 0.61 |
| Tc1_Mariner | 1,849 | 852,149 | 0.21 |
| hAT | 12,179 | 4,521,301 | 1.13 |
| polinton | | | |
| **nonTIR** | | | |
| helitron | 121,956 | 49,275,078 | 12.28 |
| **repeat_region** | 47,380 | 13,429,450 | 3.35 |
| **Total** | 336,606 | 149,277,551 | 37.19 |

**Supplementary Table 8.** Statistics of the annotation of protein-coding genes of *P. koreana* genome.

| **Gene annotation statistics** | |
|---|---|
| Gene number | 37,072 |
| Average gene length (bp) | 3779.29 |
| Mean exon number per mRNA | 5.05 |
| Mean CDS number per mRNA | 4.92 |
| Average CDS length (bp) | 1136.04 |
| Average intron length (bp) | 1997.64 |
| Average single exon length | 318.16 |
| Average single CDS length | 230.77 |
| Average single intron length | 492.81 |

**Supplementary Table 9.** Functional annotation of protein-coding genes of *P. koreana* genome.

| **Dataset** | **Number(percentage)** |
|---|---|
| Pfam | 26,037(70.23%) |
| Interproscan | 33,307(89.84%) |
| KEGG | 11,253(30.35%) |
| NR | 34,300(92.52%) |
| Swiss-Prot | 26,971(72.75%) |
| KOG | 30,774(83.01%) |
| COG | 12,532(33.80%) |
| Tremble | 34,802(93.88%) |
| GO | 27,460(74.07%) |
| Unannotated | 1,692(4.56%) |
| Total | 37,072 |

**Supplementary Table 10.** Statistics of the annotated non-coding RNA.

| RNA type | | number |
|---|---|---|
| miRNA | | 3240 |
| tRNA | | 669 |
| SnRNA | | 91 |
| SnoRNA | | 516 |
| rRNA | 5S rRNA | 56 |
| | 5.8S rRNA | 6 |
| | 18S rRNA | 14 |
| | 28S rRNA | 14 |

**Supplementary Table 11(Excel).** Gene Ontology (GO) enrichment analysis of genes in expanded families of *P. koreana* genome.

**Supplementary Table 12 (Excel).** Geographical sampling information and summary statistics of whole-genome resequencing data for samples used in this study.

**Supplementary Table 13.** Environmental variables used in this study derive from WorldClim

| Code | Variable |
| --- | --- |
| Temperature-related | |
| BIO1 | Annual Mean Temperature |
| BIO2 | Mean Diurnal Range |
| BIO3 | Isothermality (BIO2/BIO7) |
| BIO4 | Temperature Seasonality |
| BIO5 | Maximum Temperature of Warmest Month |
| BIO6 | Minimum Temperature of Coldest Month |
| BIO7 | Temperature Annual Range (BIO5-BIO6) |
| BIO8 | Mean Temperature of Wettest Quarter |
| BIO9 | Mean Temperature of Driest Quarter |
| BIO10 | Mean Temperature of Warmest Quarter |
| BIO11 | Mean Temperature of Coldest Quarter |
| Precipitation-related | |
| BIO12 | Annual Precipitation |
| BIO13 | Precipitation of Wettest Month |
| BIO14 | Precipitation of Driest Month |
| BIO15 | Precipitation Seasonality |
| BIO16 | Precipitation of Wettest Quarter |
| BIO17 | Precipitation of Driest Quarter |
| BIO18 | Precipitation of Warmest Quarter |
| BIO19 | Precipitation of Coldest Quarter |

**Supplementary Table 14 (Excel)** Detailed information of environmental associated variants identified in this study.

**Supplementary Table 15** The number and proportion of the functional effects of the environmental-associated variants identified by LFMM, both LFMM and RDA relative to the whole genome level.

| | LFMM | LFMM&RDA | Genome |
|---|---|---|---|
| 3'UTR | 60(1.75%) | 28(1.57%) | 181,752(3.05%) |
| 5'UTR | 81(2.36%) | 55(3.09%) | 83,429(1.40%) |
| Upstream | 724(21.08%) | 284(15.96%) | 1,144,408(19.24%) |
| Downstream | 509(14.82%) | 278(15.63%) | 792,457(13.32%) |
| Nonsynonymous | 97(2.82%) | 56(3.15%) | 148,847(2.50%) |
| Synonymous | 64(1.86%) | 35(1.97%) | 126,672(2.13%) |
| Intron | 382(11.12%) | 187(10.51%) | 682,145(11.47%) |
| Intergenic | 1,501(43.70%) | 850(47.78%) | 2,767,838(46.52%) |
| Total | 3,438(99.51%) | 1,773(99.66%) | 5,927,548(99.63%) |

**Supplementary Table 16.** Summary of some functional important genes with relatively high numbers of environmental-associated variants detected.

| PK_GENE | Ptri_gene | Atha_gene | symbol | Chromosome | Associated variants | Related environment variables |
|---|---|---|---|---|---|---|
| Pokor00456 | Potri.001G412077 | AT4G27280 | CMI1 | LG01 | 14 | BIO15 |
| Pokor01235 | Potri.001G330900 | AT3G01910 | SOX | LG01 | 9 | BIO13,16,18 |
| Pokor01408 | Potri.001G312300 | AT1G17840 | ABCG11 | LG01 | 4 | BIO5 |
| Pokor02467 | Potri.001G205300 | AT5G09420 | TOC64-V | LG01 | 5 | BIO3 |
| Pokor09179 | Potri.005G052800 | AT4G14640 | CAM8 | LG03 | 3 | BIO2 |
| Pokor10559 | Potri.002G172800 | AT4G23980 | ARF9 | LG04 | 7 | BIO13,16,18 |
| Pokor10689 | Potri.002G159300 | AT2G45880 | BMY4 | LG04 | 17 | BIO12,13,14,16,17,18,19 |
| Pokor11081 | Potri.014G016300 | AT5G67500 | VDAC2 | LG04 | 2 | BIO12,13,14,16,17,18,19 |
| Pokor12242 | Potri.002G005000 | AT2G33620 | AHL10 | LG04 | 3 | BIO13,18 |
| Pokor12247 | Potri.002G004500 | AT2G33590 | CRL1 | LG04 | 31 | BIO12,13,14,16,17,18,19 |
| Pokor12248 | Potri.002G004100 | AT2G33590 | CRL1 | LG04 | 73 | BIO12,13,14,16,17,18,19 |
| Pokor13059 | Potri.004G085500 | AT5G37630 | EMB2656 | LG05 | 13 | BIO12,14,17,18,19 |
| Pokor13254 | Potri.004G103200 | AT5G39050 | PMAT1 | LG05 | 13 | BIO13 |
| Pokor14452 | Potri.004G230100 | AT3G18990 | VRN1 | LG05 | 38 | BIO14,17,19 |
| Pokor17228 | Potri.003G039475 | AT3G13860 | HSP60-3A | LG07 | 62 | BIO5,8,12,13,14,16,17,18,19 |
| Pokor18546 | Potri.003G168700 | AT5G14090 | LAZY1 | LG07 | 8 | BIO13,16,18 |
| Pokor18547 | Potri.003G168800 | AT1G27980 | DPL1 | LG07 | 12 | BIO13,16,18 |
| Pokor19962 | Potri.014G111400 | AT2G46970 | PIL1 | LG08 | 3 | BIO15 |
| Pokor21575 | Potri.011G151700 | AT3G15850 | FAD5 | LG09 | 82 | BIO5,8,10,12,13,14,16,17,18,19 |
| Pokor21577 | Potri.011G152200 | AT1G32900 | GBSS1 | LG09 | 20 | BIO8,10,12,13,14,16,17,18,19 |
| Pokor22025 | Potri.011G041700 | AT4G05070 | WIP2 | LG09 | 4 | BIO5 |
| Pokor23027 | Potri.008G079700 | AT5G16300 | COG1 | LG10 | 6 | BIO13 |
| Pokor25841 | Potri.007G138800 | AT3G60030 | SPL12 | LG12 | 87 | BIO12,13,14,16,17,18,19 |
| Pokor27800 | Potri.013G067500 | AT1G08810 | MYB60 | LG13 | 3 | BIO13 |
| Pokor28275 | Potri.013G120800 | AT4G35160 | ASMT1 | LG13 | 6 | BIO8,10,12,13,16,17,18,19 |
| Pokor29050 | Potri.018G051300 | AT5G07990 | TT7 | LG14 | 5 | BIO5 |
| Pokor29543 | Potri.018G101300 | AT2G23950 | CIK2 | LG14 | 20 | BIO1,2,4,6,7,9,11 |
| Pokor32219 | Potri.012G097300 | AT1G20960 | EMB1507 | LG16 | 27 | BIO5 |

**Supplementary Table 17 (Excel).** Risk of non-adaptedness (RONA) for environmental variables of BIO1-BIO19 across the 24 natural populations of *P. koreana* under eight different future climate scenarios.

**Supplementary Table 18**. Sequence of primers used for qRT-PCR test on abiotic stress

| Symbol | Forward Primer | Reverse Primer |
|---|---|---|
| HSP60-3A(Pokor17228) | CTAAGCCCAGATAGATACTTGC | ATTTCATAGGAATGGACCGAC |
| CRL1 (Pokor12447) | CGAGTTGTTTTGAAACCGAA | TAGGTGCTTATGAAAATACCAG |
| UBQ-10 | CCAAGCCCAAGAAGATCAAGC | GCACCGCACTCAGCATTAGG |
